# IRIS Integrates Sparse Sequence, Experimental, and AI-Predicted Structures for Protein–RNA Affinity Prediction and Motif Discovery

**DOI:** 10.1101/2025.09.10.675247

**Authors:** Eduardo Cisneros de la Rosa, Yafan Zhang, Xingcheng Lin

**Author notes:** E-mail: Xingcheng. Contributed equally to this work (Eduardo Cisneros de la Rosa, Yafan Zhang).

## Abstract

Protein–RNA interactions are fundamental to numerous cellular processes, yet quantitatively characterizing their binding specificity remains a major challenge. We present IRIS (Integrative RNA–protein interaction prediction Informed by Structure and sequence), a biophysical framework that integrates residue-level sequence and structural features without relying on large-scale affinity data to predict binding affinities and identify binding motifs. Applied across different protein–RNA systems, IRIS predicts relative binding free energies (ΔΔ*G*) with consistent correlations and competitive error metrics, and its performance is further improved by incorporating additional high-affinity sequences into the training set. By leveraging predicted structural complexes, IRIS reveals alternative binding modes not observed in experimental structures, extends applicability to systems lacking experimental protein–RNA complexes, and generates a library of favorable RNA-binding motifs at protein–RNA interfaces. Collectively, these results establish IRIS as a versatile framework that leverages increasingly accurate structural predictions to enable quantitative modeling and rational engineering of protein–RNA interactions.

## Introduction

Protein–RNA interactions govern diverse molecular and cellular processes, including post-transcriptional regulation,^1–3^ epigenetic control,^4,5^ and liquid–liquid phase separation of RNA condensates.^6–10^ Sequence- and structure-specific binding affinity quantitatively determines interaction strength and thereby governs specificity, regulatory dynamics, and downstream functional outcomes. Accurate prediction of binding affinities is therefore essential for understanding molecular mechanisms in health and disease.^11–14^

Experimental approaches for characterizing protein–RNA interactions have expanded substantially in both throughput and precision.^10,15,16^ High-throughput enrichment assays such as RNA Bind-n-Seq (RBNS),^17,18^ RNAcompete,^19^ and CLIP-based methods (eCLIP,^20^ iCLIP,^21,22^ PAR-CLIP^23^) can profile millions of sequences or in vivo binding footprints to reveal motifs and contextual preferences. Complementary platforms, including protein–RNA binding arrays,^24^ microfluidic systems (e.g., MITOMI-derived designs^25^ and the Riboreactor^26^), enable parallel, quantitative measurements of relative binding affinities across multiple targets with reduced sample consumption. Classical biophysical assays, including electrophoretic mobility shift assays (EMSA),^27^ surface plasmon resonance (SPR),^28^ isothermal titration calorimetry (ITC),^29^ fluorescence anisotropy, and microscale thermophoresis (MST)^30^ provide calibrated affinity and kinetic parameters. Each platform offers distinct strengths but also trade-offs: enrichment assays generally require external standards for absolute affinity calibration,^31^ CLIP-based methods can be biased by crosslinking chemistry and may not distinguish direct from indirect interactions^32–34^, and array/microfluidic systems require careful normalization (e.g., correcting for RNA/protein loading, background fluorescence, and reference scaling).^24,25^ Meanwhile, traditional biophysical assays provide precision but are inherently low throughput.

Computational models now complement experimental approaches by predicting binding specificity and affinity directly from sequence and structure. Sequence-based deep learning models such as DeepBind,^35^ ProBound,^31^ and DeepPNAP^36^ leverage large binding datasets, while transformer architectures trained on eCLIP-seq data (e.g., Reformer^37^) achieve single-nucleotide resolution. However, these methods are typically protein-specific, with their performance largely depending on the availability of training data for the target RNA-binding proteins (RBPs). When applied to novel RBPs, retraining on target-specific high-throughput binding affinity data is often required to achieve optimal results.

Nowadays, structure-based resources have expanded dramatically: RNAproDB^38^ catalogs over 3,500 experimentally determined protein–RNA complexes,^39^ and new structure-prediction frameworks, including AlphaFold3 (AF3),^40^ RosettaFoldNA,^41^ Chai-1,^42^ and Boltz,^43,44^ extend modeling to complexes beyond experimental reach. Deep learning methods such as PrismNet^45^ and HDRNet^46^ integrate sequential and structural features, but generally cast binding prediction as a binary classification problem, limiting their ability to yield quantitative affinities. Physical modeling approaches such as Rosetta–Vienna RNP-ΔΔG,^47^ FoldX,^48,49^ and PredPRBA^50^ can estimate relative affinities with reasonable accuracy, but require careful structural relaxation and are not well-suited for high-throughput applications. Despite these advances, a broadly applicable framework that simultaneously incorporates structural interpretability, quantitative affinity prediction, and generalizability to previously unseen RBPs, without requiring experimental binding affinity data for the specific target RBPs, remains lacking.

In this study, we present IRIS (Integrative RNA–protein interaction prediction Informed by Structure and sequence), a biophysical model integrating structural and sequence features from protein–RNA complexes, without relying on large-scale affinity data, to optimize physicochemical amino-acid-nucleotide interactions for predicting sequence-specific binding affinities. We first demonstrate the predictive accuracy of IRIS using the bacteriophage MS2 coat protein–RNA system, trained on experimentally determined protein–RNA complex structure. We then show that incorporating high-affinity sequence variants increases training sequence diversity and further enhances predictive performance. By leveraging AF3-modeled complexes,^40^ IRIS can identify alternative binding modes for RNAs with large sequence variation and enable affinity prediction in systems lacking experimental structures. In addition, we construct libraries of favorable RNA-binding motifs by combining structural interface information with systematic sequence variation. Finally, we demonstrate the general applicability of IRIS across multiple protein–RNA systems, including cases where only predicted structures are available.

## Materials and Methods

### IRIS Architecture: Residue-Level Interaction Energy Modeling

IRIS uses a residue-level biophysical framework that integrates detailed physicochemical modeling with structural data, either experimentally determined or predicted by structure-prediction tools,^40–43^ to quantify sequence-specific protein–RNA binding affinities (Figure 1). The model estimates relative binding free energies, ranks binding strengths, and converts these free energies into dissociation constants via the thermodynamic relation: ^51^

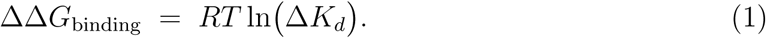

**Figure 1:**
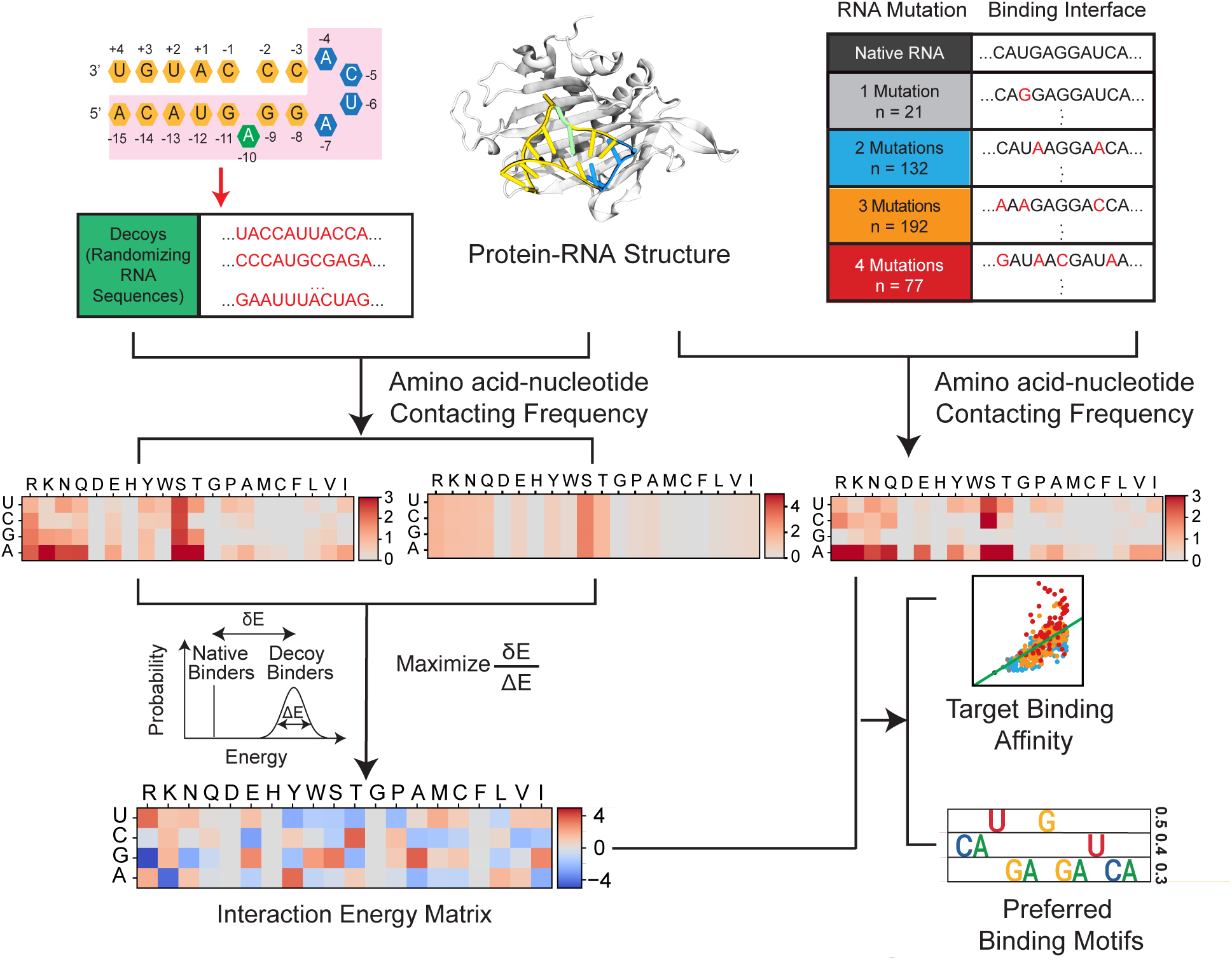
IRIS model architecture and evaluation using experimental protein– RNA structures. (A) Schematic overview of the IRIS pipeline, which integrates protein–RNA sequences and structures through a biophysical framework. Amino-acid–nucleotide contacting frequencies at the protein–RNA interface are used to optimize a residue-based energy matrix capturing favorable interactions. Target RNA sequences can replace (or be threaded into) the native RNA in the structure to generate a new contact frequency matrix, which is combined with the optimized energy matrix to predict sequence-specific binding affinities and identify preferred RNA-binding motifs. The bacteriophage MS2 protein–RNA complex (PDB ID: 2C4Q) is shown as an example, with nucleotides near the interface shaded in pink. These positions were used to generate 422 RNA sequences carrying single to quadruple substitutions for model evaluation.

IRIS infers residue-level protein–RNA binding free energies for each protein–RNA complex by optimizing a system-specific 20-by-4 interaction energy matrix *γ*(*a, n*), which quantifies the interaction strength between each type of amino acid *a* and nucleotide *n*. The model is trained using contact interactions extracted from the corresponding native complex structure, together with the associated native sequence and generated RNA decoy sequences.

This energy matrix is learned independently for each protein–RNA system by training on its structures and sequences. No experimental binding affinity data are used during model training. By integrating structural and sequence signatures of the protein–RNA binding interface, IRIS optimizes an interpretable energy model that enables accurate prediction of protein–RNA interactions.

#### Interface Definition and Contact Frequency Matrix Construction

Each complex is represented by a 20-by-4 amino-acid-nucleotide contact frequency matrix that captures pairwise contact frequencies between each amino acid (*a_i_*) and nucleotide (*n_j_*) at the protein–RNA interface, using C*α* (protein) to phosphorus atom (RNA backbone) distances, computed using a smooth hyperbolic tangent equation:

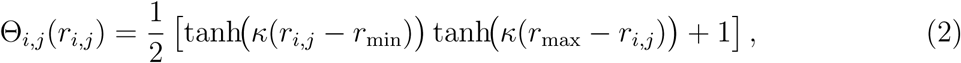

where *r*_max_ = *r*_min_ = 0.95 nm defines the outer contact boundary, and *κ* = 7.0 nm^−1^ controls the smoothness of the transition. This continuous weighting avoids discontinuities associated with hard cutoffs.

The amino-acid-nucleotide contact frequency matrix is computed as

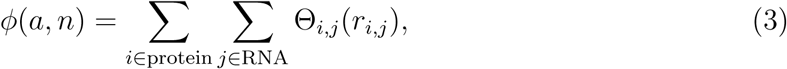

which yields a 20 × 4 matrix describing the total weighted contact frequencies for each residue-pair type.

#### Binding Energy Computation

The solvent-averaged binding free energy of the target sequence is defined as

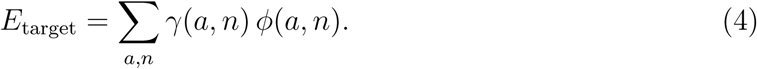

Here, *γ*(*a, n*) is a 20-by-4 energy matrix encoding residue-type specific interaction strength between amino acid (*a*) and nucleotide (*n*). By defining binding free energy in this manner, our model captures sequence-specific protein–RNA binding while considering their structural context. When training IRIS, *γ*(*a, n*) is optimized for each complex to maximize energetic separation between the native RNA sequence and an ensemble of RNA decoy sequences derived from that same complex.

The construction of RNA decoy ensembles is critical for defining the energetic landscape explored during training. As shown in Figure S1, decoy sequences are generated under controlled mutation distributions that either mimic experimentally observed binder/non-binder profiles or follow a uniform sampling across sequence space (see Section *Decoy Distribution Variants and Sequence Similarity Analysis* for details). These decoy sets span a range of Hamming distances and binding propensities, enabling the model to learn discriminative interaction patterns that distinguish high-affinity binders from weak or non-binding variants. This strategy ensures that the optimized interaction matrix *γ*(*a, n*) reflects both sequence composition and the statistical structure of biologically relevant mutation landscapes.

#### Energy Matrix Optimization

The model optimizes *γ*(*a, n*) energy matrix by maximizing the energy gap between the given strong binders and their corresponding decoy binders. This approach draws inspiration from methods used in protein folding, protein-protein, and protein-DNA interaction studies.^52–57^ In practice, IRIS computes the average energy gap between strong and decoy binders as *δE* = ⟨*E*_decoy_⟩ − ⟨*E*_strong_⟩, and the standard deviation of decoy binding energies as Δ*E* = Std(*E*_decoy_), where Std denotes the standard deviation.

Using Equation 3, define

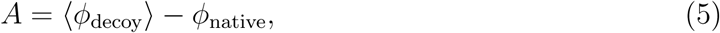

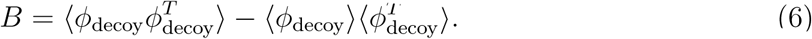

The normalized separation between strong and decoy binding energies becomes

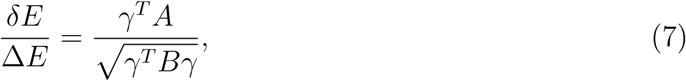

which is maximized when

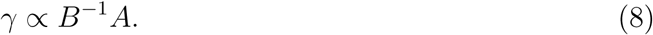

To ensure numerical stability, *B* is regularized via eigenvalue filtering: (1) diagonalizing *B* = *P* Λ*P* ^−1^, (2) retaining the top *N* eigenmodes, (3) replacing smaller eigenvalues with the *N* th largest value, and (4) reconstructing *B*^−1^ from these filtered components. The performance of the model with restricting different numbers *N* of eigenmodes is provided in Table S1. The optimized interaction energy matrix *γ* can be visualized in the 20×4 form (Figure 1), revealing the physicochemical preference at amino acid–nucleotide resolution.

#### Target Sequence Evaluation and ΔΔ*G* Calculation

To predict the binding affinity of a target protein–RNA sequence, we substitute it into the native high-affinity protein–RNA complex structure and recompute the corresponding *ϕ* contact frequency matrix using Equation 3. This matrix is then combined with the trained interaction energy *γ* matrix to calculate the binding free energy via Equation 4.

Since we focused our predictions on various RNA sequences binding to the same native RNA-binding protein, we hypothesized that all of them share the same conformational entropy and approximated the difference in binding free energy ΔΔ*G*_target_ to be the difference in computed solvent-average binding free energy *E*_target_ of different RNA sequences. Additionally, due to the presence of an undetermined scaling factor when optimizing the *γ* energy matrix, the predicted free energies are presented in reduced units. Despite this, the relative binding free energies ΔΔ*G*_target_ = *E*_target_ − *E*_native_ can be used to accurately rank binding affinities of target protein–RNA pairs, and the relative free energies can be converted to Δ*K_d_* using Equation 1. Here, *E*_native_ is the solvent-averaged binding free energy of the native sequence associated with the high-affinity training structure. Finally, the predicted binding free energies can be used to identify preferred RNA binding motifs.

In addition to the experimentally determined structures, predicted protein–RNA complex structures^40–43^ can also be used to compute *ϕ* matrices and the corresponding binding free energies using Eqs. 3 and 4. This enables evaluation of mutant sequences using either experimentally resolved or high-confidence predicted structures.

The constant conformational entropy hypothesis can be revised when considering plasticity in RNA structural ensembles,^58^ especially for RNAs with more drastic sequence mutations. As demonstrated in Figure S2, considering differences in predicted RNA secondary structures^59^ due to sequence variation can improve the predictive performance for sequences with large RNA mutations (see Supplementary Information Section *Incorporation of Predicted Secondary Structure in IRIS Prediction* for details).

#### Summary of Datasets in this Study

To systematically evaluate the model performance and benchmark against other state-of-the-art predictive methods, we collected three datasets (Table 1) based on data quality and structural availability:

1. Dataset 1 - MS2–RNA: High-quality binding affinity data measured by RNA-MaP,^24^ with an experimentally resolved crystal structure (PDB ID: 2C4Q).^60^ Mutations were restricted to positions within 0.5 nm of the protein–RNA interface. Because the crystal structure (PDB ID: 2C4Q) does not include the 5^′^ nucleotide, this nucleotide was excluded during IRIS evaluation to ensure consistency with the experimental structure, even though the 5^′^ nucleotide falls within the 0.5 nm mutation distance cutoff.
2. Dataset 2 - hnRNPK–RNA: A large affinity dataset from fluorescence binding assays,^61^ without available experimental complex structures. Predictions were performed using AF3-predicted structures across three sequence categories: 10 biological sequences, two-cytosine (two-C)-patch variants, and three-cytosine (three-C)-patch variants, following the classification in the experimental paper.^61^ Because we do not have an experimental structure, we did not filter the data based on the distance between the mutational site and proteins.
3. Dataset 3 - Four additional protein–RNA complexes obtained from the dataset reported by Kappel et al.^47^ This dataset has limited binding affinity data measured by earlier binding assays,^62–66^ spanning both RNA and protein mutations, including Fox-1–RNA, Pumilio homolog 1 (PUM1)–RNA, signal recognition particle (SRP)–RNA, and human U1 snRNP (U1A)–RNA and has experimental protein–RNA structures available (PDB IDs: 2ERR, 1M8W, 1HQ1, and 1URN).^62–64,66^ This dataset provides an additional benchmark for the model’s predictive performance, but should be considered with caution, as the data were collected from earlier experiments under different solution conditions, and include both protein and RNA mutations.

**Table 1:** Summary of datasets used in this study. The table lists the total number of RNA sequences (*n*_total_) in each dataset, along with their corresponding protein–RNA complexes and source of affinity data. Datasets are grouped into three categories: Dataset 1 (MS2– RNA), Dataset 2 (hnRNPK–RNA), and Dataset 3 (additional protein–RNA complexes).

| RNA Binding Protein | Dataset | $n_{\text{total}}$ | Structure | Source |
| --- | --- | --- | --- | --- |
| MS2 | 1 | 346 | PDB ID: 2C4Q | RNA-MaP Assay<sup>24</sup> |
| hnRNPK | 2 | 200 | Predicted Structure<sup>40</sup> | Fluorescence Binding Assay<sup>61</sup> |
| Fox-1 | 3 | 16 | PDB ID: 2ERR | Surface Plasmon Resonance Measurement<sup>64</sup> |
| PUM1 | 3 | 16 | PDB ID: 1M8W | EMSA<sup>65</sup> |
| SRP | 3 | 11 | PDB ID: 1HQ1 | Nitrocellulose Filter Binding Assay<sup>62</sup> |
| U1A | 3 | 17 | PDB ID: 1URN | Nitrocellulose Filter Binding Assay<sup>67</sup> |

#### Ablation Study

To evaluate model robustness, we focused on the most comprehensive and up-to-date MS2-RNA dataset (Dataset 1 in Table 1). Individual parameters were systematically varied while all others were held fixed. For each parameter configuration, the IRIS training procedure was repeated to generate a new *γ* energy matrix, which was then used to predict binding affinities. We also emphasize that IRIS is a biophysics-based, data-driven model, and many hyperparameters were chosen based on physical principles; accordingly, they were varied only within physically reasonable ranges. Details of the examined parameters are provided below, and the results of these independent ablation studies are summarized in Tables S1 and S2.

- **Decoy number**: The number of decoys was chosen to balance coverage of the protein– RNA interaction space with computational feasibility. We systematically varied the number of RNA decoys from 1k to 100k, and protein decoys from 0 to 50k.
- **Tanh function parameters**: A hyperbolic tangent function was used to quantify the contact frequencies between protein–RNA residues, using C*α* (protein) and phosphorus atom (RNA backbone). This function has two parameters: a contact distance cutoff *r*_max_ = *r*_min_ (with *r*_min_ being a dummy variable, since distances are positive) and a slope parameter *κ*. The default cutoff of 0.95 nm corresponds to the Debye-Hückel length at physiological salt concentration of 100 mM. We varied this cutoff from 0.75 nm (corresponding to 165 mM ions) to 1.15 nm (corresponding to 70 mM ions) to reflect a reasonable ionic screening length. The slope parameter *κ* controls the smoothness of the transition from non-contact to contact and follows coarse-grained protein-protein and protein-nucleic acid simulation potential.^68,69^ To test its robustness, we varied *κ* from 6.0 nm^−1^ to 8.0 nm^−1^.
- **Decoy distance cutoff** : This parameter defines the spatial region near the protein-RNA interface from which residues are selected for decoy sequence generation. We used the closest heavy atoms of each residue to calculate this distance, varying it from 0.95 nm to 1.5 nm.
- **Eigenmodes**: The selected number of eigen modes used in model optimization needs to strike a balance between making full use of the training data and avoiding utilizing unreliable eigenmodes with small eigenvalues.^54^ We varied the number of eigenmodes from 10 to 40. A finer grid search was performed for the case with 10k RNA decoys, where model performance was near optimal.
- **RNA decoy distribution**: We evaluated the model performance with decoy distribution generated to better match the experimental dataset. We implemented two strategies (Figure S1): (1) matching the nucleotide distribution at each residue position; (2) matching the Hamming distance distribution relative to the wild-type sequence. We used two reference datasets to generate decoys: one consisting of sequences with experimentally measured binding affinities, called “Binders”; and another comprising weak binders with *K_d_ >* 2,000 nM (“Non-Binders”), which fall outside the reliable experimental measurement range. Further details are provided in Section *Decoy Distribution Variants and Sequence Similarity Analysis*.
- **Mutation distance cutoff** : This parameter defines the filtering criterion applied during benchmarking against experimental mutational affinity data. Variants are retained if the mutated residue (protein or RNA) lies within a specified distance of the protein–RNA interface in the native training structure. Distance is defined as the minimum heavy-atom separation between the mutated residue and any residue across the interface. Smaller cutoffs restrict evaluation to mutations proximal to the binding interface. This filtering is applied only to the evaluation dataset and does not affect model training.

#### Model Generalization and Performance Metrics

The IRIS model is trained independently for each protein–RNA complex to obtain a system-specific *γ* energy matrix. As a result, an energy matrix trained on one complex cannot be directly applied to predict binding affinities for RNA sequences interacting with a different RNA-binding protein (Figure S3). The advantage of this target-specific training strategy is its ability to accurately model sequence-dependent binding affinities by directly learning the interaction energetics of a given protein–RNA interface. Thus, the transferability of IRIS lies in its training protocol rather than in a universally applicable energy matrix.

With the emergence of deep-learning-based protein–RNA structure prediction tools, ^40–44^ it has become increasingly feasible to generate training structures for optimizing target-specific energy matrices, provided that at least one strong-binding RNA sequence is known. In particular, our results on the hnRNPK–RNA system (Figure 7; Tables 3, S2, and S3) demonstrate the utility of incorporating deep-learning-predicted structures into model training, provided that high-confidence protein–RNA interfaces are used. Therefore, while a trained energy matrix is not transferable across different RNA-binding proteins, the IRIS framework can be readily retrained to predict RNA-binding affinities for new protein targets. To evaluate predictive performance, we report Pearson correlation (*r*), Spearman correlation (*ρ*), area under the Receiver Operating Characteristic (ROC) curve (AUC), area under the precision-recall curve (PR-AUC), and error metrics including mean absolute error (MAE) and root-mean-square error (RMSE). Pearson (*r*) and Spearman (*ρ*) correlations were computed directly between predicted and experimental affinity values. AUC and PR-AUC scores evaluate binary classification performance for distinguishing weak (W) and strong (S) binders, defined within each complex based on the top and bottom quartiles of experimental ΔΔ*G* values, respectively. MAE and RMSE were computed relative to a regression line constrained to pass through the origin and are reported with bootstrap 95% confidence intervals. We note that MAE and RMSE should be interpreted cautiously when comparing across models due to differences in output scaling.

### Correlation and Linear Regression Analysis of Binding Free Energies

To assess predictive accuracy, we compared IRIS-predicted and experimentally measured relative binding free energies (ΔΔ*G*) for MS2 protein binding across mutated RNA sequences.

Sequences were grouped into two categories: those with 0–3 substitutions and those with four substitutions. Pearson’s *r* and Spearman’s *ρ* were calculated to quantify linear agreement and rank consistency, respectively. For the four-substitution group, correlations were computed using only quadruple mutants, with high-affinity outliers retained in the analysis but omitted from the *Sub 4* legend for clarity. This grouping reflects two distinct regimes of sequence variation: variants with 0–3 substitutions generally preserve the native protein–RNA binding interface assumed during model construction, whereas variants with four substitutions or more frequently introduce structural deviations or alternative binding modes.

Two types of linear regressions were performed for visualization:

- **Through-origin regression:** A linear regression constrained through the origin (intercept = 0) was fitted to the 0–3 substitution group:

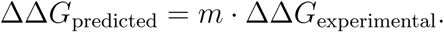

This line was used as a common reference across both the 0–3 and 4-substitution panels, enabling direct visual comparison (Figures 2A, 3C).

- **Free-intercept regression:** Free-intercept linear regressions were also computed for each complex and condition to evaluate potential systematic offsets. These plots are included in Figure S5.

**Figure 2:**
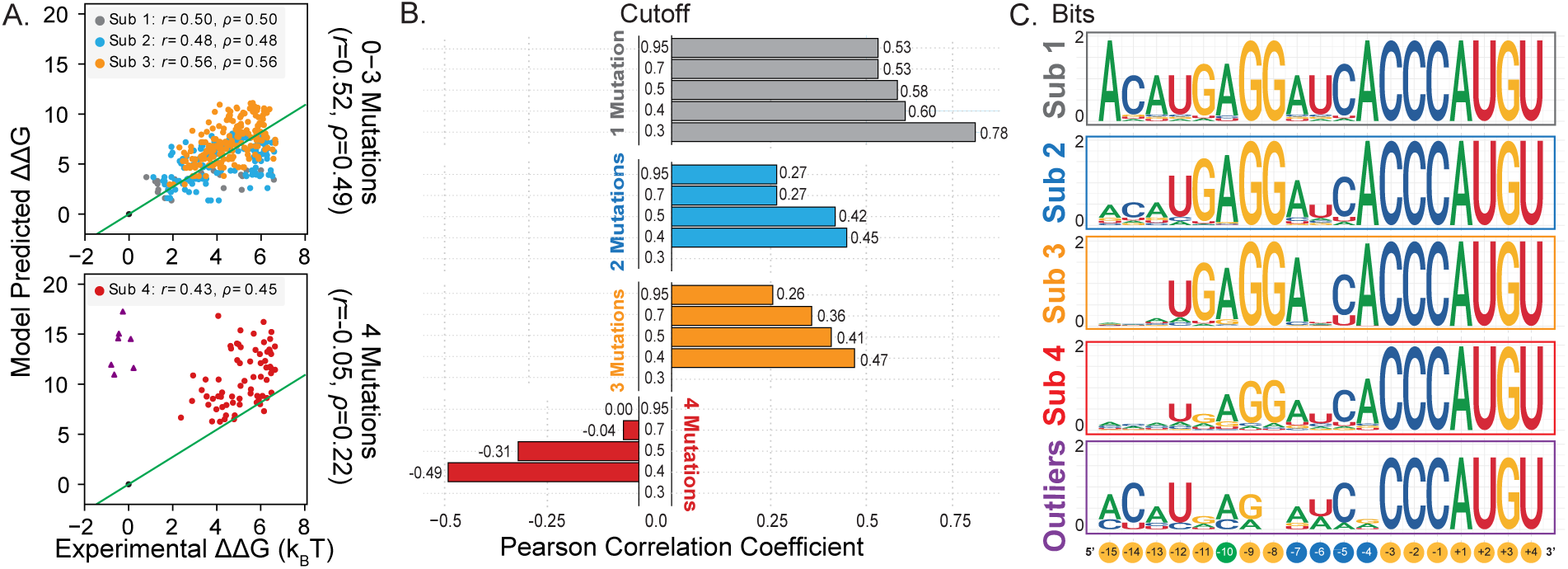
Predicted binding affinities of the MS2 protein–RNA complex for single to quadruple RNA mutations. (A) Comparison of predicted binding affinities (converted to binding free energies) with experimental measurements.^24^ The best-fit regression line was computed using 0–3 substitutions and applied to the quadruple mutants for reference. High-affinity outliers are highlighted with purple triangles. Correlation values (Pearson’s *r* and Spearman’s *ρ*) include all sequences, while the Sub4 legend omits outliers for clarity. (B) Predicted binding affinities across different protein–RNA interface distance cutoffs (nm), trained on the crystal structure (PDB ID: 2C4Q) and grouped by the total number of RNA mutations, including mutations outside the protein–RNA interface (pink region in Figure 1). (C) Sequence logos showing nucleotide preferences by substitution number. Sequences with 1–3 substitutions largely retain native patterns, whereas outliers display distinct sequence divergence.

All plots used standardized axis limits and consistent color and shape encodings, facilitating visual comparison across groups. This design highlights both the model’s overall predictive strength and its limitations when applied to highly divergent sequences.

### Decoy Distribution Variants and Sequence Similarity Analysis

To evaluate whether the choice of decoy distribution influences IRIS predictive performance, we systematically varied the RNA decoy generation strategy and retrained the model to predict RNA binding affinities.

In the default setting, RNA decoys were generated from the native sequence by introducing random nucleotide substitutions while preserving sequence length. Mutations were sampled uniformly across positions and nucleotide identities. Alternatively, decoys can be constructed to probe the influence of sequence composition and experimental bias, including base-composition–matched decoys and distributions enriched for experimentally observed strong or weak binders.

To examine the effect of decoy distribution, we focus on the MS2–RNA dataset,^24^ which includes experimentally measured binders (*K_d_ <* 2, 000 nM) and non-binders (*K_d_ >* 2, 000 nM). Based on these data, we implemented two strategies: (1) matching the nucleotide distribution at each residue position and (2) matching the Hamming distance distribution relative to the wild-type sequence. The resulting sequence distributions and the predicted binding free energies for these decoys are presented in Figure S1. For each decoy distribution, the full IRIS training procedure was independently repeated, yielding a new energy matrix *γ* for predicting binding affinities.

### Binding Interface Cutoff Modulates Predictive Accuracy

To evaluate how the definition of the binding interface influences predictive performance, we calculated Pearson correlations between predicted and experimental binding free energies across mutation distance cutoffs ranging from 0.3 to 0.95 nm. This analysis was performed on five protein–RNA complexes with experimentally resolved structures: 2C4Q, 2ERR, 1IM8, 1HQ1, and 1URN (Figures 2B and 5B). For each complex, predicted binding free energies were computed at five mutation distance cutoffs, and correlations with experimental values were determined. For MS2 (2C4Q), results were further stratified by the number of nucleotide substitutions (1–4) at each cutoff. The data were visualized as faceted bar plots, including a zero-correlation reference line to highlight thresholds where predictive accuracy diminishes or reverses. This analysis underscores the model’s sensitivity to interface definitions and helps identify optimal distance thresholds for accurate binding affinity prediction. More detailed calculations using other evaluation metrics are provided in Tables S5-S10.

### Bit Score Calculation and Sequence Logo Analysis

To assess nucleotide conservation and identify enriched RNA motifs, we calculated perposition information content using standard information-theoretic measures. At each alignment position *i*, the frequency of each nucleotide *f_i_*(*b*) was computed across all sequences, and the information content in bits is calculated as:

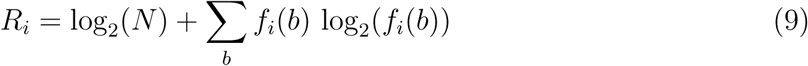

where *N* = 4 represents the four RNA nucleotides (A, C, G, U). Higher *R_i_* values indicate greater conservation at a given position.

Sequence logos (Figure 2C) were generated by stacking letters at each position, scaled according to the computed information content. The height of each stack corresponds to total conservation, while the relative height of each nucleotide reflects its frequency at that position. Columns with low information content appear short or empty, even if multiple nucleotides are present, highlighting positions with high sequence variability.

For practical implementation, sequences were grouped by mutation class (e.g., single, double, triple, and quadruple substitutions), and sequence logos were generated using the ggseqlogo package in R^70^ with the method="bits" setting. This approach allows direct visualization of per-position conservation and nucleotide preference, facilitating the identification of motifs that contribute most strongly to protein–RNA binding.

### Binding Interface Motifs Across Distance Cutoffs

To examine how the definition of the binding interface affects conserved RNA-binding motifs, we analyzed the base composition of interface residues at different distance cutoffs (0.3, 0.4, and 0.5 nm) using sequence logo representations. This analysis was performed on the MS2 protein–RNA complex, focusing on sequences with 1 and 4 substitutions (Figure 3D), as well as on test sequences from four additional protein–RNA complexes: 2ERR, IM8W, 1HQ1, and 1URN (Figure 5A).

**Figure 3:**
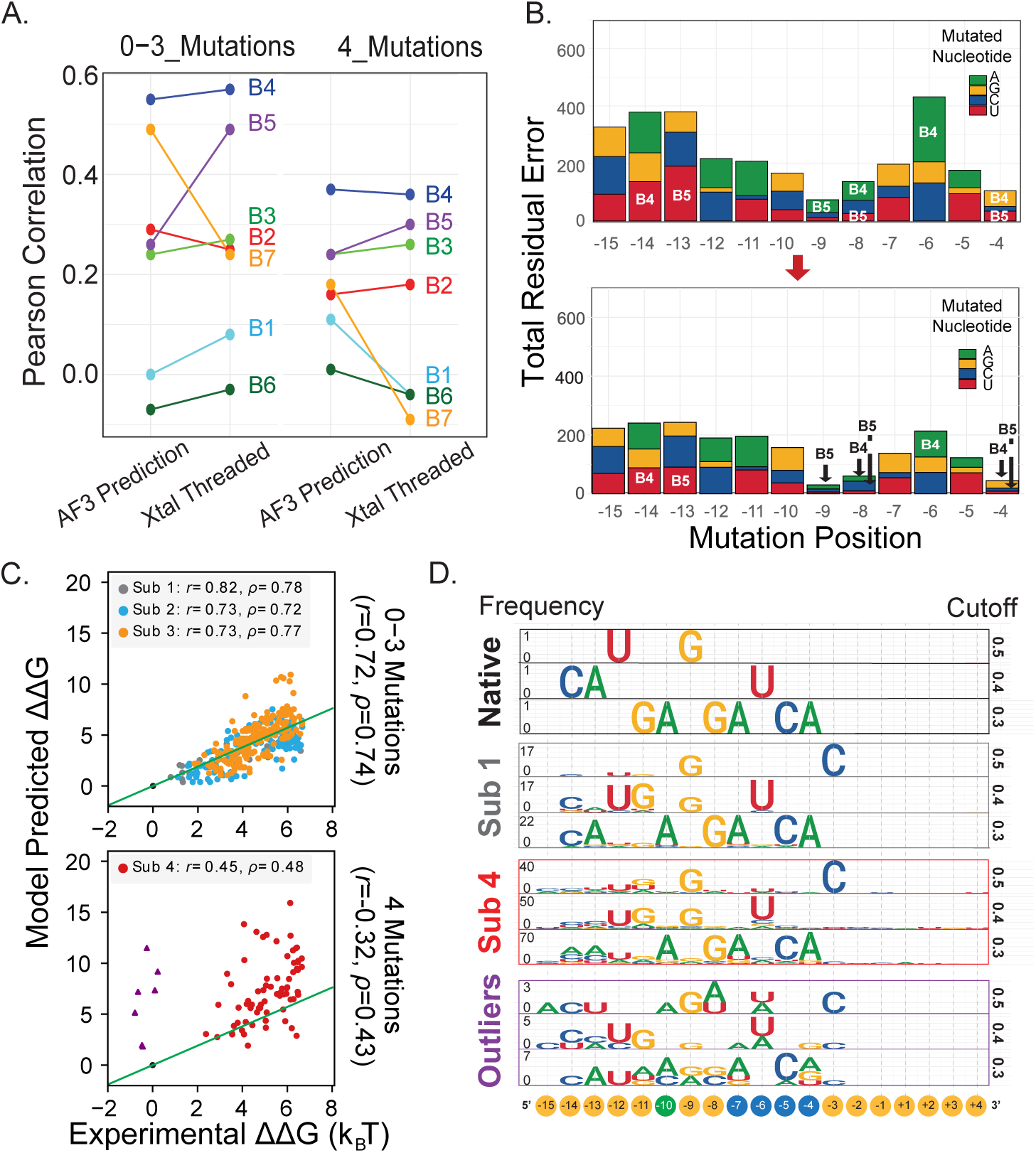
IRIS predictions reveal how training-set diversity improves protein– RNA binding accuracy. (A) Comparison of IRIS-predicted binding affinities using either AF3-predicted structures or B1-B7 outlier sequence threaded into the native crystal structure (PDB ID: 2C4Q). AF3 predictions improve accuracy for B7, the strongest binder, suggesting an alternative binding mode not captured by the crystal structure. (B) Total Residual Error at each mutated nucleotide position, shown as a stacked bar plot colored by nucleotide identity. High-affinity B4 and B5 sequences contribute disproportionately to total error, highlighting positions missed by the original model. (C) Correlation between predicted and experimental binding free energies after retraining the model with the native 2C4Q structure and sequence, plus the threaded sequences of B4 and B5. Both Pearson’s *r* and Spearman’s *ρ* are shown; the Sub4 legend excludes outliers for clarity, whereas the reported correlations include them. (D) Structure-based sequence logo from AF3-predicted structures of the native, Sub1, Sub4, and high-affinity outlier sequences. Positions are stratified by distance to the protein interface. Native and Sub1 share similar interfaces, Sub4 retains partial conservation, and high-affinity outliers show distinct binding patterns, reflecting structural and sequence divergence. This logo provides a “binding fingerprint” to guide motif identification and model training.

For each cutoff, RNA nucleotide identities were extracted using residue-level annotations and mapped to sequence positions. To isolate the contributions of each distance threshold, residues were assigned hierarchically: residues within 0.3 nm were included in the closest-contact group, residues between 0.3–0.4 nm were included only if not already assigned to the 0.3 nm group, and residues between 0.4–0.5 nm were included only if not assigned to the smaller cutoffs. This procedure ensures that each cutoff captures distinct layers of the protein–RNA interface without redundancy.

Nucleotide frequencies (A, C, G, U) were then computed for each group to generate position-specific frequency matrices. Stacked sequence logos were constructed using the ggseqlogo package in R,^70^ enabling direct comparison of nucleotide enrichment patterns across distance-defined interface layers.

These visualizations reveal how sequence preferences vary with proximity to the protein, distinguishing core interface contacts from more peripheral interactions. This analysis provides insight into which distance thresholds capture meaningful RNA-binding motifs and informs the identification of high-affinity binding features (Figure 6).

### Total Residual Error Across Mutation Positions and Nucleotide Identities

To investigate how specific nucleotide substitutions impact the predictive accuracy of MS2 coat protein–RNA interactions, we computed the *Total Residual Error* stratified by both mutation position and nucleotide identity. All mutational variants and the wild-type RNA sequences used in Figure 1 were obtained and processed as described in Section *Target Sequence Evaluation and* ΔΔ*G Calculation*, where binding free energies were predicted by substituting the native RNA sequence and computing ΔΔ*G* values from the *ϕ* and *γ* matrices. We leveraged the linear regression model described in Section *Correlation and Linear Regression Analysis of Binding Free Energies*, which was fitted through the origin on sequences containing three or fewer substitutions (Substitution ≤ 3). This regression of predicted versus experimentally measured ΔΔ*G* values provides the expected predicted binding free energy for a given experimental measurement under near-native mutational load, with the best-fit line shown in Figures 2A and 3C.

For each sequence, the *residual error* was calculated as the absolute difference between the predicted binding energy and the value projected by the regression model:

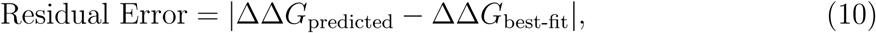

where ΔΔ*G*_best-fit_ = *m* · ΔΔ*G*_experimental_ and *m* is the slope of the fitted linear regression.

Residual errors were then aggregated across all sequences, grouped by mutation position and substituted nucleotide (A, C, G, or U). This yields a total residual error for each nucleotide at each position. The results are visualized as a stacked bar plot (Figure 3B), with bars colored by nucleotide identity and a fixed *y*-axis for consistent scaling. This representation highlights positions and substitutions that disproportionately contribute to prediction errors, revealing model blind spots and guiding potential refinement.

Additional residual errors were also computed for the other four protein–RNA complexes, using origin-constrained regression (Figure S6) and free-intercept regression (Figure S7). Further details are provided in Supplementary Information Section *Parameter Calibration and Error Analysis of IRIS Predictions*.

### Jaccard Similarity Heatmap of RNA Sequences Against Known MS2 Motifs

To compare sequence similarity between input RNA sequences and known RNA-binding motifs, we generated a Jaccard similarity heatmap using a k-mer–based approach. For each pairwise comparison between a test RNA sequence and a motif, the Jaccard similarity was computed using all unique k-mers of length 3 (*k* = 3), defined mathematically as:

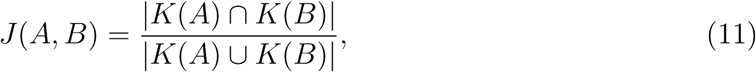

where *K*(*A*) and *K*(*B*) are the sets of k-mers in sequences *A* and *B*, respectively.

To compare similarity among MS2-binding RNA sequences, we selected 10 RNA sequences reported by Buenrostro et al.,^24^ including the native sequence in the crystal structure (PDB ID: 2C4Q), seven strong-binding quadruple-mutation RNA sequences (B1–B7), and two additional RNA mutants (R1 and R2) that were not included in Dataset 1 due to their high Hamming distance (*>* 4) from the native sequence, or the presence of mutations outside the binding interface (Figure 1, pink shaded region). Experimentally identified MS2-RNA binding motifs from prior CLIP-seq and CryoEM studies^71–73^ (Motifs I–XIV) were also included. Each sequence was converted into a set of 3-mers, and pairwise Jaccard similarities were calculated between the two groups.

The resulting similarity matrix was visualized as a heatmap, with experimentally identified motifs on the y-axis, and RNA sequences from Buenrostro et al.^24^ on the x-axis (Figure 4A). This k-mer–level analysis enabled rapid comparison of short RNA sequences and facilitated identification of the motifs most closely matched by each test sequence. Because this approach is alignment-free, it is robust to minor shifts or indels, making it particularly well-suited for comparing short, structured RNAs.

### K-mer Graph Construction and Clustering of High-Affinity RNA Fragments

To identify and visualize structure-informed patterns of RNA fragments associated with high protein-binding affinity, we extracted contiguous stretches of residue IDs and their corresponding nucleotide identities from all the AF3-predicted RNA-binding structures. Blocks of protein-interacting RNA residues were defined under varying interaction distance cutoffs, and for each residue, we recorded the smallest cutoff at which it remained in contact with the protein.

Contiguous segments longer than four residues were further divided into overlapping sub-k-mers (e.g., 4-mers, 5-mers). Each k-mer was annotated with its sequence, length, per-position cutoff values, and the experimentally measured ΔΔ*G*_exp_ (relative binding energies with respect to the native wild-type sequences) of its associated RNA sequence. To standardize comparisons across the dataset, ΔΔ*G*_exp_ values were converted into Z-scores:

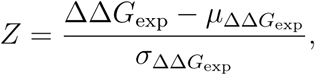

where *µ*_ΔΔ*G*exp_ and *σ*_ΔΔ*G*exp_ denote the mean and standard deviation of all ΔΔ*G*_exp_ values for each protein–RNA complex, computed across the full dataset reported for that complex prior to k-mer segmentation, thereby avoiding bias introduced by fragment-level counting.

After collecting all k-mers, Z-scores were aggregated for each unique sequence. We then computed a pairwise string similarity matrix using the Levenshtein distance via the stringdist R package.^74^ This matrix served as input for agglomerative hierarchical clustering with average linkage (implemented using hclust in R). The resulting dendrogram (Figure 6) highlights relationships among k-mers based on sequence similarity, revealing distinct sequence families. This approach uncovers recurring structural and sequence features of RNA that strongly bind proteins and provides guidance for designing high-affinity RNA sequences.

### AF3-Predicted HnRNPK–B Motif Structures and IRIS Affinity Prediction

In the absence of experimentally determined hnRNPK-B motif complexes, we used AF3^40^ to generate predicted structures for IRIS model training and binding-affinity prediction. The hnRNPK protein sequence was obtained from Uniprot^75^ (https://www.uniprot.org/ uniprotkb/P61978/entry#sequences), and the B motif RNA GCAGCCCCAGCCCCAGC-CCCUACCCCUGCCCCUGCCCCUGC was selected for its stable secondary structure and cytosine-rich patches. ^61^ We performed twenty independent AF3 runs using different random seeds and observed modest overall confidence (highest-confident model: ipTM = 0.5 for the complex; average pLDDT scores of 66.22 for protein, and 43.05 for the RNA). Particularly, lower confidence was observed in the intrinsically disordered RG/RGG region (average pLDDT score = 40.71) and the KH3–RNA binding interface (average pLDDT scores = 81.99 for KH3, and 46.77 for the contacting RNA region), indicating increased uncertainty in these regions (Figure 7).

To focus on regions with more confident protein–RNA structural predictions, we trained the IRIS model using the KH1+KH2-B motif interface from the top-ranked predicted structure. The resulting *γ* energy matrix was applied to three prediction tasks: binding affinities for ten biologically relevant RNAs (10bio), for B-motif variants containing two cytosine patches (two-C patches), and for variants containing three cytosine patches (three-C patches).

When target RNAs differ in length from the training sequence (e.g., biologically relevant RNAs), we adopted a sliding-window approach based on the training sequence. Specifically, when the target sequence is longer than the training sequence, we segmented the target sequence into a series of 41-bp segments from the 5^′^ to 3^′^ end in 1-nucleotide increments, and calculated the binding free energy for each target segment. If the target sequence is shorter than the training sequence, we scan the target sequence to replace a portion of the training sequence, incrementing one nucleotide at a time, thereby generating a set of new target sequences for binding free energy evaluation. For example, the 43 bp Sirloin motif yields 43 − 41 + 1 = 3 overlapping 41 bp fragments, and the shorter 36 bp Ucp2 sequence yields 41 − 36 + 1 = 6 fragments within the same 41 bp framework. IRIS predicts a binding free energy for each fragment, and we report the lowest (i.e., strongest binding) value among them as the final predicted ΔΔ*G* for the full-length target RNA sequence. This approach ensures that IRIS identifies the highest-affinity sequence segment for each target.

We also evaluated the model trained on the lower-confidence full-length hnRNPK-B-RNA complex and the KH1+KH2+RG/RGG-RNA complex, as well as subsets of protein-RNA interface filtered using progressively higher confidence thresholds. The results are presented in Figures S8-S10 and Tables S2-S3.

## Results

### Sequence-Specific Protein–RNA Affinity Predicted Using Structure Optimization

IRIS is a residue-based, data-driven protein–RNA model that uses an optimized residue-based energy matrix to predict sequence-specific binding affinities of protein–RNA systems. IRIS builds on our previous IDEA framework, originally developed for predicting protein– DNA binding affinity,^57^ and extends it to RNA. As illustrated in Figure 1, IRIS integrates sequence and structural information from protein–RNA complexes to optimize a 20-by-4 interaction energy matrix that encodes residue-type–specific interaction strengths between amino acids and nucleotides. Given target protein–RNA sequences, we substitute them into the training protein–RNA complex structure to calculate residue contact matrices, which count the frequencies of amino acid–nucleotide contacts at the binding interface. These matrices are then combined with the trained energy matrix to compute target binding free energies. The training structure can be derived from experimentally determined structures^76^ or from confidently predicted structures.^40–43^ More details are provided in Section *IRIS*

### Architecture: Residue-Level Interaction Energy Modeling

To evaluate IRIS in predicting sequence-specific protein–RNA binding affinities, we first trained the model on protein–RNA complexes with available experimental structures (dataset details are provided in the *Materials and Methods* Section, *Summary of Datasets in this Study*). We selected the bacteriophage MS2 coat protein, for which both high-resolution crystal structures and high-throughput experimental binding data are available.^24,60^ IRIS was trained on a single protein–RNA complex (PDB ID: 2C4Q^60^) and used to predict binding affinities, subsequently converted to binding free energies using Equation 1 for 422 RNA variants carrying mutations near the protein–RNA interface (Figure 1, pink shaded region). To assess how spatial proximity influences prediction quality, we filtered these variants based on their distance to protein C*_α_* atoms, defining a *mutation distance cutoff* (see *Materials and Methods* Section, *Ablation Study*), and systematically evaluated how varying this cutoff affects prediction accuracy across different mutation regimes.

Our results reveal that both the number of mutations and the proximity within the protein–RNA interface strongly affect IRIS’ predictive performance. As shown in Figure 2A, IRIS correctly predicts the sequence-specific binding affinities at smaller (single to triple) mutations, but fails for RNA sequences with quadruple mutations. In addition, reducing the mutation distance cutoff from 0.95 nm to 0.3 nm increases the correlation between experimental and predicted binding free energies (Figure 2B), indicating that focusing IRIS on residues closer to the binding interface improves its ability to capture binding specificity. However, decreasing the cutoff does not rescue the predictions for quadruple-mutation data, with smaller cutoff values showing more negative correlations. Our initial choice of protein–RNA binding interface (pink shaded region in Figure 1) aligns most closely with the 0.5 nm cutoff, with the addition of the missing 5^′^ terminal nucleotide at position -15 that is absent from the 2C4Q structure. This definition strikes a balance between data size and predictive quality and is therefore used in the following discussion.

The negative correlation observed in the four-mutation cases was primarily due to a failure in predicting the binding affinities of seven strong-binding RNA sequences (Figure 2A, bottom, purple triangles). To examine whether sequence conservation contributes to this effect, we grouped RNAs by the number of mutated nucleotides, aligned the sequences, and analyzed information content at each position in bits (Figure 2C, see *Materials and Methods* Section, *Bit Score Calculation and Sequence Logo Analysis*). Single to triple mutants showed strong sequence conservation at specific positions, while quadruple mutants excluding outliers (Sub4) retained weaker but still detectable conservation. In contrast, the quadruple-mutation outliers exhibited strongly divergent sequence features, particularly between positions −15 and −9, yet maintained high binding affinity. This suggests that the training set contains too few high-affinity sequences with motif patterns distinct from the dominant training examples. Including such sequences would provide broader variation at key binding positions and help the model better learn how sequence differences influence binding affinity.

### Expanding Training Sequence Diversity Improves Predictive Accuracy

To evaluate whether increasing sequence variation in the training set improves prediction accuracy, we retrained IRIS by incorporating experimentally identified strong-binding sequences. We focused on the seven strong-binding quadruple-mutation variants (hereafter “B1” to “B7”) that were poorly predicted by the native single-structure-sequence training scheme (Figure 3A). Using the Total Residual Error metric stratified by mutation position and nucleotide identity, we quantified the absolute differences between experimental ΔΔ*G* values and predictions from a best-fit linear model trained on sequences with three or fewer substitutions (Figure 3B) (see Methods, *Total Residual Error Across Mutation Positions and Nucleotide Identities*).

Among these variants, B4 and B5 contain substitutions at positions associated with the largest residual errors and show consistent or improved predictions when threaded onto the native crystal structure (Figure 3A), identifying them as strong candidates for augmenting the training dataset. Sequence diversity analysis further supports this selection: Jaccard similarity calculation and principal component analysis indicate that B4 and B5 exhibit the lowest sequence similarity relative to the native sequence (Figures S11 A, B). Moreover, unlike B7, they preserve the native protein–RNA structural interface (see *Results*, Section *Alternative Protein–RNA Binding Interfaces Revealed by Structural Prediction and Motif Analysis*). Incorporating B4 and B5 into training substantially improved model performance, reducing accumulated residual errors (Figure 3B), increasing the Pearson correlation coefficient above 0.73 for single-to triple-mutants, and restoring a positive correlation of 0.32 for quadruple mutants (Figure 3C). In addition, retraining yields a more consistent relationship between sequence similarity and experimental binding affinities compared to the original single-structure model (see Supplementary Information, Sections *Effect of Sequence Similarity on IRIS Predictions* and *Selection of Training Sequences for Augmenting IRIS Prediction*).

Together, these results demonstrate that augmenting the training set with sequences that expand sequence diversity and target high-error regions enhances IRIS generalization and predictive accuracy across diverse variants.

### Integrating Deep-Learning-Based Structure Predictions to Expand Training Data

Recent advances in deep-learning-based structural prediction^40–44^ provide an opportunity to further expand the training data when experimental structures are unavailable. Building on our analysis of sequence diversity, we next tested whether incorporating predicted protein– RNA complex structures could improve IRIS performance. Specifically, we used AF3^40^ to predict structures for B1 to B7, selected the highest-confidence models based on the average predicted Local Distance Difference Test (pLDDT) score, retrained IRIS with these structures, and then evaluated performance by computing the Pearson correlation (*r*) and Spearman correlation (*ρ*) between experimental and predicted ΔΔ*G* values (Figure 3A).

For most binders, threading the sequences through the crystal structure (PDB ID: 2C4Q) slightly outperformed the use of AF3-predicted structures (Figure 3A). Notably, however, AF3 predictions provided a substantial improvement for B7, which has the highest experimentally measured binding affinity. This suggests that B7 may adopt an alternative binding mode not captured by the crystal structure.

To further examine the structural variation introduced by AF3, we constructed a structure-based sequence preference logo at the protein–RNA interface. Each sequence was modeled with AF3, and sequence logos were generated by stratifying nucleotide positions according to their distance from the protein (Figure 3D) (see *Materials and Methods*, Section *Binding Interface Motifs Across Distance Cutoffs*). This approach yields a structural “fingerprint” of sequence preferences across the binding interface, highlighting how predicted structures can complement experimental data in refining model training.

### Alternative Protein–RNA Binding Interfaces Revealed by Structural Prediction and Motif Analysis

Our analysis of the strong-binding outlier B7 suggested that it may engage the protein through an alternative binding site, as captured by the AF3-predicted structure (Figure 3A). To explore this further, we compared the predicted B7 structure with the native crystal structure (PDB ID: 2C4Q) using VMD.^77^ The predicted B7 RNA adopts a distinct binding interface compared to the native sequence (Figure 4B), which may explain the poor predictions obtained when threading B7 onto the crystal structure (Figure 3A). Consistent with this observation, aligned protein–RNA contact maps (Figure S12) show that B1–B6 largely preserve the native interaction pattern, whereas B7 exhibits pronounced differences in contact distributions, confirming a shift to an alternative binding interface.

**Figure 4:**
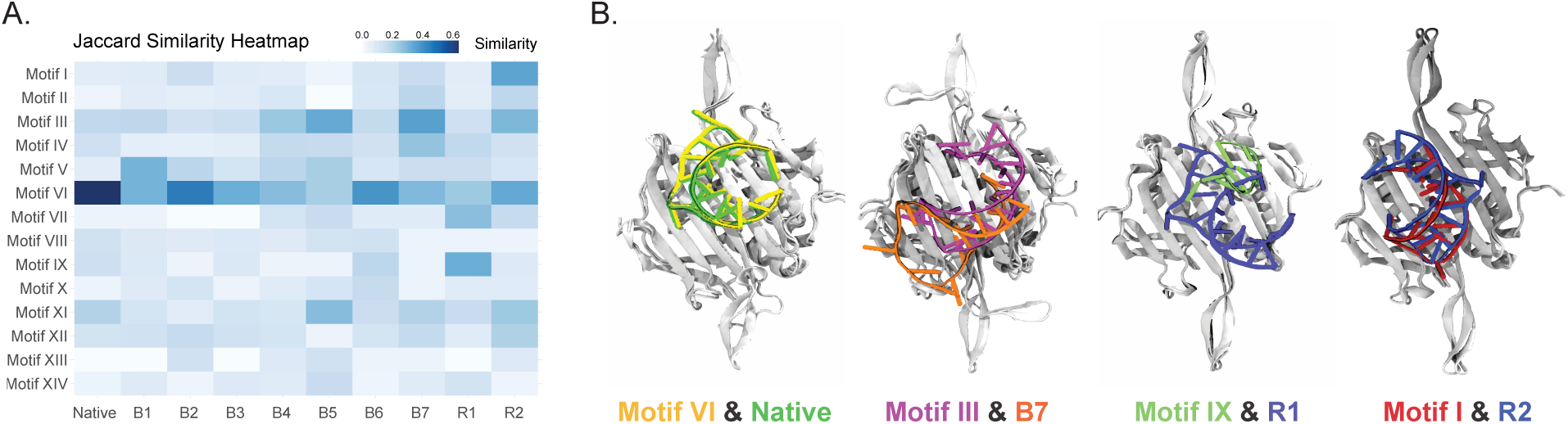
Alternative RNA binding modes revealed by structural predictions and motif similarity. (A) Jaccard similarity heatmap comparing strong-binding RNA sequences (Native, B1–B7, R1, and R2) with 14 experimentally identified MS2 motifs (Motifs I–XIV).^71–73^ The native sequence aligns most closely with Motif VI, B7 with Motif III, R1 with Motif IX, and R2 with Motif I, indicating distinct sequence signatures. (B) Structural comparison of AF3-predicted RNA sequences (B7, R1, R2) with the native MS2 crystal structure (PDB ID: 2C4Q), highlighting how sequence-dependent structural variations generate distinct binding interfaces that may influence model prediction accuracy.

The structural flexibility of RNA^58^ allows small sequence variations to produce distinct conformations and, in turn, alternative binding modes with RNA-binding proteins. To confirm that the B7 binding mode is not an artifact of structural prediction, and to assess whether alternative binding sites exist, we compared B1–B7 sequences against experimentally identified MS2 RNA binding motifs^71–73^ using Jaccard similarity (Figure 4A, see Methods, *Jaccard Similarity Heatmap of RNA Sequences Against Known MS2 Motifs*). This metric captures overlap in interaction-relevant sequence features and aligns with how IRIS evaluates sequence similarity (see Supplementary Information, Section *Effect of Sequence Similarity on IRIS Predictions*). The native sequence shows the strongest similarity to Motif VI, whereas B7 aligns most closely with Motif III. Consistently, AF3-predicted structures of Motif VI and Motif III closely resemble the native crystal structure and the B7-predicted structure, respectively.

Expanding this analysis to all fourteen known MS2 motifs, we identified two additional candidates with distinct binding interfaces: Motif IX and Motif I (Figure 4B). In agreement, two sequences from the RNA-MaP^24^ dataset with the highest sequence similarity, R1 (ACAUGAAGAGCACUCAUGU) and R2 (ACUUGAGUAACAACCAUGU) (Figure 4A), differ only modestly from the native sequence (ACAUGAGGAUCACCCAUGU) yet adopt distinct binding modes. These results further reinforce the idea that sequence perturbations can shift RNA binding toward alternative structural configurations not captured by a single-template model.

Together, these findings support the presence of multiple protein–RNA binding modes that can arise from minor sequence changes. This structural diversity likely explains the reduced predictive accuracy observed in certain quadruple mutants and strong-binding outliers, as these RNAs may engage the protein through conformations not represented by the native structure. Incorporating such alternative binding modes into the training framework would likely improve IRIS predictions by better capturing the accessible interaction landscape and reducing systematic bias against structurally divergent, high-affinity sequences.

### Generalizability of IRIS Predictions Across Diverse Protein–RNA Complexes

To assess the broader applicability of IRIS beyond the MS2–RNA system, we next evaluated its predictive performance on four additional protein–RNA complexes with available binding affinity data (Dataset 3 in Table 1; PDB IDs: 2ERR,^64^ 1M8W,^66^ 1HQ1,^62^ and 1URN^47,63^). For each complex, we calculated Pearson correlation values across a range of mutation distance cutoffs and examined how mutations in either the RNA or protein influenced the nucleotides included within the cutoff-defined binding interface.

Consistent with our observations for MS2, predictive accuracy improves as the mutation distance cutoff decreases across all four complexes (Figure 5A). Notably, 2ERR and 1URN required more stringent cutoffs to achieve strong correlations, though this reduced the number of retained sequences due to a more restricted interface definition (Tables S4, S6-S9). For 1URN, all mutations occur in the protein rather than the RNA, which may explain the model’s reduced performance, as IRIS is trained using only RNA decoys. To assess whether incorporating protein variation improves predictions, we introduced protein decoys (Table S5). Across RNA–protein complexes, adding protein decoys sometimes improved regression and classification metrics (e.g., RMSE, MAE, ROC-AUC, and PR-AUC) depending on the decoy level; however, the improvements were not consistent across systems or decoy settings, and no single configuration generalized robustly.

To further examine how sequence and structural divergence shape predictions, we generated structure-based sequence logos for the four complexes using AF3-predicted structures (Figure 5B). Similar to our findings in the MS2 complex (Figure 3D), some interface regions remained conserved across RNA variants, whereas others showed either the introduction of alternative nucleotides (e.g., position 8 in 1HQ1) or positional shifts of the same nucleotide within the interface (e.g., position 5 in 1M8W). These results highlight that both nucleotide identity and spatial positioning relative to the protein interface are critical determinants of binding specificity and affinity across diverse protein–RNA systems.

**Figure 5:**
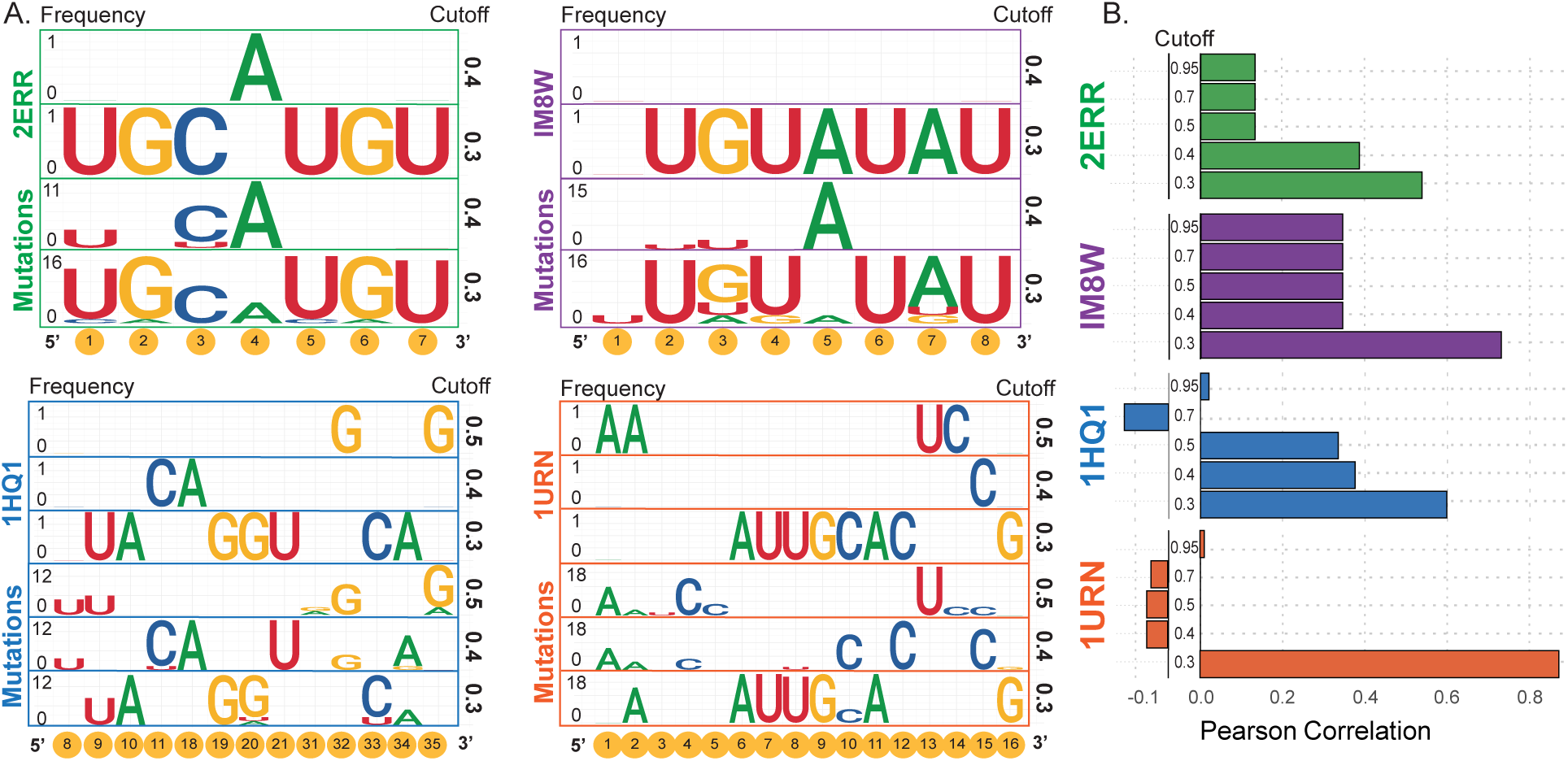
IRIS captures conserved and variable binding patterns across multiple protein–RNA complexes. (A) Structure-based sequence logos derived from AF3-predicted structures of four protein–RNA complexes (PDB IDs: 2ERR, 1IM8, 1HQ1, and 1URN), showing conserved and variable regions within the RNA binding interfaces. Some nucleotide positions remain consistent across mutations, while others exhibit substitutions or positional shifts, reflecting structural determinants of binding specificity. (B) Pearson correlations between IRIS-predicted and experimentally measured binding free energies across the four complexes, evaluated over different mutation distance cutoffs. Stricter interface definitions generally improve predictive accuracy, whereas lower correlations in complexes such as 1URN reflect protein-only mutations outside IRIS’s RNA-focused training data.

### Conserved K-mers Reveal Shared Binding Features Across Protein– RNA Complexes

The structure-based sequence logos serve as a “fingerprint” of RNA-binding proteins, highlighting conserved patterns in strong-binding RNA sequences. To investigate whether these patterns are shared across different protein–RNA complexes, we collected high-affinity RNA binding motifs, or “k-mers,” from AF3-predicted structures of selected mutated RNA sequences. For 2C4Q, we included the sub-1 and sub-4 mutants along with the strong-binding outliers, while for the other complexes (2ERR, 1M8W, 1HQ1, 1URN), we included all mutated sequences corresponding to their crystal structures. K-mers were extracted from nucleotides located within 0.5 nm of the protein interface, based on the minimum separation between the heavy atoms of the protein and RNA residues.

To assess similarity among these k-mers, we computed pairwise Levenshtein distances between all sequences, reflecting the minimal number of nucleotide edits required to convert one k-mer into another. Hierarchical clustering was then applied using these distance metrics, grouping k-mers with high sequence similarity and generating a dendrogram to visualize the relationships between clusters. Average relative binding free energies from normalized experimental ΔΔ*G*_exp_ values were calculated for each cluster to evaluate binding strength patterns (Figure 6; see *Materials and Methods*, *K-mer Network Construction and Clustering of High-Affinity RNA Fragments*).

**Figure 6:**
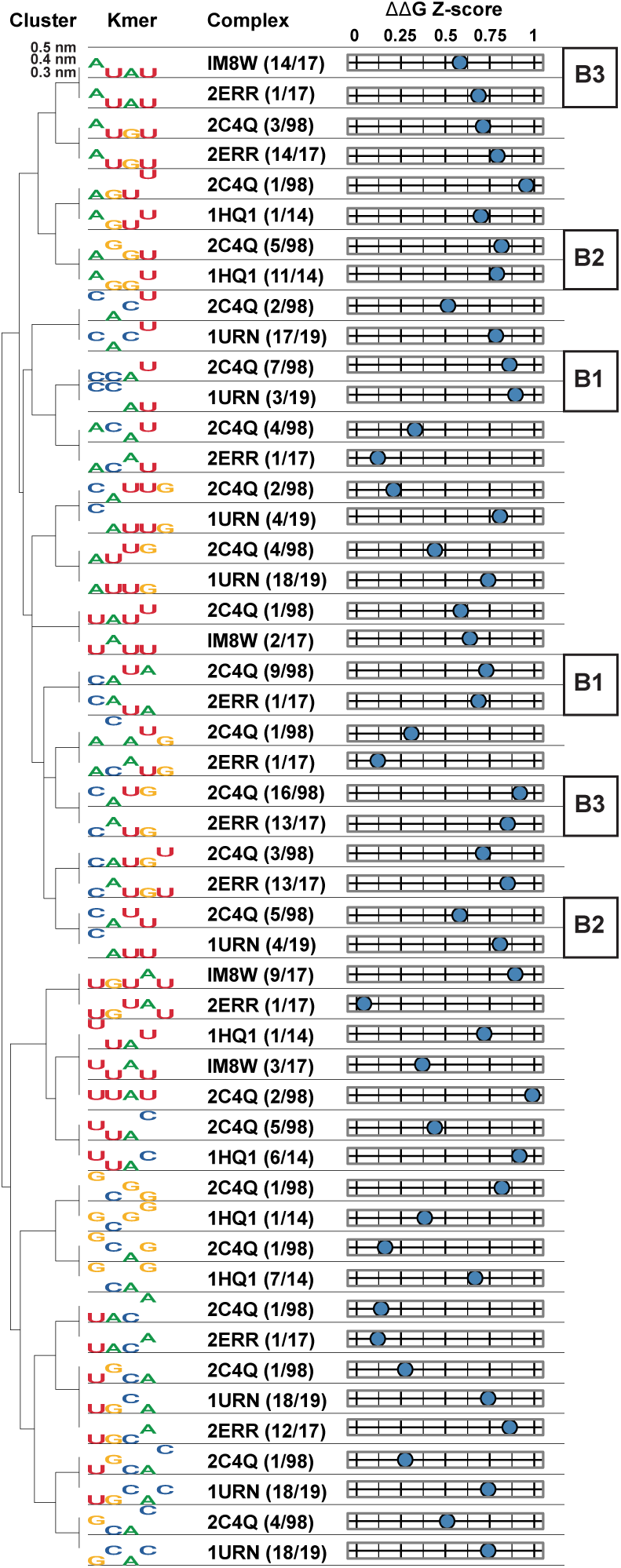
Hierarchical clustering of high-affinity RNA k-mers identifies favorable RNA binding motifs at the protein–RNA interface. High-affinity protein-interacting RNA fragments were extracted from AF3-predicted structures of the binding sequences and segmented into overlapping k-mers (length ≥ 4). Each k-mer was annotated with sequence identity, distance to the protein surface (0.3–0.5 nm), and experimental binding free energy (ΔΔ*G*_exp_), which was converted into normalized Z-scores. Pairwise Levenshtein distances were computed between k-mers to quantify sequence similarity, and hierarchical clustering was applied to group similar k-mers into motif families. The resulting dendrogram reveals clusters of RNA fragments with high protein-binding potential, providing a framework for motif discovery and aptamer design.

This analysis produced a library of shared structure-based sequence logos across the protein–RNA complexes examined, where certain k-mers, and sometimes their reverse sequences, displayed similar predicted binding affinities. Notably, some previously identified MS2 outlier binders (B1 to B3) shared k-mers with the strong-binding clusters in this library, confirming their high-affinity nature. As experimental binding data expand across diverse RNA-binding proteins, this approach may enable the compilation of a comprehensive library of conserved RNA-binding motifs, complementing motifs identified from high-throughput experiments,^17,78–80^ and facilitating studies of conserved RNA-protein binding patterns and their functional consequences.^81,82^

### Incorporating Predicted Structures Extends IRIS Predictability to Cases Lacking Experimental Structures

Our previous analyses (Figures 3 and 4) highlight how deep-learning-based structure predictors can reveal novel protein–RNA binding sites for the target protein. To evaluate their broader applicability for predicting binding affinities in the absence of experimental structures, we applied IRIS to heterogeneous nuclear ribonucleoprotein K (hnRNPK), a prototypical RNA-binding protein essential for gene regulation^83–86^ (Dataset 2 in Table 1). HnRNPK contains four positively charged RNA-binding domains, including three KH (K homology) domains connected by a 100-residue arginine-glycine (RG/RGG) linker ^61,87^ (Figure 7A). Experimentally determined binding affinities are available for these domains, with fluorescence anisotropy assays measuring interactions between full-length hnRNPK and ten biologically relevant RNA sequences, as well as cytosine-patch variants.^61^

**Figure 7:**
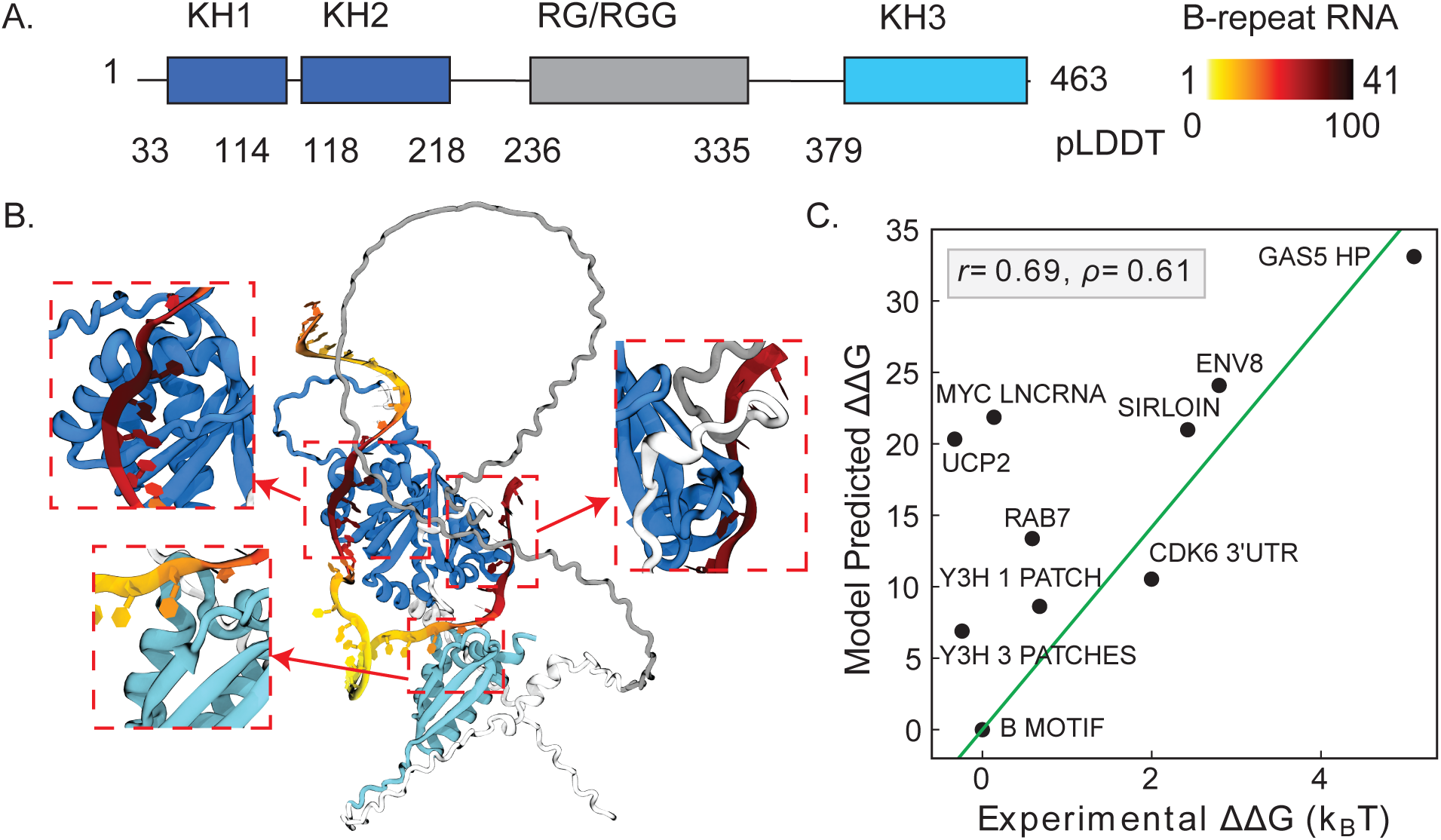
AF3-predicted structure of the hnRNPK–B-motif complex and IRIS-predicted binding affinities for biological RNA sequences. (A) Domain architecture of hnRNPK, showing KH1, KH2, RG/RGG, and KH3 regions in different colors. The bound B-motif RNA is colored according to the pLDDT score (yellow to red, low to high confidence) predicted by AF3.^40^ (B) AF3-predicted structure of hnRNPK bound to the B-motif RNA, with protein and RNA domains colored as in panel A. Three protein–RNA interaction interfaces are highlighted and shown in zoomed-in views (red dashed boxes). (C) Predicted binding affinity (ΔΔ*G*) from IRIS trained on the high-confidence KH1+KH2–B-motif binding interfaces, compared with experimental ΔΔ*G* values for 10 biological RNA sequences. Pearson and Spearman correlation coefficients are shown in the legend.

Using AF3,^40^ we predicted the complex structure of hnRNPK bound to the wild-type B-motif RNA, which contains cytosine-rich patches and interacts with hnRNPK with high affinity to form a stable complex.^61,88^ The top-ranked prediction (The interface predicted template modeling (ipTM) score = 0.5) indicated heterogeneous confidence levels across different regions. Specifically, structural predictions assigned low confidence to the disordered RG/RGG domain and the KH3–RNA interface (Figure 7B) (see *Materials and Method*, *AF3-Predicted HnRNPK–B Motif Structures and IRIS Affinity Prediction* for details), in agreement with experiments showing that the RG/RGG domain is intrinsically disordered^89^ and that the KH3 domain is dispensable for RNA binding.^61^ To focus on the most reliable structural interface, we trained the energy matrix on the KH1/2-RNA interface and used it to predict binding affinities for 10 biological RNA sequences. The prediction correlates strongly with experimental measurements (Figure 7C, Pearson correlation coefficient 0.69, Spearman’s rank correlation coefficient 0.61). Including the RG/RGG-RNA binding interface (Figure S8A) or both the RG/RGG- and KH3-RNA interfaces (Figure S8B) in training slightly reduces predictive accuracy, underscoring that higher-confidence structure prediction leads to more accurate binding predictions.

We further evaluated the model on two additional datasets: the two-cytosine-patch and three-cytosine-patch variants of the B-repeat RNAs. Consistent with the results for biological sequences, training on the KH1/2–RNA structural interface yielded the strongest correlation with experimental affinities (Figures S9A and S10A), whereas including the RG/RGG–RNA interface (Figures S9B and S10B) or both the RG/RGG– and KH3–RNA interfaces (Figures S9C and S10C) reduced predictive accuracy. The model showed lower performance on the three-cytosine-patch variants, likely because their narrower range of experimental affinities demands greater sensitivity to discriminate among RNA binders.

We further assessed whether AF3 confidence could be used to identify more reliable interfaces for IRIS training. Using increasing pLDDT thresholds, we found that the unfiltered full interface (pLDDT = 0) already yielded reasonable predictive performance, while higher thresholds improved accuracy, with the best threshold-based results obtained at pLDDT ≥ 60 (Table S2). However, the KH1/KH2–RNA interface alone still produced the highest overall accuracy (Figure 7C; Table S3), indicating that AF3 confidence is useful for filtering reliable structural interfaces, but that biologically informed domain selection further improves IRIS performance. Together, these results suggest that higher-confidence AF3 interfaces are generally associated with improved IRIS predictions, although determining a universal confidence threshold will require validation across larger protein–RNA datasets.

### Benchmarking Performance Against Sequence- and Structure-Based Models

To systematically evaluate IRIS, we benchmarked it against representative sequence-based, structure-based, and hybrid models, including DeepBind,^35^ Reformer,^37^ Rosetta–Vienna RNP-ΔΔG,^47^ DeePNAP,^36^ and FoldX.^49^ Implementation details are provided in the Supplementary Information, Section *Benchmarking IRIS Predictive Performance with Other Models*.

Across all datasets (Table 1), IRIS achieves consistently comparable or better performance (Tables 2, 3, 4) without training on experimental binding affinity data. For Dataset 1 (MS2–RNA, Table 2), which includes recent high-quality high-throughput affinity data and an experimental structure, the augmented IRIS model (trained on B4, B5, and the native sequence) performs best across all metrics, while the native-only IRIS model ranks second. For Dataset 2 (hnRNPK-RNA, Table 3), which contains extensive affinity measurements but no experimental structures, IRIS trained on the AF3-predicted complex performs comparably to DeepBind,^35^ ranking among the top two methods for biological sequences and two-cytosine variants (Figures S8 and S9), and achieving the best performance for three-cytosine variants (Figure S10). This demonstrates IRIS’s ability to predict sequence-specific protein–RNA binding affinities in the absence of experimental structures using high-quality predicted structures. While sequence-based models such as DeepBind remain effective in this setting, other structure-based methods fail to capture hnRNPK-RNA binding specificity, suggesting limited compatibility with predicted structures.

**Table 2:**
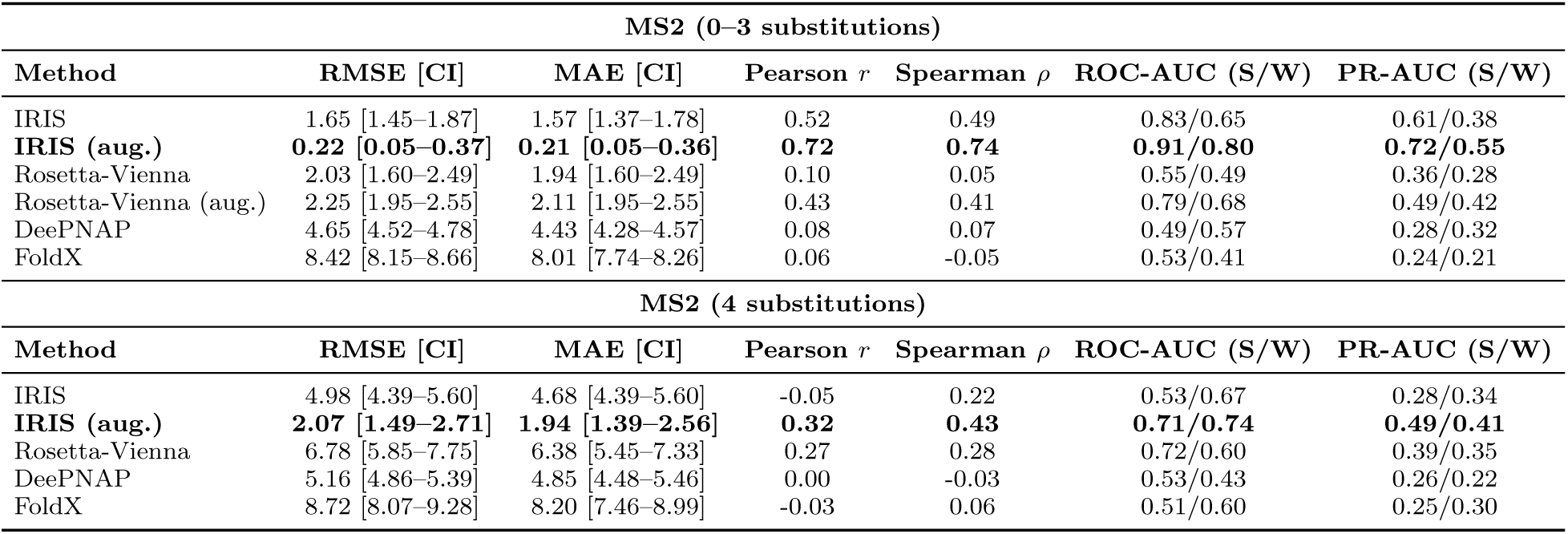
Prediction error, correlation, and classification performance on MS2 datasets (0–3 substitutions and 4 substitutions). Reported metrics were computed by comparing predicted and experimental ΔΔ*G* values. Mean absolute error (MAE) and root-mean-square error (RMSE) were calculated relative to the best-fit line through the origin, with 95% confidence intervals obtained via bootstrap resampling. Pearson (*r*) and Spearman (*ρ*) correlations were computed between predicted and experimental values. Strong (S) and weak (W) binders were defined as the lowest and highest experimental ΔΔ*G* quartiles, respectively. Classification performance was evaluated using ROC-AUC and PR-AUC, based on quartile-based binarization of predicted ΔΔ*G*. Rows in **bold** indicate the best-performing method for each group. IRIS results were obtained using the default parameters highlighted in bold in Table S1 (0 protein decoys, 10,000 RNA decoys). Rosetta-Vienna RNP-ΔΔ*G*^47^ predictions are labeled as “Rosetta-Vienna”. Reformer^37^ and DeepBind^35^ were not evaluated due to the lack of pre-trained models for the MS2-RNA system.

| MS2 (0–3 substitutions) |  |  |  |  |  |  |
| --- | --- | --- | --- | --- | --- | --- |
| Method | RMSE [CI] | MAE [CI] | Pearson $r$ | Spearman $\rho$ | ROC-AUC (S/W) | PR-AUC (S/W) |
| IRIS | 1.65 [1.45–1.87] | 1.57 [1.37–1.78] | 0.52 | 0.49 | 0.83/0.65 | 0.61/0.38 |
| **IRIS (aug.)** | **0.22 [0.05–0.37]** | **0.21 [0.05–0.36]** | **0.72** | **0.74** | **0.91/0.80** | **0.72/0.55** |
| Rosetta-Vienna | 2.03 [1.60–2.49] | 1.94 [1.60–2.49] | 0.10 | 0.05 | 0.55/0.49 | 0.36/0.28 |
| Rosetta-Vienna (aug.) | 2.25 [1.95–2.55] | 2.11 [1.95–2.55] | 0.43 | 0.41 | 0.79/0.68 | 0.49/0.42 |
| DeePNAP | 4.65 [4.52–4.78] | 4.43 [4.28–4.57] | 0.08 | 0.07 | 0.49/0.57 | 0.28/0.32 |
| FoldX | 8.42 [8.15–8.66] | 8.01 [7.74–8.26] | 0.06 | -0.05 | 0.53/0.41 | 0.24/0.21 |
| MS2 (4 substitutions) |  |  |  |  |  |  |
| Method | RMSE [CI] | MAE [CI] | Pearson $r$ | Spearman $\rho$ | ROC-AUC (S/W) | PR-AUC (S/W) |
| IRIS | 4.98 [4.39–5.60] | 4.68 [4.39–5.60] | -0.05 | 0.22 | 0.53/0.67 | 0.28/0.34 |
| **IRIS (aug.)** | **2.07 [1.49–2.71]** | **1.94 [1.39–2.56]** | **0.32** | **0.43** | **0.71/0.74** | **0.49/0.41** |
| Rosetta-Vienna | 6.78 [5.85–7.75] | 6.38 [5.45–7.33] | 0.27 | 0.28 | 0.72/0.60 | 0.39/0.35 |
| DeePNAP | 5.16 [4.86–5.39] | 4.85 [4.48–5.46] | 0.00 | -0.03 | 0.53/0.43 | 0.26/0.22 |
| FoldX | 8.72 [8.07–9.28] | 8.20 [7.46–8.99] | -0.03 | 0.06 | 0.51/0.60 | 0.25/0.30 |

**Table 3:**
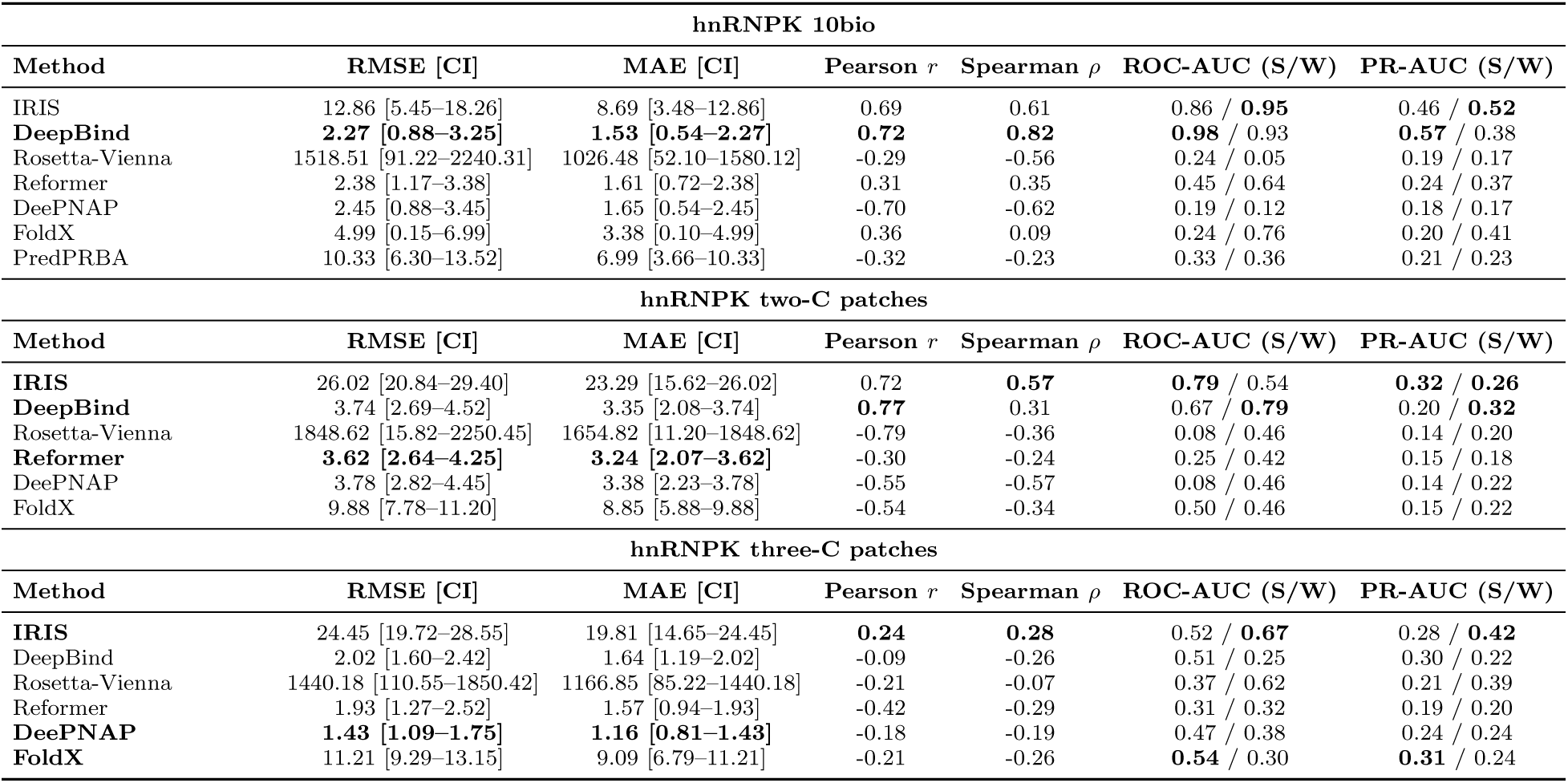
Benchmark comparison of representative methods on three hnRNPK datasets: 10 biological (10bio), two-cytosine (two-C) patch, and three-cytosine (three-C) patch variants. Mean absolute error (MAE) and root-mean-square error (RMSE) were calculated relative to the best-fit line through the origin, with 95% confidence intervals obtained via bootstrap resampling. Pearson (*r*) and Spearman (*ρ*) correlations were computed directly between predicted and experimental values. Strong (S) and weak (W) binders were defined as the lowest and highest experimental ΔΔ*G* quartiles, respectively. Classification performance was evaluated using ROC-AUC and PR-AUC, based on quartile-based binarization of predicted ΔΔ*G*. Rows in **bold** indicate the best-performing method for each group. PredPRBA^50^ results for the hnRNPK 10bio dataset are included for reference. Additional details on these models are provided in the Supplementary Information, Section *Benchmarking IRIS Predictive Performance with Other Tools*.

| hnRNPK 10bio |  |  |  |  |  |  |
| --- | --- | --- | --- | --- | --- | --- |
| Method | RMSE [CI] | MAE [CI] | Pearson $r$ | Spearman $\rho$ | ROC-AUC (S/W) | PR-AUC (S/W) |
| IRIS | 12.86 [5.45–18.26] | 8.69 [3.48–12.86] | 0.69 | 0.61 | 0.86 / **0.95** | 0.46 / **0.52** |
| **DeepBind** | **2.27 [0.88–3.25]** | **1.53 [0.54–2.27]** | **0.72** | **0.82** | **0.98** / 0.93 | **0.57** / 0.38 |
| Rosetta-Vienna | 1518.51 [91.22–2240.31] | 1026.48 [52.10–1580.12] | -0.29 | -0.56 | 0.24 / 0.05 | 0.19 / 0.17 |
| Reformer | 2.38 [1.17–3.38] | 1.61 [0.72–2.38] | 0.31 | 0.35 | 0.45 / 0.64 | 0.24 / 0.37 |
| DeePNAP | 2.45 [0.88–3.45] | 1.65 [0.54–2.45] | -0.70 | -0.62 | 0.19 / 0.12 | 0.18 / 0.17 |
| FoldX | 4.99 [0.15–6.99] | 3.38 [0.10–4.99] | 0.36 | 0.09 | 0.24 / 0.76 | 0.20 / 0.41 |
| PredPRBA | 10.33 [6.30–13.52] | 6.99 [3.66–10.33] | -0.32 | -0.23 | 0.33 / 0.36 | 0.21 / 0.23 |
| hnRNPK two-C patches |  |  |  |  |  |  |
| Method | RMSE [CI] | MAE [CI] | Pearson $r$ | Spearman $\rho$ | ROC-AUC (S/W) | PR-AUC (S/W) |
| **IRIS** | 26.02 [20.84–29.40] | 23.29 [15.62–26.02] | 0.72 | **0.57** | **0.79** / 0.54 | **0.32** / **0.26** |
| **DeepBind** | 3.74 [2.69–4.52] | 3.35 [2.08–3.74] | **0.77** | 0.31 | 0.67 / **0.79** | 0.20 / **0.32** |
| Rosetta-Vienna | 1848.62 [15.82–2250.45] | 1654.82 [11.20–1848.62] | -0.79 | -0.36 | 0.08 / 0.46 | 0.14 / 0.20 |
| **Reformer** | **3.62 [2.64–4.25]** | **3.24 [2.07–3.62]** | -0.30 | -0.24 | 0.25 / 0.42 | 0.15 / 0.18 |
| DeePNAP | 3.78 [2.82–4.45] | 3.38 [2.23–3.78] | -0.55 | -0.57 | 0.08 / 0.46 | 0.14 / 0.22 |
| FoldX | 9.88 [7.78–11.20] | 8.85 [5.88–9.88] | -0.54 | -0.34 | 0.50 / 0.46 | 0.15 / 0.22 |
| hnRNPK three-C patches |  |  |  |  |  |  |
| Method | RMSE [CI] | MAE [CI] | Pearson $r$ | Spearman $\rho$ | ROC-AUC (S/W) | PR-AUC (S/W) |
| **IRIS** | 24.45 [19.72–28.55] | 19.81 [14.65–24.45] | **0.24** | **0.28** | 0.52 / **0.67** | 0.28 / **0.42** |
| DeepBind | 2.02 [1.60–2.42] | 1.64 [1.19–2.02] | -0.09 | -0.26 | 0.51 / 0.25 | 0.30 / 0.22 |
| Rosetta-Vienna | 1440.18 [110.55–1850.42] | 1166.85 [85.22–1440.18] | -0.21 | -0.07 | 0.37 / 0.62 | 0.21 / 0.39 |
| Reformer | 1.93 [1.27–2.52] | 1.57 [0.94–1.93] | -0.42 | -0.29 | 0.31 / 0.32 | 0.19 / 0.20 |
| **DeePNAP** | **1.43 [1.09–1.75]** | **1.16 [0.81–1.43]** | -0.18 | -0.19 | 0.47 / 0.38 | 0.24 / 0.24 |
| **FoldX** | 11.21 [9.29–13.15] | 9.09 [6.79–11.21] | -0.21 | -0.26 | **0.54** / 0.30 | **0.31** / 0.24 |

For Dataset 3 (Table 4), no single method consistently outperforms all others. IRIS maintains competitive accuracy, particularly for interface-proximal mutations, while Rosetta-Vienna^47^ and FoldX^48^ occasionally achieve better performance. The originally reported Rosetta-Vienna (augmented) results^47^ show stronger performance than those obtained from the public server, likely reflecting system-specific optimization of the scoring function. We note that Dataset 3 originates from early experiments and may partially overlap with the training data used by some external models.

**Table 4:**
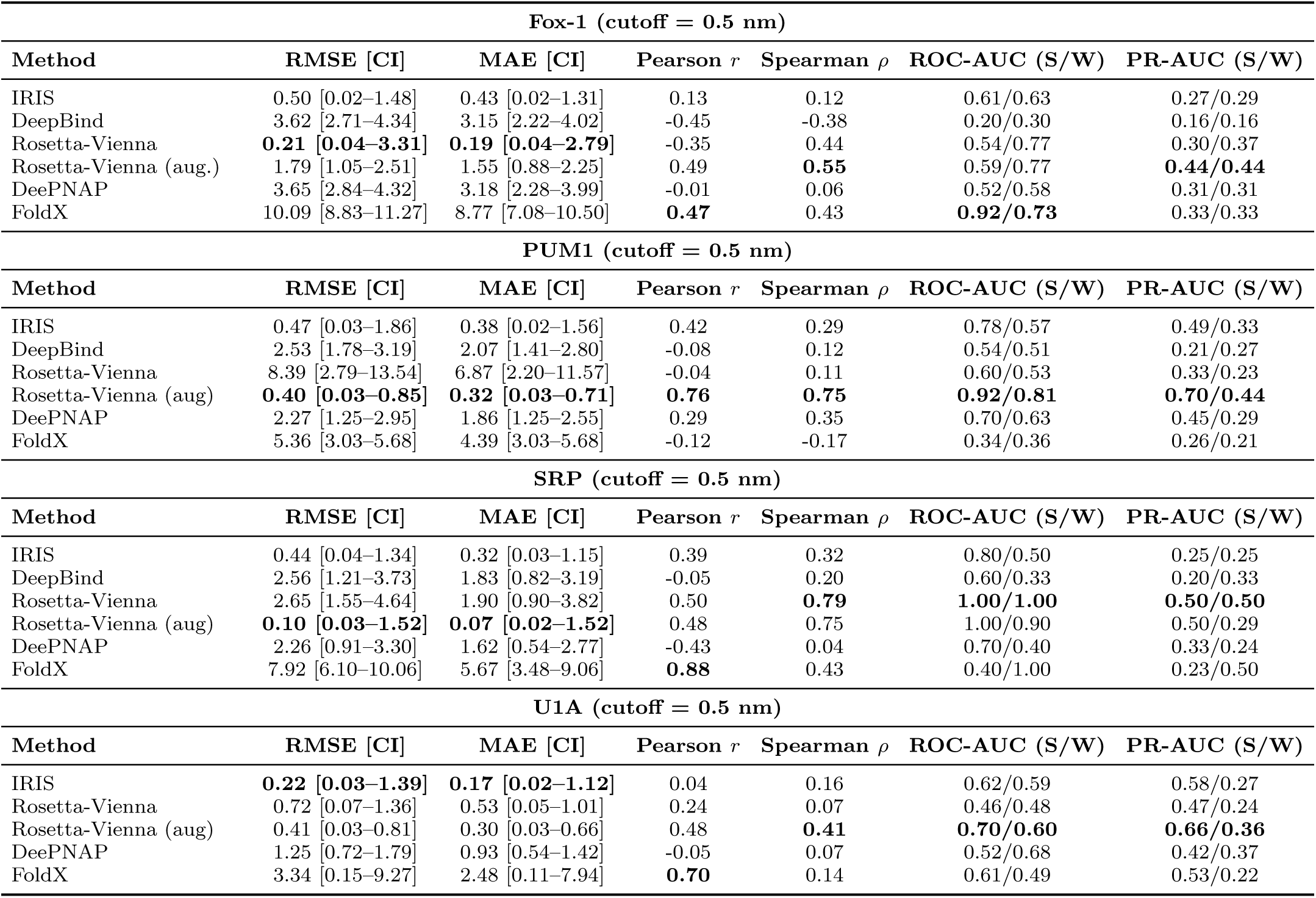
Performance comparison on protein–RNA complexes (Fox-1, PUM1, SRP, and U1A) at the 0.5 nm mutation distance cutoff. Reported metrics include RMSE and MAE (with confidence intervals), Pearson (*r*), Spearman (*ρ*), and classification performance measured by ROC-AUC and PR-AUC (reported as strong/weak binder classification). IRIS predictions were generated using the default parameters described in Table S1, with 0 protein decoys and 10,000 RNA decoys for Fox-1, and 5,000 protein decoys and 10,000 RNA decoys for PUM1, SRP, and U1A. All other parameters correspond to the bolded defaults in Table S1. Reformer was not evaluated due to the lack of pre-trained models for these four proteins. DeepBind was not evaluated for U1A for the same reason.

Overall, IRIS provides a balanced approach that bridges sequence-based learning and structure-informed modeling, enabling robust generalization across diverse protein–RNA systems without relying on task-specific affinity training data.

## Discussion

One of the main challenges in computational characterization of protein–RNA interactions lies in the scarcity of high-quality training data and the inherent flexibility of RNA structures.^58^ These limitations have largely restricted the development of predictive tools capable of estimating sequence-specific protein–RNA binding affinities and identifying high-affinity binding motifs, in contrast to the abundant predictive tools available for protein–DNA interactions.^31,35,57,90–95^ Our results demonstrate that IRIS can mitigate this limitation by leveraging sequence and structural information from only one or a few high-affinity protein–RNA complexes, enabling accurate affinity predictions without requiring large training datasets. Furthermore, the framework can be extended by incorporating structural models predicted by emerging tools such as AF3,^40^ Boltz,^43,44^ Chai-1,^42^ and RoseTTAFoldNA,^41^ highlighting a path toward progressively more accurate and generalizable protein–RNA affinity prediction. Analyses of MS2 coat protein binding demonstrate that IRIS recovers the relative ranking of affinities across a broad spectrum of mutants (Figures 2 and 3). Predictions are generally more reliable when the mutated sequences are similar to the training set and when sequence variations occur near the binding interface, highlighting the importance of local structural context in determining binding affinity. Incorporating AF3-predicted structures reveals alternative binding interfaces and improves predictions for sequences with multiple or distal mutations (Figures 3A and 4). Furthermore, MS2 motif sequences with high Jaccard similarity displayed similar AF3-predicted structures, often adopting the same binding sites (Figure 4). These results indicate that conserved structural patterns among similar motif sequences may underlie IRIS’ predictive accuracy.

Recent advances in deep-learning-based structure predictions have enabled *in silico* modeling of protein-nucleic acid complexes directly from sequences.^40–44^ We evaluated the utility of incorporating these predicted structures into IRIS for protein–RNA binding affinity prediction with hnRNPK (Figure 7). Despite the presence of disordered regions such as the RG/RGG domain, IRIS predictions based on the confidently modeled KH1/2 domains closely aligned with experimental binding data. In contrast, inclusion of lower-confidence regions in training reduces predictive performance, highlighting the importance of high-quality structural input. While these findings based on predicted structures are promising, deep-learning-based structural predictions of protein–RNA complexes are still imperfect, especially when structural templates are unavailable.^96^ Therefore, these structural predictions should be interpreted cautiously, and experimental validation is necessary to confirm both binding affinities and RNA interaction sites.

The success of IRIS in binding affinity prediction demonstrates the importance of structure-sequence relationships underlying protein–RNA interactions. Our structure-informed sequence logo (Figures 3D and 5A) reveals conserved interfacial structure-sequence patterns enriched in the high-affinity binders. Inspired by this, we constructed a k-mer binding motif library (Figure 6) that summarizes strong binding motifs uncovered in this study. With the increasing availability of protein–RNA binding data and more accurate structural predictions, such libraries of high-affinity RNA motifs could facilitate mechanistic understanding of sequences critical for RNA recognition^78,81,82^ and inform the design of therapeutic aptamers.^11,12,97,98^

Although our primary focus has been to delineate conditions under which IRIS provides the most accurate RNA-binding predictions, it consistently demonstrates comparable or improved performance relative to other state-of-the-art models (see Sections *Benchmarking Performance Against Sequence- and Structure-Based Models* and Supplementary Information, *Benchmarking IRIS Predictive Performance with Other Models* ; Figures S3-S5; Tables 2, 3, and 4). For example, IRIS predictions for hnRNPK binding affinities are comparable to leading RNA-affinity prediction models,^35–37,47,49^ while its MS2 predictions outperform all evaluated state-of-the-art models, which themselves surpass earlier approaches such as GLM-Score^99^ and Rosetta.^100^

Nonetheless, because IRIS primarily models sequence variation at the protein–RNA binding interface, it may not fully capture the effects of mutations distal to the binding site.

Such distal effects are likely mediated by allosteric or structural rearrangements, which can be modeled by methods that incorporate RNA secondary and tertiary structure.^47,59,101,102^ As an initial step toward addressing this limitation, we find that incorporating predicted RNA folding energies from ViennaRNA^59^ improves MS2–RNA binding affinity predictions for quadruple mutants (Figure S2; see Supplementary Information Section *Incorporation of Predicted Secondary Structure in IRIS Prediction*). Future integration of IRIS with such approaches may enable more comprehensive modeling that captures both interface-proximal and distal sequence effects.

Finally, *in vitro* predictions of protein–RNA binding affinity and specificity represent only an initial step toward a comprehensive understanding of the cellular interactome.^103–105^ *In vivo*, additional factors such as competition among RNA-binding proteins,^9^ RNA modifications,^106,107^ and phase separation^6,7,108,109^ significantly influence binding outcomes. Incorporating these complexities will extend IRIS beyond *in vitro* settings and facilitate its translation toward clinically relevant and therapeutic applications.^11,12,97,98^

## Acknowledgment

We are grateful to the Reformer authors for their assistance in using the pretrained hnRNPK model. We acknowledge the use of AI tools, such as ChatGPT and Grammarly, for grammar checking and improving the clarity of the manuscript.

## Author Contributions

Conceptualization: E.C.R, Y.Z., X.L.

Methodology: E.C.R., Y.Z., X.L.

Investigation: E.C.R., Y.Z., X.L.

Visualization: E.C.R, Y.Z., X.L.

Supervision: X.L.

Writing—original draft: E.C.R., Y.Z., X.L.

Writing—review & editing: E.C.R., Y.Z., X.L.

## Data and Materials Availability

The implementation of the IRIS model, along with training and prediction examples, is available at our GitHub repository. All training and testing datasets used in this study were collected from previously published articles and are publicly available from open-source repositories.

