## Supplementary Information for "IRIS Integrates Sparse Sequence, Experimental, and AI-Predicted Structures for Protein–RNA Affinity Prediction and Motif Discovery"

§*Contributed equally to this work*

#### Summary

**1 Effect of Sequence Similarity on IRIS Predictions** **S-3**

**2 Benchmarking IRIS Predictive Performance with Other Models** **S-4**

**3 Parameter Calibration and Error Analysis of IRIS Predictions** **S-7**

**4 Classification of Strong and Weak Protein–RNA Binders** **S-7**

**5 Structural Interpretation of Protein–RNA Binding Interface** **S-9**

**6 Selection of Training Sequences for Augmenting IRIS Prediction** **S-9**

**7 Incorporation of Predicted Secondary Structure in IRIS Prediction** **S-11**

**8 Quantifying Sequence and Structural Contributions of IRIS** **S-12**

**9 Supplementary Figures** **S-14**

**10 Supplementary Tables** **S-30**

### 1 Effect of Sequence Similarity on IRIS Predictions

To understand differences between models trained on the native crystal structure (PDB ID: 2C4Q) alone versus those augmented with additional sequences (B4 and B5), we examined how IRIS interprets sequence similarity between training and test variants.

IRIS predicts binding free-energy changes ($\Delta\Delta G$) by evaluating how sequence perturbations alter protein–RNA interaction patterns relative to the training template. Consequently, predicted $\Delta\Delta G$ values increase monotonically with the number of substitutions from the template sequence (Figure S13A). In contrast, the experimental measurements exhibit a characteristic flag-shaped distribution: the maximum $\Delta\Delta G$ remains relatively stable across mutation classes, whereas the minimum increases with mutational load. This discrepancy is most pronounced for quadruple mutants, where highly divergent variants can still retain strong binding affinity.

To quantify similarity in terms of interaction-relevant sequence content, we computed Jaccard similarity between training and test sequences. This metric captures overlap in sequence elements that contribute to protein–RNA interactions and more closely reflects the internal scoring mechanism of IRIS. Although it follows trends similar to sequence identity, Jaccard similarity provides a clearer view of how sequence overlap influences predicted $\Delta\Delta G$ values (Figure S13B). The analysis reveals a systematic bias: models trained on a single structural template tend to over-penalize distant variants and underestimate the diversity of high-affinity sequences.

Including additional sequences (B4 and B5) expands the training set and reshapes the similarity landscape between training and test variants. This augmentation reduces residual errors for single-, double-, and triple-mutant predictions and corrects mispredictions associated with specific outlier variants (Figure S13C). Importantly, predictions for sequences already well described by the original template remain largely unchanged. These results demonstrate that expanding the diversity of training sequences improves IRIS predictive accuracy by better representing the accessible sequence landscape of the protein–RNA interaction.

#### 2 Benchmarking IRIS Predictive Performance with Other Models

To evaluate the predictive performance of IRIS on protein–RNA interactions, we benchmarked it against several state-of-the-art computational tools using the three datasets described in the main text Table 1 and the *Materials and Methods* section *Summary of Datasets in this Study.* These tools were selected because they represent the current state of the art in machine learning and biophysical modeling, and they provide well-maintained, user-friendly implementations.

We first benchmarked IRIS against DeepBind,[S1] a widely used deep-learning framework for modeling sequence specificities of RNA-binding proteins. Each RNA sequence was submitted to the DeepBind web server (`http://www.rnainter.org/DeepBind/`), which returns a binding-specificity score directly from sequence input. Because DeepBind does not currently provide an MS2-specific model and the U1A dataset mainly contains protein mutation data rather than RNA sequence variation, we focused the comparison on hnRNPK–RNA binding prediction and three additional protein datasets where sequence-based prediction is applicable.

We next evaluated Rosetta-Vienna RNP-$\Delta\Delta G$,[S2] a method that integrates thermodynamic calculations with RNA secondary structure predictions and has demonstrated improved accuracy over earlier approaches such as GLM-Score[S3] and Rosetta-based molecular modeling.[S4] For MS2 predictions and the four additional proteins, we used the same PDB structures as those used by IRIS. For each mutant RNA sequence, RNA folding free energies were computed using the ViennaRNA package.[S5] These sequences and corresponding folding energies were then submitted to the RNP-$\Delta\Delta G$ server (`https://rosie.rosettacommons.org/rnp_ddg`), which performs automated structural relaxation and $\Delta\Delta G$

calculations (Figure S14A). We also compared IRIS predictions with an augmented version of Rosetta-Vienna RNP-$\Delta\Delta G$, in which model predictions were optimized using half of the experimental MS2–RNA binding data (Supplementary Data: https://www.pnas.org/doi/suppl/10.1073/pnas.1819047116/suppl_file/pnas.1819047116.sd01.txt; labeled as Rosetta-Vienna (aug.) in Table 1). To ensure fair comparisons using the same test set, we excluded sequences used during optimization and those containing quadruple mutations (Figure S14B).

For hnRNPK predictions, we employed the same AlphaFold3 (AF3)-generated protein–RNA complex used by IRIS. This structure was first relaxed into 30 conformations using the ROSIE portal (https://r2.graylab.jhu.edu/auth/login?next=/apps/view/relax/41201)[S6] to optimize structural geometry. The relaxed structures and corresponding folding energies were then submitted to the RNP-$\Delta\Delta G$ server for binding energy estimation. We thank the ROSIE team for maintaining a public record of webserver outputs (IDs: 127677, 127732–127735, 127752–127753, 131413, 131515, 131519, 131614) generated during our predictions, which enabled direct verification of the benchmarks reported here.

We examined Reformer,[S7] a transformer-based architecture designed for modeling protein–RNA interactions at single-base resolution. We used the pre-trained hnRNPK model (prefix HNRNPK_K562) available at https://github.com/xilinshen/Reformer, following the authors published protocol. Because the currently released Reformer weights are restricted to human RNA-binding proteins, the method cannot be applied to the viral MS2 protein. For the remaining datasets, Reformer provides a trained model for PUM1. However, Reformer was specifically designed for RNA sequence mutations affecting binding affinity, whereas the PUM1 dataset primarily contains protein mutations, limiting its applicability. Therefore, we restricted the reformer predictions to the hnRNPK-RNA dataset (Dataset 2).

We evaluated DeepPNAP,[S8] a recently proposed sequence-based machine learning framework for predicting protein–RNA binding affinities. Following the authors GitHub repository (https://github.com/StructuralBiologyLabIISERTirupati/DeePNAPWebsite), we implemented a customized script to enable batch prediction of multiple RNA sequences simultaneously. DeepNAP is flexible for sequence input and does not require retraining for new sequences.

We tested FoldX,[S9] a widely used structure-based method for energy calculation and molecular design, that has been refined using large-scale mutation datasets for more than two decades. Its recent versions support protein–RNA binding affinity prediction.[S10] Because FoldX requires PDB structures as input, we replaced the RNA sequence of the native protein–RNA structure with target sequences, which were processed using the `RepairPDB` command to correct structural issues and ensure compatibility with FoldX. We then used the `AnalyseComplex` command to estimate the binding affinity.

We evaluated PredPRBA,[S11] a biophysics-informed model requiring PDB structural input. However, when applied to SRP–RNA complexes with target-sequence replacement (Dataset 3), the model produced identical predictions across all variants, indicating a lack of sensitivity to sequence changes in this setting. We further tested PredPRBA using AF3-predicted hnRNPK–RNA structures (Dataset 2), but observed poor correlation with experimental binding affinities. Additionally, its implementation as an online server (`http://predprba.denglab.org/`) limits scalability for systematic benchmarking. For completeness, we report its performance on the hnRNPK 10bio dataset as a reference.

To provide a comprehensive evaluation of predictive performance, we report the following metrics: (1) Pearson correlation coefficient to measure the linear correlation between predicted and experimental $\Delta\Delta G$ values; (2) Spearmans rank correlation coefficient to assess agreement in ranked ordering; (3) ROC-AUC (Area Under the Receiver Operating Characteristic Curve); (4) precision–recall AUC (PR-AUC) to evaluate the models ability to distinguish high-affinity from low-affinity binding sequences; and (5) MAE and RMSE values calculated with the fitted regression line through the origin. However, we need to be cautious when comparing models using MAE/RMSE values, as models' original outputs are on different scales.

### 3 Parameter Calibration and Error Analysis of IRIS Predictions

To evaluate the calibration of IRIS predictions, we compared predicted and experimental $\Delta\Delta G$ values using linear regression models with both free-intercept and origin-constrained fits. When the intercept is allowed to vary (Figure S5), predictions for single to triple mutants show strong agreement with experimental measurements. However, correlation decreases for higher-order mutants, particularly variants containing quadruple substitutions. This trend reflects the increasing difficulty of accurately predicting binding energies for sequences that diverge substantially from the structural template used during model construction. Regression constrained through the origin (Figure S4) isolates slope calibration independent of intercept offsets. Comparing the two regression approaches suggests that part of the systematic error arises from a constant offset between predicted and experimental $\Delta\Delta G$ values. The free-intercept fit can absorb this offset, making residuals appear smaller and more normally distributed (Figure S7), whereas the origin-constrained fit removes this flexibility, revealing the underlying systematic error more clearly (Figure S6).

For single-to-triple substitutions, residuals remain approximately normally distributed, indicating stable predictive behavior. In contrast, residuals for quadruple substitutions become more uniformly distributed, reflecting increased variability as mutational load increases. Together, these analyses provide a quantitative assessment of IRIS prediction accuracy across mutation classes and highlight the model's limits when extrapolating to highly mutated RNA variants.

### 4 Classification of Strong and Weak Protein–RNA Binders

In addition to predicting continuous binding free energies, IRIS can also be used to classify RNA variants according to binding strength. To evaluate this capability, we performed rank-based classification analyses separating strong and weak binders.

Strong binders were defined as variants within the lowest quartile of experimentally measured $\Delta\Delta G$ values, whereas weak binders corresponded to the highest quartile. Using these definitions, receiver operating characteristic (ROC) and precision-recall (PR) curves were computed by applying rank thresholds to predicted $\Delta\Delta G$ values (Figures S15 and S16). The resulting classification metrics across complexes and protein decoy conditions are summarized in Table S5.

For the MS2 system (2C4Q, 0–3 substitutions), IRIS achieves strong discrimination of high-affinity variants, with ROC-AUC values of approximately 0.83 for strong binders under low–protein decoy conditions. However, classification performance decreases as the number of protein decoys increases, consistent with the reduction in correlation observed for these conditions (Table S5). In contrast, MS2 variants containing four or more substitutions show near-random classification performance, reflecting the reduced predictive accuracy for highly mutated sequences.

Across additional protein–RNA complexes, classification performance varies depending on the system and decoy sampling level. For example, the PUM1–RNA complex exhibits relatively strong discrimination of strong binders, with ROC-AUC values reaching $\sim$0.77 and PR-AUC values $\sim$0.57 under moderate protein decoy sampling. Similarly, the SRP–RNA complex improved classification with increasing protein decoy sampling, reaching ROC-AUC values of approximately 0.80 for strong binders. In contrast, systems such as FOX1–RNA and U1A–RNA exhibit poor classification performance across any protein decoy sampling.

Overall, IRIS predictions capture meaningful energetic trends that enable identification of strong RNA binders in several protein–RNA systems, while also highlighting variability in classification performance across structural templates and mutation regimes.

### 5 Structural Interpretation of Protein–RNA Binding Interface

Beyond predictive accuracy, IRIS provides interpretable interaction maps that reveal how sequence mutations perturb protein–RNA contacts. To characterize key protein–RNA interactions that contribute to binding, and to illustrate the alternative protein–RNA binding interface revealed by structural predictions, we constructed residue-based contact maps from the MS2–RNA crystal structure (PDB ID: 2C4Q) and the AF3-predicted structures of B1 to B7 variants (Figure S12).

These aligned contact maps for the MS2–RNA systems show that B1-B6 variants maintain similar interaction patterns to the native crystal structure. In contrast, B7 exhibits pronounced differences in protein–RNA contacts, confirming an alternative binding interface that leads to distinct binding behavior. These structural analyses demonstrate how IRIS predictions can be interpreted mechanistically, linking sequence mutations to changes in protein–RNA interaction interface and binding energetics.

### 6 Selection of Training Sequences for Augmenting IRIS Prediction

We hypothesized that increasing the diversity of training sequences would improve the IRIS prediction performance. To test this, we analyzed strong-binding quadruple-mutation sequences (B1 - B7) that were poorly predicted by the original single-structure IRIS training scheme. Sequence-level analysis using Jaccard similarity across multiple k-mer lengths revealed differences in motif conservation, with B2, B4, B5, and B7 showing the greatest divergence from the native sequence associated with the crystal structure (PDB ID: 2C4Q) (Figure S11A).

To further quantify sequence diversity, we performed principal component analysis (PCA) on positional 3-mer representations of the RNA sequences. Each RNA sequence was divided into three equal-length regions to retain positional information. Within each region, frequencies of all possible 3-mers over the RNA alphabet {A, C, G, U} were computed, yielding a 64-dimensional vector per region ($4^3 = 64$). These regional vectors were concatenated to form a 192-dimensional feature representation for each sequence.

To reduce the influence of absolute count differences, 3-mer frequencies within each region were normalized by the total number of observed 3-mers, and the resulting 192-dimensional feature vectors were further L2-normalized prior to PCA. Projection into two dimensions revealed that B4, B5, and B7 are the most distant from the native sequence along the first principal component (Figure S11B), indicating substantial deviations in coarse-grained positional 3-mer composition.

Among these variants, B7 exhibits the greatest divergence and, as shown in Section *Alternative Protein-RNA Binding Interfaces Revealed by Structural Prediction and Motif Analysis*, adopts an alternative protein–RNA binding interface. Because this structural shift is incompatible with the original crystal structure, B7 cannot be incorporated into the same training framework. We therefore selected B4 and B5 to increase sequence diversity while preserving structural consistency.

We further evaluated RNA secondary structure using the ViennaRNA package.[S5] Among the candidate sequences, B4 retains the greatest similarity to the native secondary structure (Figure S11C), consistent with its strong predictive performance in models trained on individual B1-B7 variants (Figure 3A).

Overall, incorporating B4 and B5 balances increased sequence diversity with preservation of the protein–RNA structural interface, leading to reduced prediction error and improved accuracy (Figures 3B and 3C).

### 7 Incorporation of Predicted Secondary Structure in IRIS Prediction

We examined whether predicted RNA secondary structure could provide complementary information to improve IRIS-based affinity prediction. For each RNA sequence, secondary structure was predicted using the ViennaRNA package,[S5] and the corresponding minimum free energy (MFE) was used as a proxy for structural stability. Because IRIS-predicted binding free energies and ViennaRNA-predicted MFEs are reported on different scales, both quantities were transformed into normalized Z-scores using $Z = \frac{\text{score} - \mu_{\text{score}}}{\sigma_{\text{score}}}$, where $\mu_{\text{score}}$ and $\sigma_{\text{score}}$ denote the mean and standard deviation, respectively.

Previous studies have shown that many RNA-binding proteins preferentially bind to less structured RNA motifs.[S2,S12,S13] Consistent with this trend, the seven high-affinity outlier sequences (B1–B7) in the MS2 quadruple-mutant dataset exhibit higher secondary-structure MFEs than the remaining quadruple-mutation sequences, indicating reduced structural stability. This observation suggests that secondary-structure stability contains information not fully captured by the IRIS binding score alone. As a preliminary approach, we combined two normalized Z-scores, from the IRIS-predicted binding free energy and the ViennaRNA-predicted MFE, into a composite score, $Z_{\text{Composite}} = Z_{\text{IRIS}} - Z_{\text{ViennaRNA}}$, to evaluate whether incorporating predicted secondary structure stability improves IRIS predictive performance.

As shown in Figure S2B, $Z_{\text{Composite}}$ improved prediction accuracy for the complete set of MS2 quadruple mutants relative to the IRIS-only model (Figure 2A bottom), with Pearson correlation increasing from 0.05 to 0.20 and Spearman correlation increasing from 0.22 to 0.36. The improvement was primarily driven by better recovery of the seven high-affinity outlier sequences. Nonetheless, for variants containing one to three mutations, inclusion of the ViennaRNA-derived term reduced predictive accuracy compared with the original IRIS model. Together, these results suggest that RNA secondary structure can act as a useful complementary predictor in specific mutational regimes, while also indicating that more refined integration strategies may be required. Future efforts may benefit from incorporating alternative RNA secondary-structure prediction models, such as EternaFold[S14] and BPFold.[S15]

We also evaluated whether incorporating the AF3-predicted structural ensemble could improve IRIS-based affinity prediction for the MS2 mutant series. In contrast to the secondary-structure-based composite score, AF3-derived structural ensembles did not generally improve predictive performance (Figure 3A). Most AF3-predicted mutant structures yielded no measurable gain in accuracy relative to the original IRIS model, with the notable exception of B7, for which AF3 suggested an alternative binding interface. This result indicates that, although AF3 can occasionally reveal potentially informative interface rearrangements, it lacks sufficient sensitivity to reliably capture subtle mutation-induced interface changes, particularly those caused by small sequence perturbations such as point mutations.[S16] Therefore, our results suggest that AF3 is more useful for generating plausible protein–RNA complex structures when experimental structures are unavailable, as in the hnRNPK–RNA case, than for routine use of mutant-specific predicted structural ensembles in IRIS affinity prediction.

### 8 Quantifying Sequence and Structural Contributions of IRIS

We performed an ablation study to compare the IRIS performance with sequence-only and structure-only modes.

To evaluate the sequence-only mode, we used two approaches, one based on the Hamming distance from the native sequence associated with the crystal structure, the other based on the Jaccard Similarity using 3-mers (see *Materials and Methods* Section *Jaccard Similarity Heatmap of RNA Sequences Against Known MS2 Motifs* for details). Performance was assessed using multiple metrics, including MAE, RMSE, Pearson correlation coefficient, Spearman's rank correlation coefficient, ROC-AUC, and PR-AUC. MAE and RMSE values should be interpreted with caution, as these predictors are not directly comparable in scale to IRIS predictions.

To evaluate the performance of the structural-only mode, we retained the component of IRIS that extracts amino-acid-nucleotide contact frequencies from training structures, while randomizing the sequence input. To fully sample sequence space for a robust evaluation, we performed 100 independent simulation runs, each using a randomly selected sequence to train the IRIS model based on the same input structure. By computing the mean and standard deviation of metrics across all 100 runs, we established a baseline to quantify how structural features contribute to the predictive accuracy and robustness of the IRIS framework.

Detailed results are reported in Table S10. The sequence-exclusive mode shows a weak positive correlation with experimentally measured binding free energies, indicating that sequence information contributes to binding affinity but must be correctly extracted. The structure-only mode exhibits near-random performance, suggesting that although structural information is useful, it needs to be integrated with correct sequence information to enable accurate model predictions. By integrating both sequence and structural information through calculating sequence-specific contact frequencies at the protein-RNA interface, IRIS achieves substantially higher predictive accuracy.

### 9 Supplementary Figures

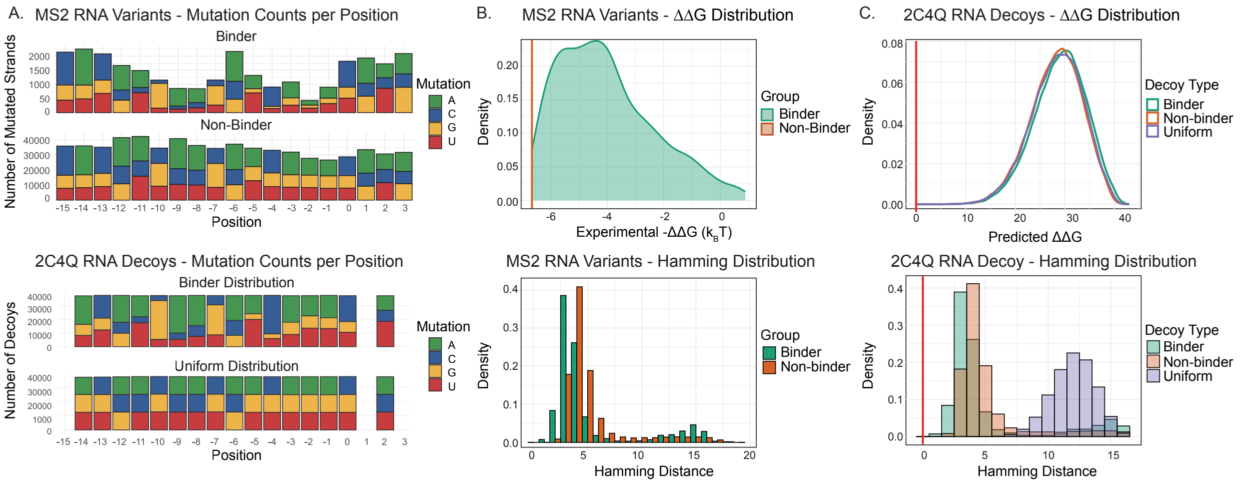


Figure S1: **MS2 mutation landscape, decoy sampling distributions, and binding energetics of RNA variants.** (A) Top: Strand-level mutation counts per RNA position across all experimental MS2-RNA variants, separated into binders and non-binders. For each nucleotide position relative to the native sequence, the number of variants carrying a substitution is shown, highlighting positional mutation tolerance and regions enriched for mutations in each class. Bottom: Mutation counts per position for RNA decoys generated from the PDB:2C4Q structure as the training template. Decoys were constructed either to match the experimental binder distribution or follow a uniform distribution across all RNA positions, enabling comparison between biologically informed and uniform sampling strategies. (B) Top: Distribution of experimentally measured $-\Delta\Delta G$ $(kT)$ values for MS2 RNA variants, separated into binders and non-binders using a classification threshold at $-\Delta\Delta G = -6.6617$ $kT$ (red line). Bottom: Corresponding Hamming distance distributions for binders and non-binders, where Hamming distance is defined as the number of nucleotide substitutions relative to the native sequence, illustrating differences in mutational burden between functional classes. (C) Top: Predicted $\Delta\Delta G$ distributions for RNA decoys generated under different sampling strategies (uniform, binder-matched, and non-binder-matched), while maintaining a uniform Hamming distance distribution across sets, and compared to the native sequence from the 2C4Q training template (red vertical line). Bottom: Hamming distance distributions for each decoy set, showing sampling strategy shapes sequence divergence relative to the native state. The native sequence is shown as a reference (red line).

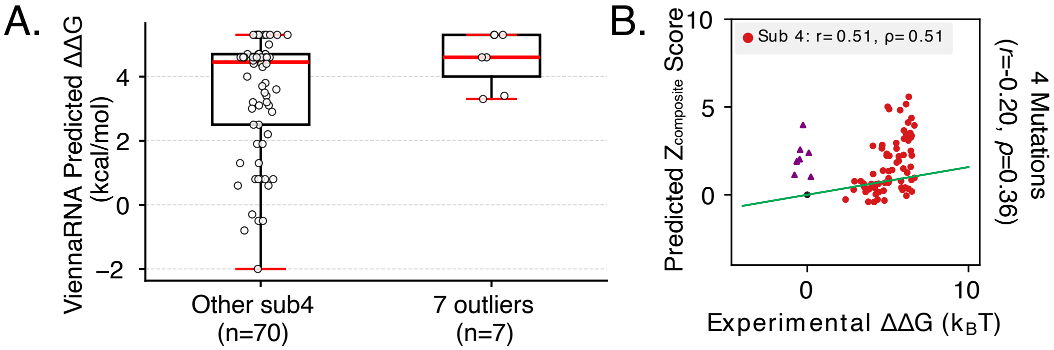


Figure S2: **Integrating RNA folding energy from Vienna improves MS2–RNA binding affinity predictions for quadruple mutants.** (A) ViennaRNA-predicted folding free energy changes ($\Delta\Delta G$), computed from minimum free energy (MFE) estimates as $\Delta\Delta G = \Delta G_{\text{mut}} - \Delta G_{\text{WT}}$, for quadruple-mutation sequences are shown as box plots, with medians indicated in red and interquartile ranges represented by the boxes. The seven outliers consistently exhibit higher $\Delta\Delta G$ values, indicating reduced structural stability relative to the wild-type and compared to the other quadruple-mutation sequences. (B) Correlation between predicted Z-composite scores and experimental $\Delta\Delta G$ values. Incorporating secondary structural information improves the agreement with experimental measurements, particularly for sequences that deviate from the main trend, supporting the role of RNA secondary structure in explaining these outliers.

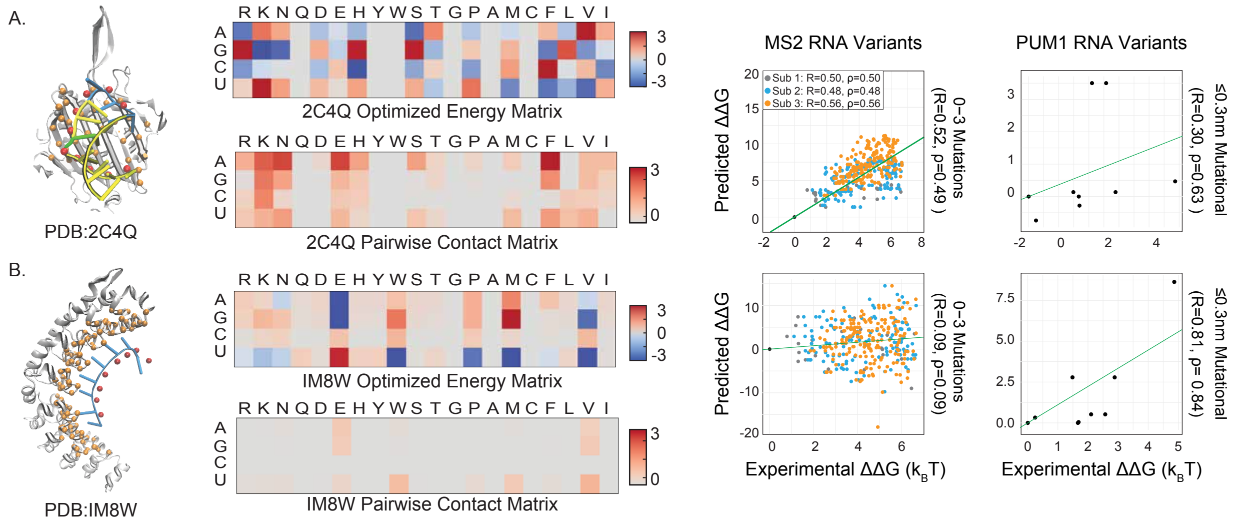


Figure S3: **Template-specific optimized energy matrices capture binding energetics of their respective protein–RNA systems.** (A) MS2–RNA complex (PDB: 2C4Q). Left: The 2C4Q structure, with protein chains highlighted by C$\alpha$ atoms and RNA chains by phosphorus atoms. Middle: Optimized interaction energy matrix derived from 2C4Q template, shown alongside the corresponding protein–RNA contact frequency matrix. Right: Comparison between experimentally measured $\Delta\Delta G$ values and predictions from the 2C4Q-derived energy matrix for MS2 and PUM1 RNA variants, restricted to mutation classes containing 0–3 substitutions or an interface distance cutoff of $\leq$ 0.3nm. The 2C4Q-derived matrix shows strong agreement with MS2 variants but only modest correlation with PUM1 variants, consistent with its specificity to the MS2 binding interface. (B) PUM1–RNA complex (PDB: IM8W). Left: The IM8W structure, with protein chains represented by C$\alpha$ atoms and RNA chains by phosphorus atoms. Middle: Optimized interaction energy matrix and corresponding protein–RNA contact frequency matrix derived from the IM8W template. Right: Comparison between experimentally measured $\Delta\Delta G$ values and predictions from the IM8W-derived energy matrix for both MS2 and PUM1 RNA variants, restricted to the same mutation classes or interface distance cutoff of $\leq$ 0.3 nm.. The IM8W-derived matrix shows strong agreement with PUM1 variants but minimal correlation with MS2 variants, reflecting its specificity to the PUM1 protein–RNA binding interface.

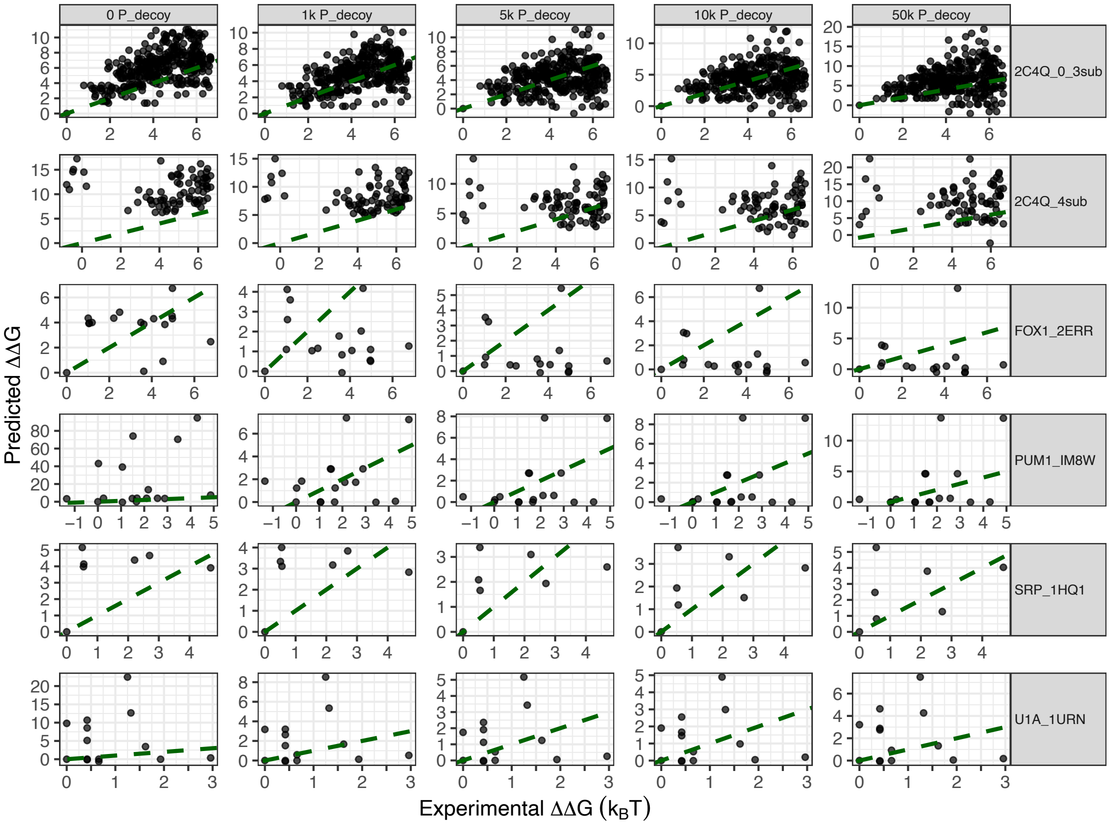


Figure S4: **Correlation between predicted and experimental binding free energies using regression constrained through the origin.** Predicted versus experimental $\Delta\Delta G$ values are shown for each protein–RNA complex across protein-decoy sampling conditions (0, 1k, 5k, 10k, and 50k $P_{\text{decoy}}$). Points represent individual RNA variants that pass the mutation distance cutoff ($\leq 0.5$nm). The dashed green line indicates the origin-constrained fit ($y = x$). Complexes are arranged by facet rows; MS2 (2C4Q) variants are further separated into sequences containing 0–3 substitutions and those with 4 substitutions. Axes are scaled independently within each complex to preserve the dynamic range of predicted and experimental energies.

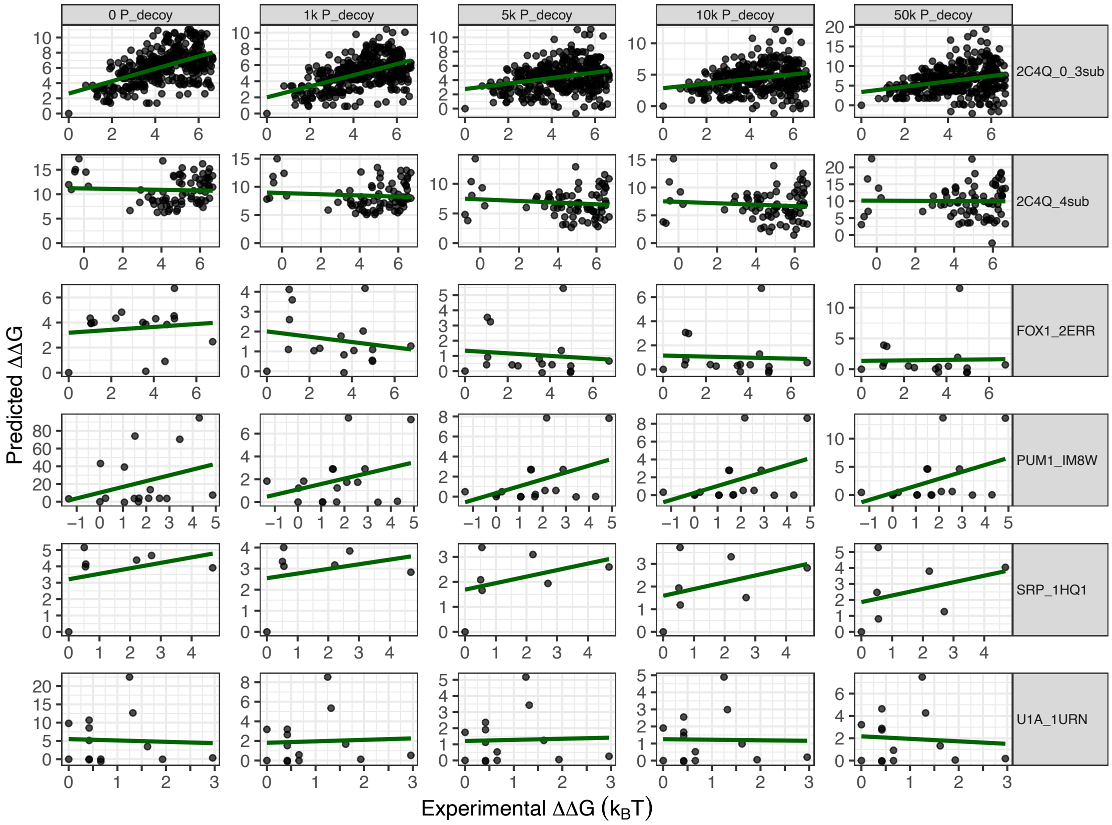


Figure S5: **Correlation between predicted and experimental binding free energies using linear regression with a free intercept.** Predicted versus experimental $\Delta\Delta G$ values are shown for each protein–RNA complex and protein-decoy condition. Points represent individual RNA variants that pass the mutation distance cutoff ($\leq$ 0.5nm). Within each panel, a linear model with a free intercept ($y = \beta_0 + \beta_1 x$) is fitted independently and shown as a solid green line. Complexes are arranged by facet rows; MS2 (2C4Q) variants are further separated into sequences containing 0–3 substitutions and those with 4 substitutions. Axes are scaled independently within each complex to preserve the dynamic range of the predicted and experimental energies.

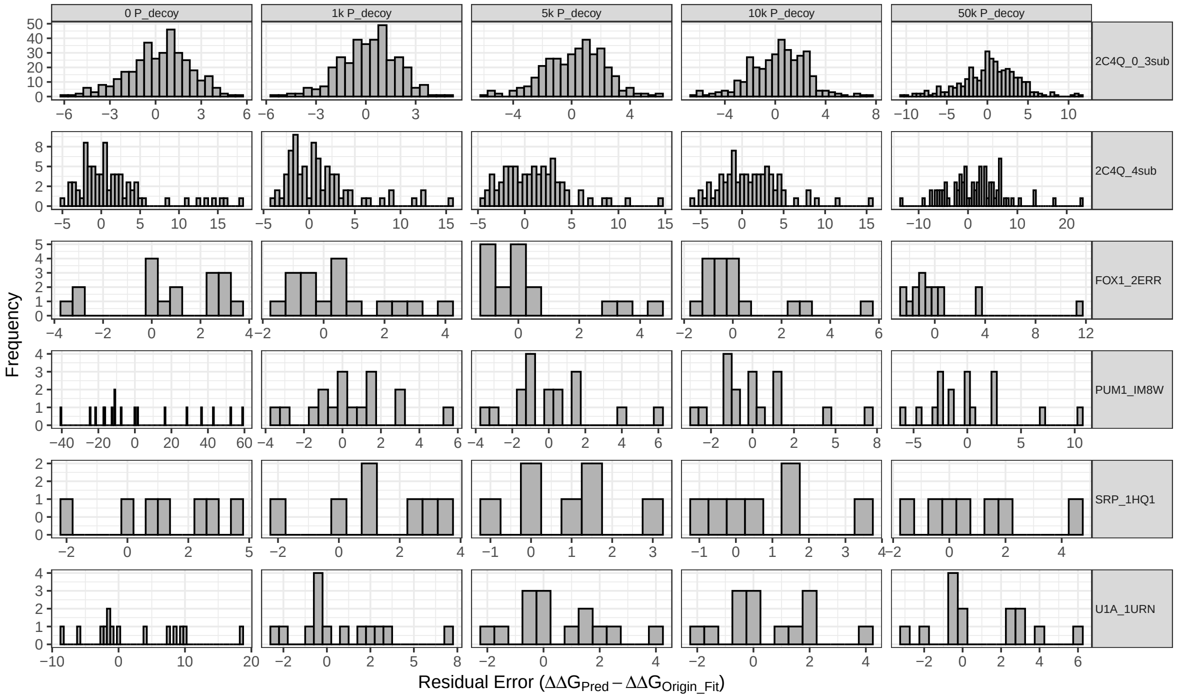


Figure S6: **Residual error distributions for origin-constrained regression.** Histograms of residual errors computed from the regression constrained through the origin (Figure S4). Residuals are defined as $\Delta\Delta G_{\text{pred}} - (\beta_{\text{origin}} \cdot \Delta\Delta G_{\text{exp}})$, where $\beta_{\text{origin}}$ is the fitted slope. Distributions are shown separately for each complex and protein-decoy condition. The bin width is $0.5\text{k}_\text{B}\text{T}$, and x-axis ranges vary across panels to reflect complex-specific error dispersion. Compared with the free-intercept model, these distributions highlight systematic offsets that are absorbed by the intercept in unconstrained regression.

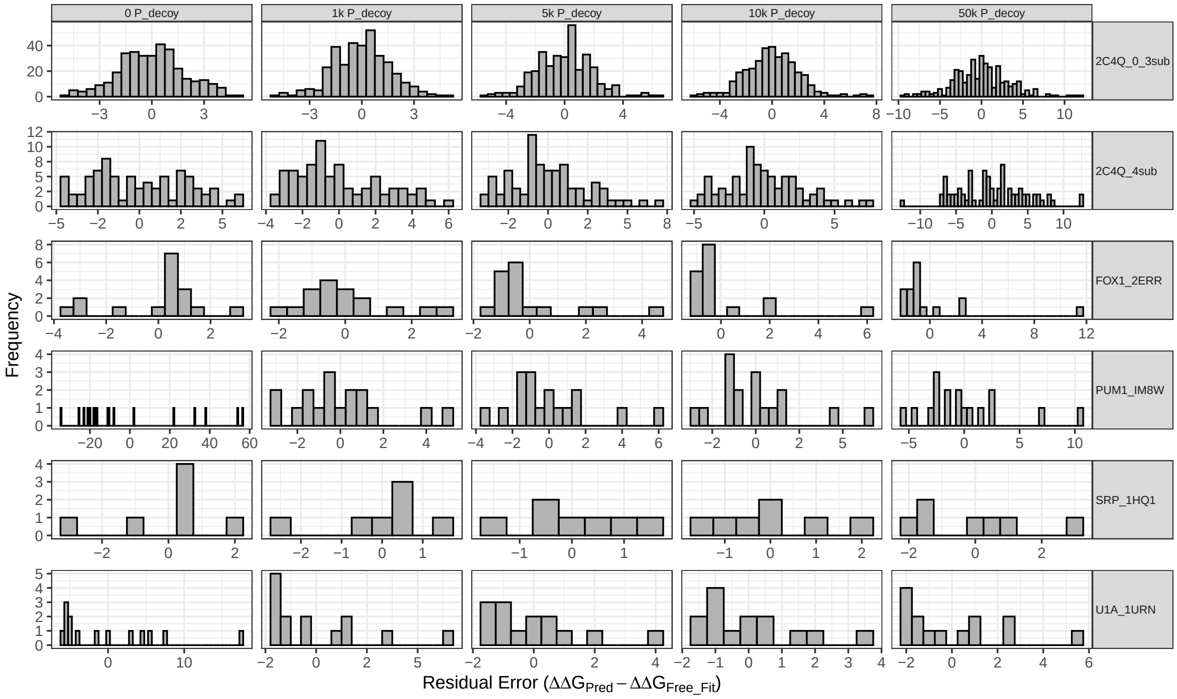


Figure S7: **Residual error distributions for free-intercept regression.** Histograms of residual errors ($\Delta\Delta G_{\text{pred}} - \Delta\Delta G_{\text{fit}}$) derived from the free-intercept linear model shown in Figure S5. Residuals are defined as $\Delta\Delta G_{\text{pred}} - (\beta_0 + \beta_1 \cdot \Delta\Delta G_{\text{exp}})$, where $\beta_0$ and $\beta_1$ are the fitted intercept and slope, respectively. Distributions are shown separately for each complex and protein-decoy condition. The bin width is $0.5\text{k}_\text{B}\text{T}$, and x-axis ranges vary across panels to reflect complex-specific error dispersion. These histograms quantify the remaining bias and variance after intercept calibration.

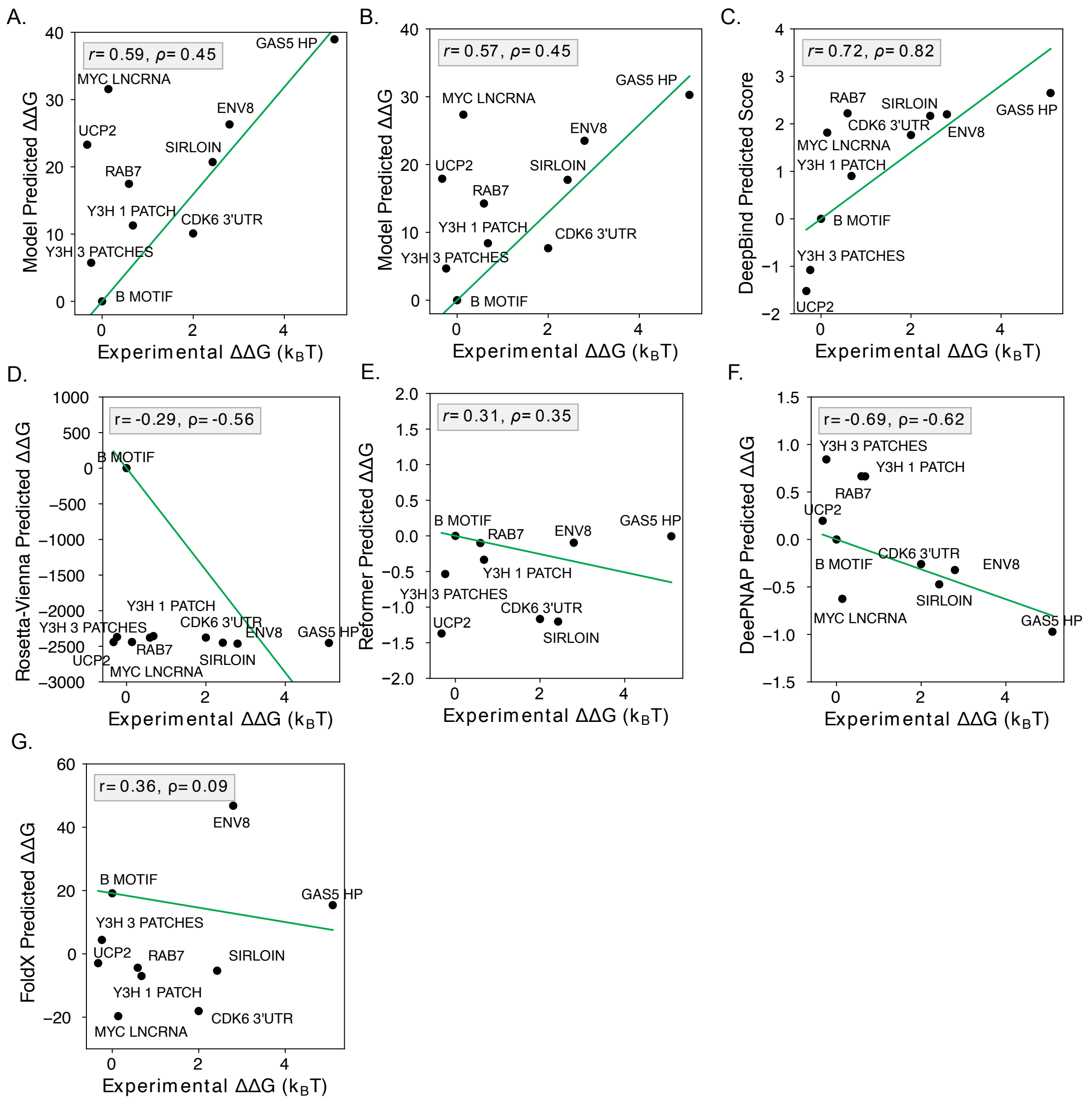


Figure S8: **Comparison of IRIS models trained on additional hnRNPKB-motif binding interfaces and other state-of-the-art methods for predicting hnRNPK binding free energies across biological RNA sequences.** (A) IRIS trained on KH1+KH2+RG/RGG-B-motif binding interfaces. (B) IRIS trained on KH1+KH2+RG/RGG+KH3 (full-length hnRNPK)–B-motif binding interfaces. (C) DeepBind.[S1] (D) Rosetta-Vienna.[S2] (E) Reformer.[S7] (F) DeepPNAP.[S8] (G) FoldX.[S10]

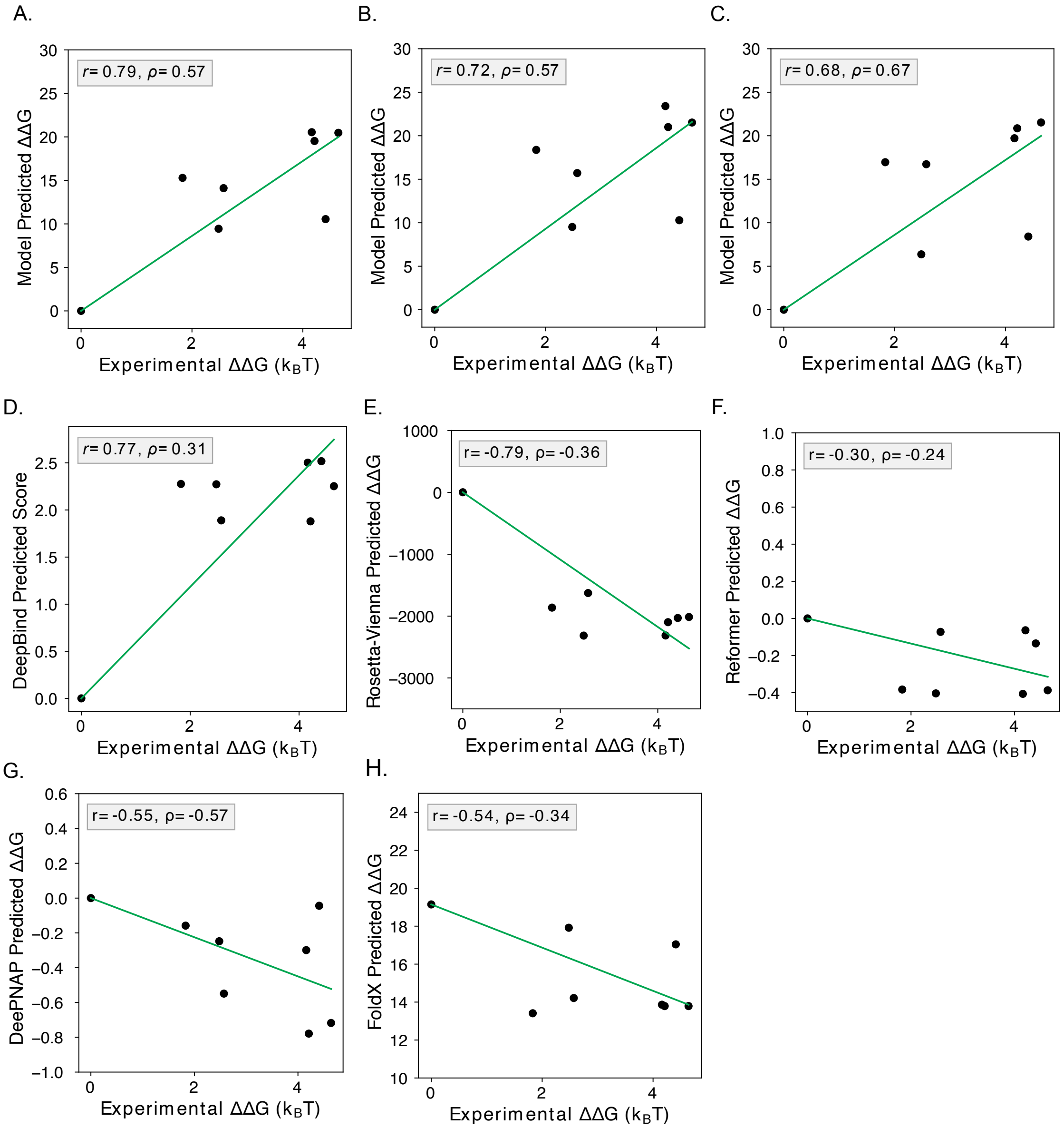


Figure S9: **Comparison of IRIS models trained on different hnRNPKB-motif binding interfaces and other state-of-the-art methods for predicting hnRNPK binding free energies across B-motif variants containing two cytosine patches.** (A) IRIS trained on KH1+KH2-B-motif binding interfaces (same as in Figure 7). (B) IRIS trained on KH1+KH2+RG/RGG-B-motif binding interfaces. (C) IRIS trained on KH1+KH2+RG/RGG+KH3 (full-length hnRNPK)-B-motif binding interfaces. (D) DeepBind.[S1] (E) Rosetta-Vienna.[S2] (F) Reformer.[S7] (G) DeepPNAP.[S8] (H) FoldX.[S10]

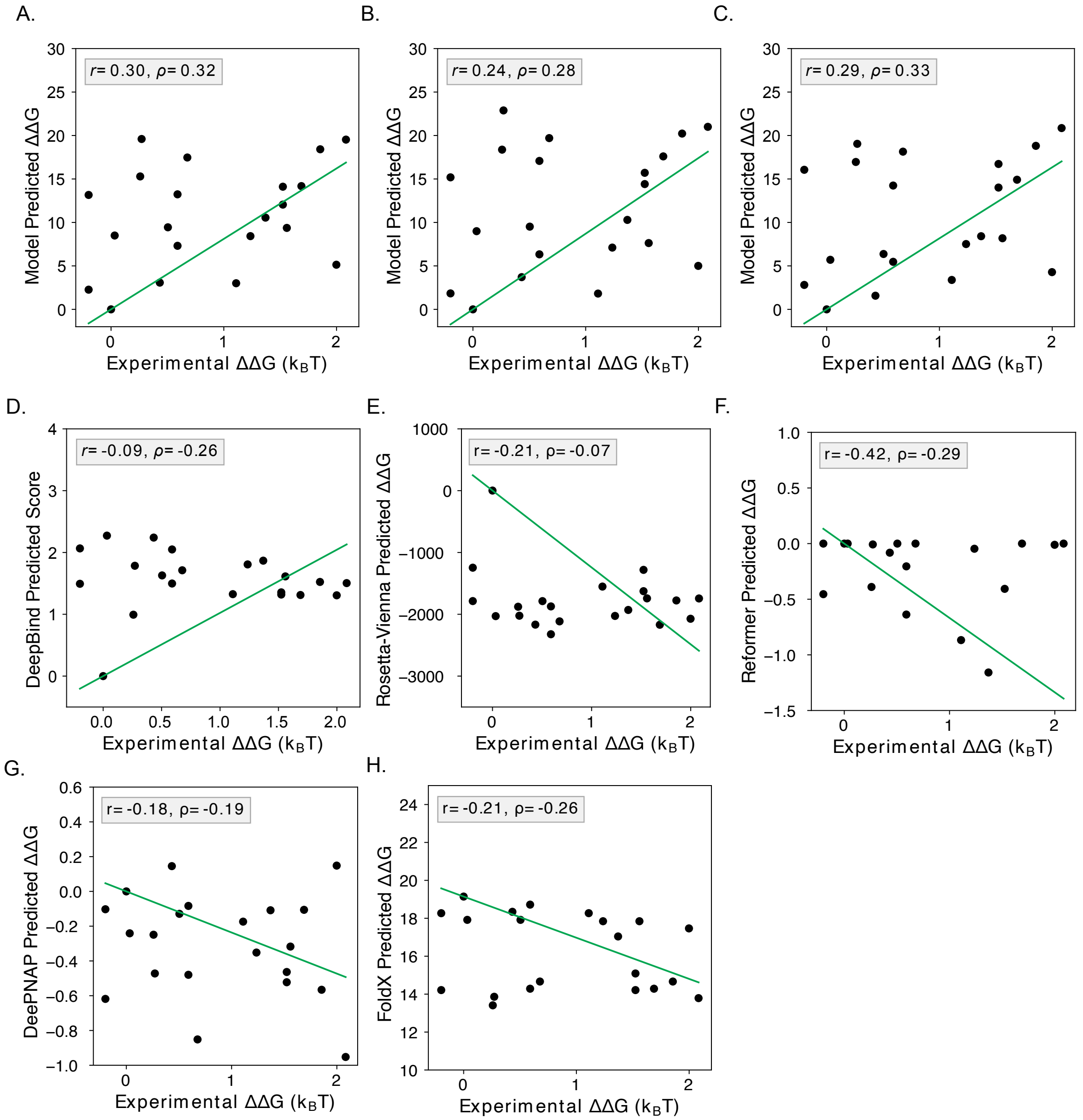


Figure S10: **Comparison of IRIS models trained on different hnRNPKB-motif binding interfaces and other state-of-the-art methods for predicting hnRNPK binding free energies across B-motif variants containing three cytosine patches.** (A) IRIS trained on KH1+KH2-B-motif binding interfaces (same as in Figure 7). (B) IRIS trained on KH1+KH2+RG/RGG-B-motif binding interfaces. (C) IRIS trained on KH1+KH2+RG/RGG+KH3 (full-length hnRNPK)-B-motif binding interfaces. (D) DeepBind.[S1] (E) Rosetta-Vienna.[S2] (F) Reformer.[S7] (G) DeePNAP.[S8] (H) FoldX.[S10]

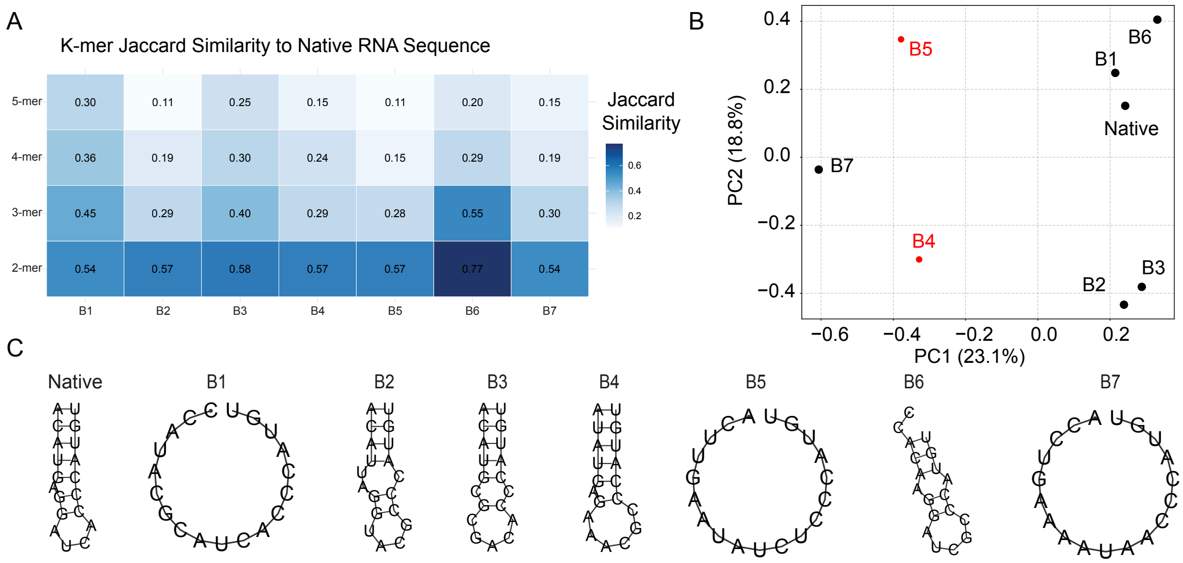


Figure S11: **Sequence diversity and secondary structure of native and outlier RNA sequences (B1-B7) in MS2–RNA complexes.** (A) Sequence similarity was quantified using Jaccard similarity over k-mer sets for $k = 2$ to 5. For each RNA sequence, all overlapping k-mers were generated and compared with those of the 2C4Q motif sequence (ACAUGAGGAUCACCCAUGU). Similarity was defined as the intersection-over-union of the corresponding k-mer sets. The heatmap shows similarity values across increasing k-mer lengths, with darker blue indicating higher similarity. Increasing $k$ increases sensitivity to contiguous motif conservation. Variants B2, B4, B5, and B7 show reduced similarity at higher k values, consistent with disruption of contiguous motif structure relative to the native sequence. (B) Principal component analysis (PCA) of positional 3-mer representations of MS2 RNA sequences (native and B1-B7), projected onto the first two principal components. (C) RNA secondary structure prediction generated using the ViennaRNA package,[S5] showing that B4 most closely resembles the native secondary structure.

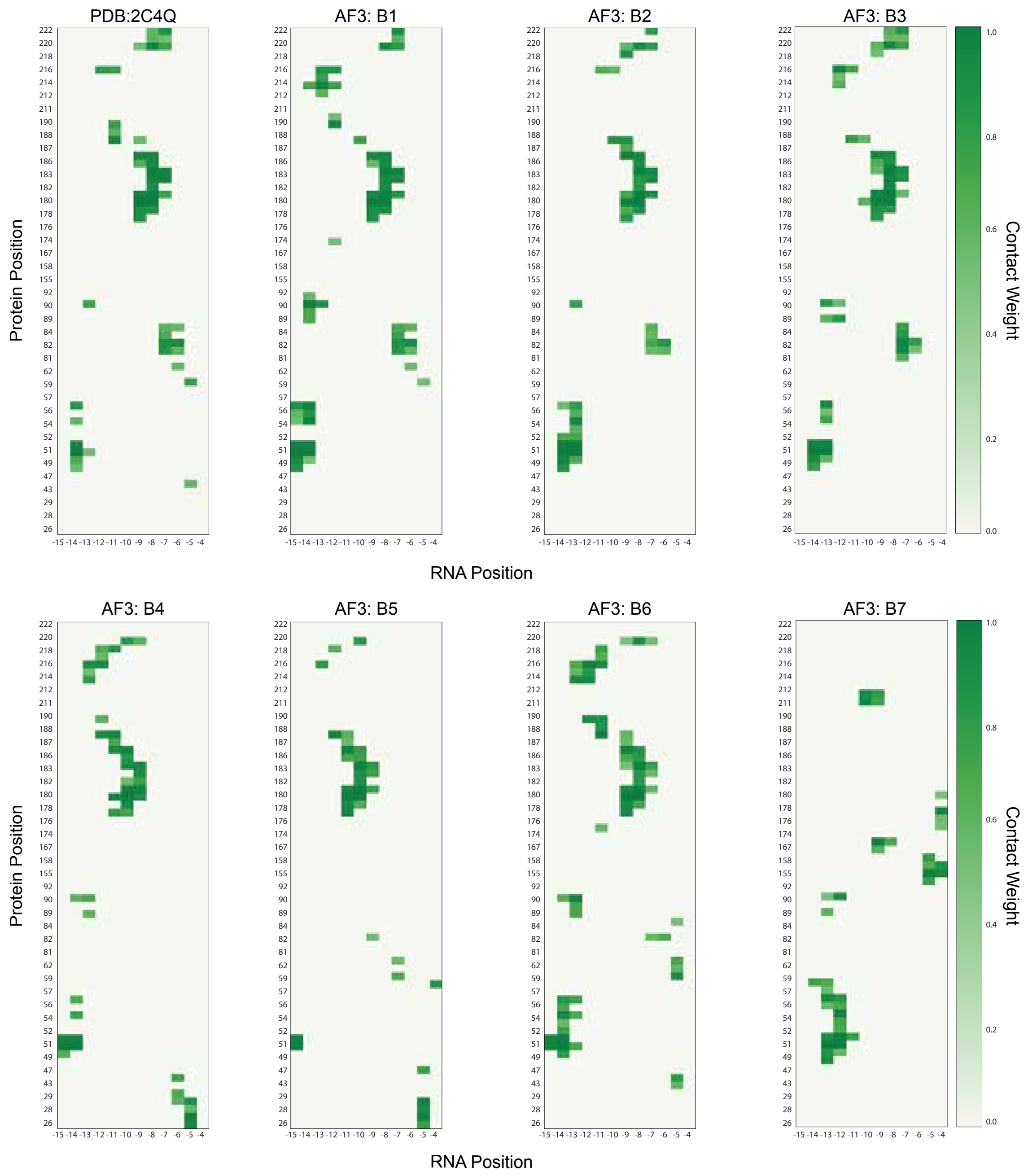


Figure S12: **Aligned protein–RNA contact maps generated using IRIS for the MS2 (2C4Q) complex and high-affinity quadruple-mutant outlier sequences (B1-B7).** Contact weights were computed using a distance cutoff of 0.95 nm and smoothed with a hyperbolic tangent function ($\kappa = 7.0\,\mathrm{nm}^{-1}$). Contact matrices were aligned across all structures, and rows or columns with zero contact across all structures were removed to ensure consistent alignment. Darker green values indicate stronger protein–RNA contacts. Protein positions correspond to C$\alpha$ residue indices, whereas RNA positions correspond to phosphorus atom indices, with residue index offsets applied where necessary (e.g., +1 for 2C4Q). Comparison across MS2 outlier sequences shows that variant B7 exhibits markedly distinct contact patterns relative to other AF3-predicted structures, highlighting sequence-specific perturbations in protein–RNA interactions.

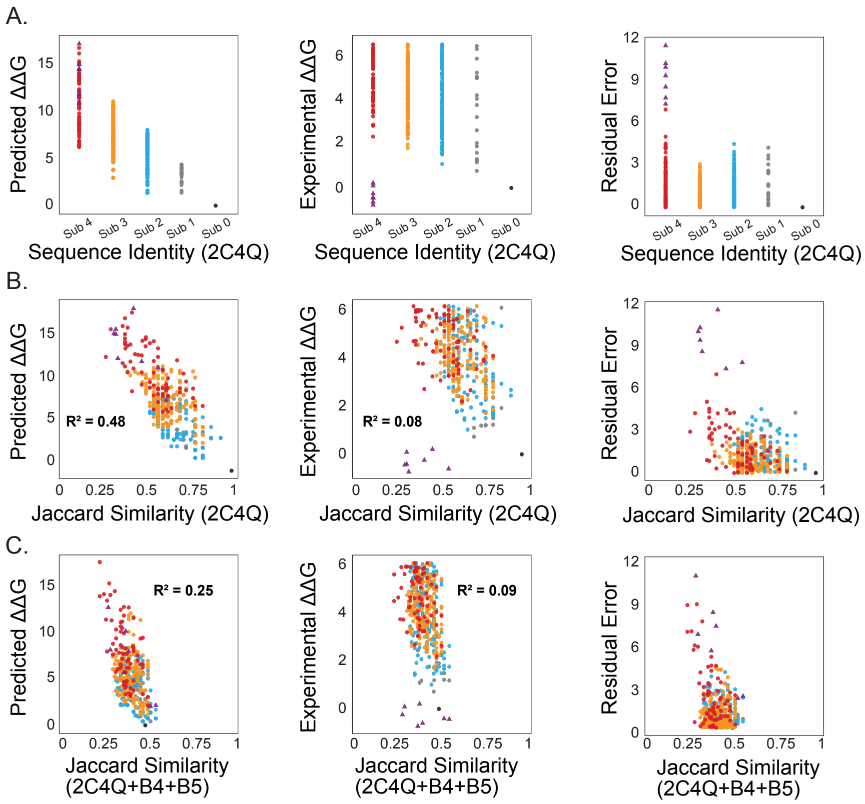


Figure S13: **Effect of sequence similarity and training set composition on IRIS prediction accuracy.** (A) Predicted and experimental $\Delta\Delta G$ values for MS2 RNA variants plotted against sequence identity to the native RNA sequence from the training structure (PDB ID: 2C4Q), separated and colored by mutation class ($0-4$ substitutions). Experimental values exhibit a flag-like distribution with increasing spread at lower sequence identity, whereas predicted values remain approximately linear. Residual errors ($|\Delta\Delta G_{\text{pred}} - \Delta\Delta G_{\text{exp}}|$) show increased dispersion at lower sequence identity, highlighting the models reduced accuracy in this regime. (B) Predicted and experimental $\Delta\Delta G$ values plotted against Jaccard similarity computed from RNA k-mer sets relative to the training sequence (PDB ID: 2C4Q). In contrast to sequence identity, Jaccard similarity produces a more linear relationship for both predicted and experimental values, reducing the flag pattern and better aligning trends across mutation classes. Residual errors increase as similarity decreases, and corresponding $R^2$ values indicate improved linearity compared to sequence identity. (C) Same analysis as in (B) after augmenting the training set with B4 and B5 RNA sequences. Experimental values show the strongest linear relationship in this setting, while predicted values exhibit slightly reduced linearity. Despite this, residual errors are more concentrated, with most variants forming a low-error cluster and a small subset exhibiting larger deviations, indicating improved overall robustness despite localized outliers.

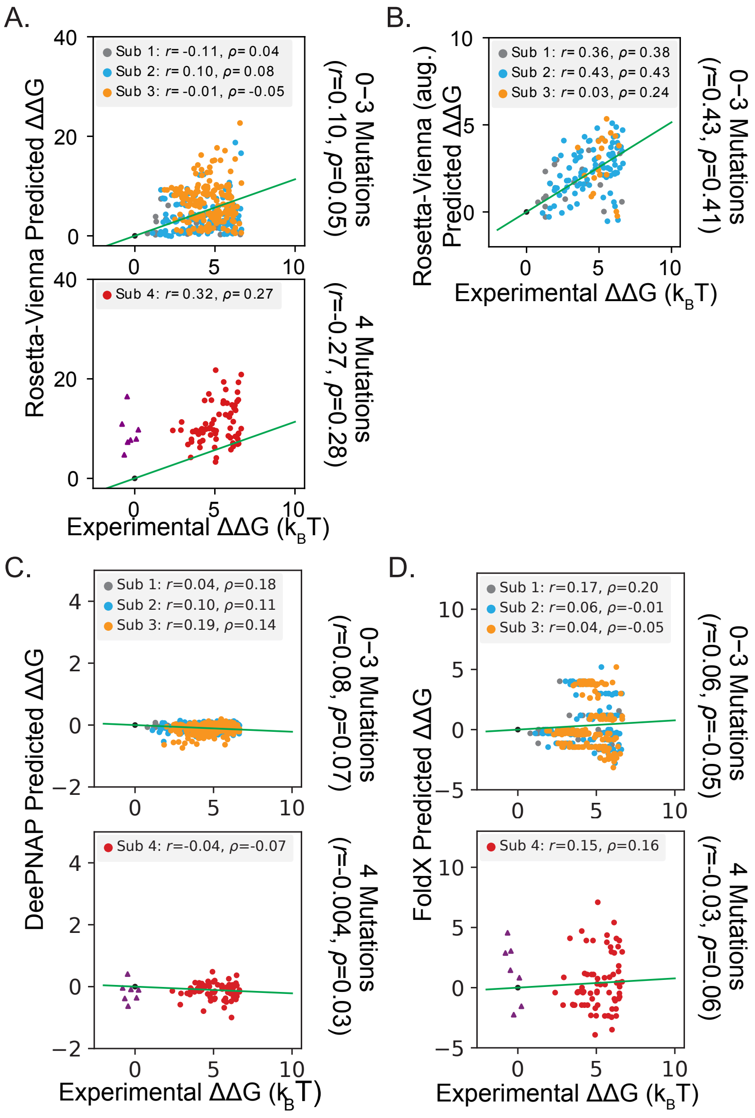


Figure S14: **Correlation plot of predicted versus experimental binding free energies ΔΔG of MS2 coat protein–RNA complexes with single to quadruple RNA mutations.** (A) Predictions from the original, purely structure-based version of the Rosetta-Vienna model.[S2] (B) Predictions from the Rosetta-Vienna version optimized using 50% of the MS2-RNA experimental binding data (Rosetta-Vienna aug.)[S2] After excluding sequences used for optimization and those containing quadruple mutations, a total of 130 test data points remain. (C) Predictions from the DeepNAP model.[S8] (D) Predictions from the FoldX model.[S10]

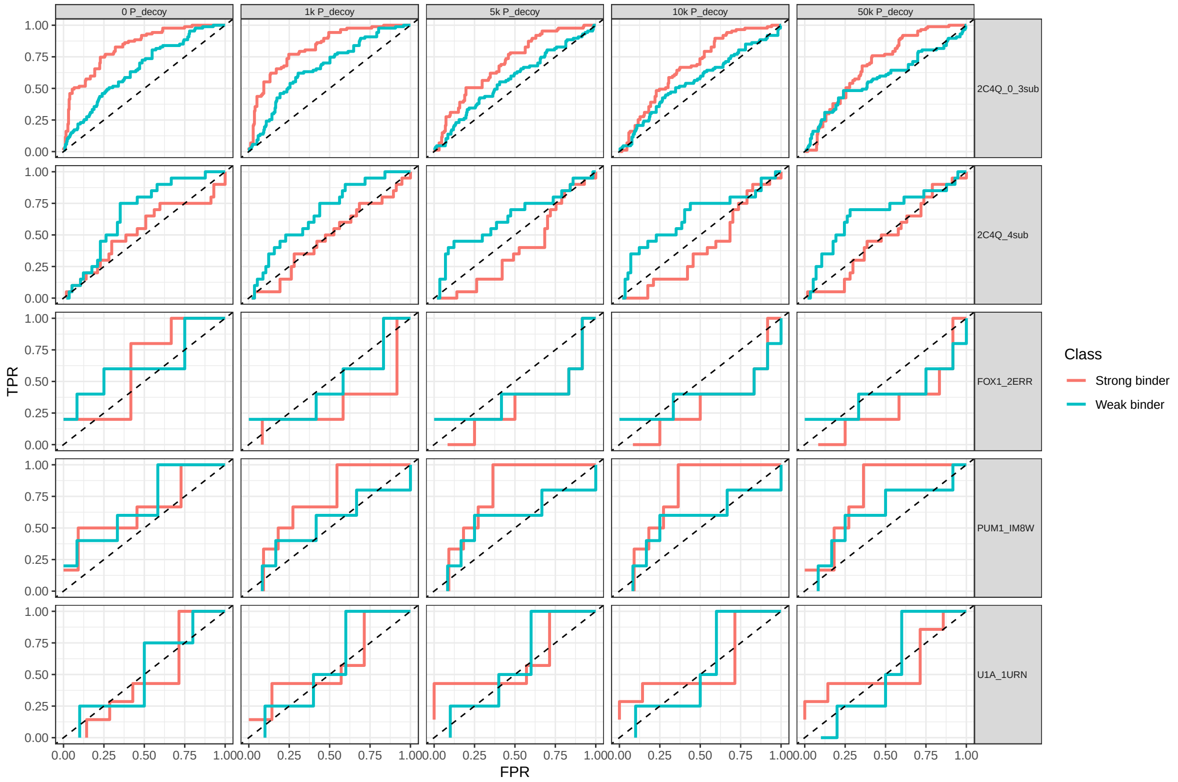


Figure S15: **Rank-based receiver operating characteristic (ROC) analysis for classification of strong and weak protein–RNA binders.** ROC curves were generated independently for each protein–RNA complex and protein-decoy condition using rank-based thresholding of predicted $\Delta\Delta G$ values. Strong binders were defined as variants in the lowest quartile of experimental $\Delta\Delta G$ values, whereas weak binders correspond to the highest quartile. For strong binders, sequences were ranked from lowest to highest predicted $\Delta\Delta G$; for weak binders, the ranking order was reversed. True positive rate (TPR) and false positive rate (FPR) were computed at each rank cutoff. The dashed diagonal line represents random classification performance. Area under the ROC curve (ROC-AUC) was computed using trapezoidal integration.

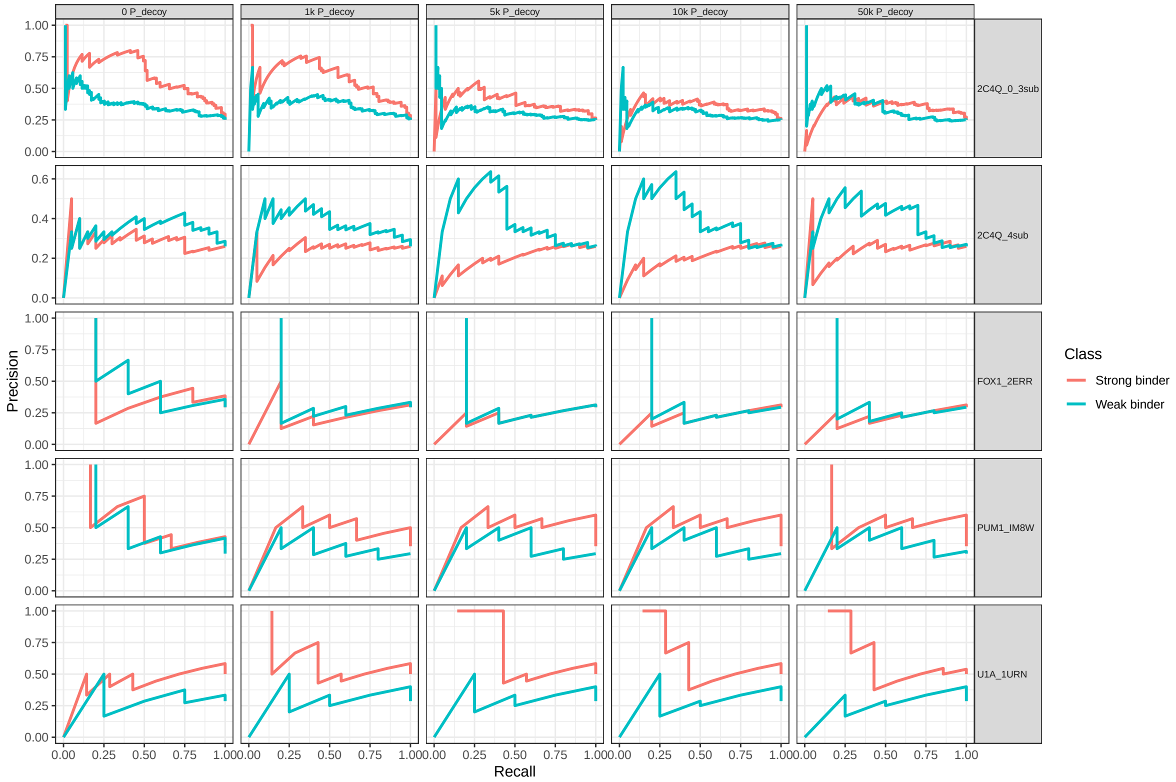


Figure S16: **Rank-based precision–recall (PR) analysis for strong and weak protein–RNA binder classification.** Precision-recall curves corresponding to the ROC analysis in Figure S15. Precision and recall were computed across progressive rank thresholds of predicted $\Delta\Delta G$ values for strong binders (lowest quartile) and weak binders (highest quartile). Curves are shown separately for each protein–RNA complex and protein-decoy condition. Area under the precision–recall curve (PR-AUC) was calculated using trapezoidal integration. PR analysis emphasizes performance under class imbalance and complements ROC-AUC evaluation.

### 10 Supplementary Tables

Table S1: **Parameter ablation analysis for IRIS.** Pearson correlation coefficients ($r$) between predicted and experimental $\Delta\Delta G$ values for the MS2–RNA system under one-parameter-at-a-time ablation. Default parameter settings used in the main text are highlighted in **bold**. RNA binder and non-binder decoy distributions, together with RNA Hamming distance distributions, correspond to those shown in Figure S1 and are described in *Materials and Methods* Section *Decoy Distribution Variants and Sequence Similarity Analysis*; these analyses use 50,000 RNA decoys with no protein decoys. All other ablation settings use 10,000 RNA decoys and no protein decoys unless otherwise specified. The main-text IRIS model uses a uniform Hamming-distance distribution without positional or mutational constraints; therefore, corresponding entries are shown as the default configuration.

| Group | Parameter | Sub 1 | Sub 2 | Sub 3 | Sub 4 | Sub 0–3 |
|---|---|---|---|---|---|---|
| 1k RNA decoys | 10 ev | 0.13 | 0.27 | 0.18 | -0.16 | 0.28 |
| | **25 em** | **0.39** | **0.40** | **0.49** | **-0.09** | **0.46** |
| | 40 em | -0.40 | 0.15 | 0.18 | 0.11 | 0.24 |
| **10k RNA decoys** | 10 em | 0.16 | 0.33 | 0.23 | -0.13 | 0.33 |
| | 15 em | 0.27 | 0.45 | 0.41 | -0.11 | 0.45 |
| | 20 em | 0.52 | 0.53 | 0.58 | -0.07 | 0.55 |
| | **25 em** | **0.50** | **0.48** | **0.56** | **-0.05** | **0.52** |
| | 30 em | 0.28 | 0.38 | 0.43 | -0.03 | 0.41 |
| | 35 em | 0.22 | 0.33 | 0.37 | 0.13 | 0.37 |
| | 40 em | 0.22 | 0.34 | 0.37 | 0.13 | 0.37 |
| 50k RNA decoys | 10 em | 0.15 | 0.33 | 0.23 | -0.13 | 0.33 |
| | **25 em** | **0.49** | **0.47** | **0.56** | **-0.04** | **0.51** |
| | 40 em | 0.49 | 0.32 | 0.37 | 0.13 | 0.36 |
| 100k RNA decoys | 10 em | 0.15 | 0.33 | 0.23 | -0.14 | 0.32 |
| | **25 em** | **0.48** | **0.47** | **0.55** | **-0.04** | **0.51** |
| | 40 em | 0.42 | 0.32 | 0.36 | 0.12 | 0.36 |
| Protein decoys | **0** $P_{\mathrm{decoy}}$ | **0.50** | **0.48** | **0.56** | **-0.05** | **0.52** |
| | 1k $P_{\mathrm{decoy}}$ | 0.36 | 0.49 | 0.50 | -0.09 | 0.51 |
| | 5k $P_{\mathrm{decoy}}$ | 0.11 | 0.25 | 0.18 | -0.11 | 0.27 |
| | 10k $P_{\mathrm{decoy}}$ | 0.12 | 0.21 | 0.15 | -0.09 | 0.24 |
| | 50k $P_{\mathrm{decoy}}$ | 0.19 | 0.27 | 0.17 | -0.01 | 0.27 |
| RNA decoy distribution | **Uniform** | **0.49** | **0.47** | **0.56** | **-0.04** | **0.51** |
| | Binder | 0.52 | 0.49 | 0.58 | -0.05 | 0.53 |
| | Non-binder | 0.53 | 0.50 | 0.59 | -0.05 | 0.53 |
| RNA hamming distribution | Binder | 0.27 | 0.41 | 0.48 | -0.07 | 0.45 |
| | Non-binder | 0.22 | 0.43 | 0.46 | -0.08 | 0.45 |
| Decoy distance cutoff | 0.95 nm | 0.46 | 0.45 | 0.55 | -0.03 | 0.50 |
| | **1.2 nm** | **0.50** | **0.48** | **0.56** | **-0.05** | **0.52** |
| | 1.5 nm | 0.46 | 0.45 | 0.53 | -0.05 | 0.49 |
| Contact distance cutoff | 0.75 nm | 0.24 | 0.36 | 0.36 | 0.06 | 0.38 |
| | 0.85 nm | 0.35 | 0.41 | 0.44 | -0.02 | 0.44 |
| | **0.95 nm** | **0.50** | **0.48** | **0.56** | **-0.05** | **0.52** |
| | 1.05 nm | 0.36 | 0.47 | 0.50 | -0.08 | 0.49 |
| | 1.15 nm | 0.10 | 0.40 | 0.38 | -0.07 | 0.41 |
| Tanh slope $\kappa$ | 6.0 $nm^{-1}$ | 0.53 | 0.50 | 0.57 | -0.05 | 0.53 |
| | **7.0** $\mathbf{nm^{-1}}$ | **0.50** | **0.48** | **0.56** | **-0.05** | **0.52** |
| | 8.0 $nm^{-1}$ | 0.47 | 0.47 | 0.54 | -0.04 | 0.50 |

Table S2: Model performance on 10 biological RNA sequences for hnRNPK across different protein predicted Local Distance Difference Test (pLDDT) thresholds.

| pLDDT | Pearson $r$ | Spearman $\rho$ | ROC-AUC (S) | ROC-AUC (W) | PR-AUC (S) | PR-AUC (W) |
|---|---|---|---|---|---|---|
| 0 | 0.57 | 0.45 | 0.81 | 0.86 | 0.81 | 0.76 |
| 20 | 0.57 | 0.45 | 0.81 | 0.86 | 0.81 | 0.76 |
| 40 | 0.53 | 0.43 | 0.81 | 0.86 | 0.81 | 0.76 |
| 60 | 0.62 | 0.54 | 0.86 | 0.90 | 0.83 | 0.81 |
| 80 | 0.62 | 0.54 | 0.86 | 0.90 | 0.83 | 0.81 |

Table S3: Model performance on 10 biological RNA sequences for hnRNPK across different protein region combinations.

| Protein Region | avg pLDDT | Pearson $r$ | Spearman $\rho$ | ROC-AUC (S) | ROC-AUC (W) | PR-AUC (S) | PR-AUC (W) |
|---|---|---|---|---|---|---|---|
| KH1+KH2 | 78.64 | 0.69 | 0.61 | 0.86 | 0.95 | 0.83 | 0.92 |
| KH1+KH2+RG/RGG | 69.26 | 0.59 | 0.45 | 0.81 | 0.86 | 0.81 | 0.76 |
| Full Protein | 69.35 | 0.57 | 0.45 | 0.81 | 0.86 | 0.81 | 0.76 |

Table S4: **IRIS generalizability across RNA–protein complexes.** Pearson correlations ($r$) between predicted and experimental $\Delta\Delta G$ values across multiple RNA–protein complexes and protein decoy variations. The MS2–RNA dataset is stratified by the number of RNA mutations relative to the wild-type sequence. The other protein–RNA datasets are stratified using mutation distance cutoffs from the protein–RNA interface. Here, RNA_mut (n) and Protein_mut (n) denote the number of RNA and protein mutations, respectively, within each subset, with $N$ representing the total number of variants.

| **MS2-RNA (PDB: 2C4Q)** | | | | | | | | |
|---|---|---|---|---|---|---|---|---|
| **Parameter** | **Sample size (N)** | **RNA_mut (n)** | **Protein_mut (n)** | **0 P_decoy** | **1k P_decoy** | **5k P_decoy** | **10k P_decoy** | **50k P_decoy** |
| 1_mut | 21 | 21 | 0 | 0.50 | 0.36 | 0.11 | 0.12 | 0.19 |
| 2_mut | 132 | 132 | 0 | 0.48 | 0.49 | 0.25 | 0.21 | 0.27 |
| 3_mut | 192 | 192 | 0 | 0.56 | 0.50 | 0.18 | 0.15 | 0.17 |
| 4_mut | 77 | 77 | 0 | -0.05 | -0.09 | -0.11 | -0.09 | -0.01 |
| 0-3_mut | 346 | 346 | 0 | 0.52 | 0.51 | 0.27 | 0.24 | 0.27 |
| **U1A-RNA (PDB: 1URN)** | | | | | | | | |
| **Parameter** | **Sample size (N)** | **RNA_mut (n)** | **Protein_mut (n)** | **0 P_decoy** | **1k P_decoy** | **5k P_decoy** | **10k P_decoy** | **50k P_decoy** |
| 0.3 nm | 3 | 0 | 3 | 0.86 | 0.96 | 0.95 | 0.93 | 0.88 |
| 0.4 nm | 13 | 0 | 13 | -0.05 | 0.05 | 0.04 | -0.02 | -0.08 |
| 0.5 nm | 13 | 0 | 13 | -0.05 | 0.05 | 0.04 | -0.02 | -0.08 |
| 0.7 nm | 14 | 0 | 14 | -0.04 | 0.05 | 0.04 | -0.01 | -0.08 |
| 0.95 nm | 17 | 0 | 17 | 0.01 | 0.10 | 0.09 | 0.04 | -0.01 |
| **FOX1-RNA (PDB: 2ERR)** | | | | | | | | |
| **Parameter** | **Sample size (N)** | **RNA_mut (n)** | **Protein_mut (n)** | **0 P_decoy** | **1k P_decoy** | **5k P_decoy** | **10k P_decoy** | **50k P_decoy** |
| 0.3 nm | 6 | 6 | 0 | 0.53 | 0.18 | 0.34 | 0.42 | 0.49 |
| 0.4 nm | 14 | 6 | 8 | 0.38 | -0.21 | -0.10 | -0.02 | 0.05 |
| 0.5 nm | 16 | 6 | 10 | 0.13 | -0.19 | -0.10 | -0.04 | 0.02 |
| 0.7 nm | 16 | 6 | 10 | 0.13 | -0.19 | -0.10 | -0.04 | 0.02 |
| 0.95 nm | 16 | 6 | 10 | 0.13 | -0.19 | -0.10 | -0.04 | 0.02 |
| **PUM1-RNA (PDB: IM8W)** | | | | | | | | |
| **Parameter** | **Sample size (N)** | **RNA_mut (n)** | **Protein_mut (n)** | **0 P_decoy** | **1k P_decoy** | **5k P_decoy** | **10k P_decoy** | **50k P_decoy** |
| 0.3 nm | 8 | 7 | 3 | 0.72 | 0.81 | 0.81 | 0.81 | 0.80 |
| 0.4 nm | 16 | 11 | 11 | 0.34 | 0.33 | 0.42 | 0.44 | 0.44 |
| 0.5 nm | 16 | 11 | 11 | 0.34 | 0.33 | 0.42 | 0.44 | 0.44 |
| 0.7 nm | 16 | 11 | 11 | 0.34 | 0.33 | 0.42 | 0.44 | 0.44 |
| 0.95 nm | 16 | 11 | 11 | 0.34 | 0.33 | 0.42 | 0.44 | 0.44 |
| **SRP-RNA (PDB: 1HQ1)** | | | | | | | | |
| **Parameter** | **Sample size (N)** | **RNA_mut (n)** | **Protein_mut (n)** | **0 P_decoy** | **1k P_decoy** | **5k P_decoy** | **10k P_decoy** | **50k P_decoy** |
| 0.3 nm | 4 | 4 | 0 | 0.59 | 0.39 | 0.37 | 0.32 | 0.28 |
| 0.4 nm | 5 | 5 | 0 | 0.37 | 0.31 | 0.36 | 0.33 | 0.29 |
| 0.5 nm | 6 | 6 | 0 | 0.33 | 0.28 | 0.39 | 0.39 | 0.36 |
| 0.7 nm | 9 | 9 | 0 | -0.13 | -0.19 | -0.12 | -0.09 | -0.08 |
| 0.95 nm | 11 | 11 | 0 | 0.02 | -0.03 | 0.03 | 0.04 | 0.03 |

Table S5: **Prediction error, correlation, and classification performance of IRIS across RNA–protein complexes.** Variants with mutations occurring within 0.5 nm of the protein–RNA interface were retained. Pearson ($r$) and Spearman ($\rho$) correlations were computed directly between predicted and experimental $\Delta\Delta G$ values. Mean absolute error (MAE) and root-mean-square error (RMSE) were calculated relative to the through-origin best-fit line, with 95% confidence intervals obtained via bootstrap resampling. Strong and weak binders were defined independently within each complex as the lowest and highest experimental $\Delta\Delta G$ quartiles, respectively. ROC and precision–recall (PR) curves were generated using rank-based thresholding on predicted $\Delta\Delta G$ values, with AUC values reported separately for strong and weak binders (ROC / PR). Rows in **bold** indicate the best protein decoy parameter used for future analysis.

| **MS2-RNA (PDB: 2C4Q) (0–3 sub)** | | | | | | |
|---|---|---|---|---|---|---|
| **Protein Decoys** | **RMSE [CI]** | **MAE [CI]** | **Pearson $r$** | **Spearman $\rho$** | **ROC-AUC (S/W)** | **PR-AUC (S/W)** |
| **0 P_decoy** | **1.65 [1.42–1.87]** | **1.57 [1.35–1.78]** | **0.52** | **0.49** | **0.83/0.65** | **0.61/0.38** |
| 1k P_decoy | 0.48 [0.29–0.67] | 0.46 [0.28–0.64] | 0.51 | 0.49 | 0.82/0.67 | 0.58/0.36 |
| 5k P_decoy | 0.18 [0.01–0.41] | 0.17 [0.01–0.39] | 0.27 | 0.25 | 0.70/0.57 | 0.38/0.31 |
| 10k P_decoy | 0.17 [0.01–0.42] | 0.16 [0.01–0.40] | 0.24 | 0.23 | 0.68/0.56 | 0.36/0.29 |
| 50k P_decoy | 1.75 [1.36–2.15] | 1.67 [1.29–2.05] | 0.27 | 0.25 | 0.69/0.58 | 0.35/0.34 |
| **MS2-RNA (PDB: 2C4Q) (4 sub)** | | | | | | |
| **Protein Decoys** | **RMSE [CI]** | **MAE [CI]** | **Pearson $r$** | **Spearman $\rho$** | **ROC-AUC (S/W)** | **PR-AUC (S/W)** |
| **0 P_decoy** | **4.98 [4.37–5.57]** | **4.68 [4.04–5.31]** | **-0.05** | **0.22** | **0.53/0.67** | **0.28/0.34** |
| 1k P_decoy | 2.69 [2.19–3.19] | 2.53 [2.02–3.04] | -0.09 | 0.15 | 0.48/0.68 | 0.24/0.37 |
| 5k P_decoy | 1.12 [0.59–1.66] | 1.06 [0.55–1.57] | -0.11 | 0.01 | 0.39/0.63 | 0.20/0.39 |
| 10k P_decoy | 1.21 [0.63–1.86] | 1.14 [0.60–1.75] | -0.09 | 0.02 | 0.40/0.64 | 0.20/0.39 |
| 50k P_decoy | 4.25 [3.17–5.34] | 4.00 [2.97–5.02] | -0.01 | 0.11 | 0.48/0.66 | 0.24/0.38 |
| **FOX1-RNA (PDB: 2ERR)** | | | | | | |
| **Protein Decoys** | **RMSE [CI]** | **MAE [CI]** | **Pearson $r$** | **Spearman $\rho$** | **ROC-AUC (S/W)** | **PR-AUC (S/W)** |
| **0 P_decoy** | **0.50 [0.03–1.54]** | **0.43 [0.03–1.37]** | **0.13** | **0.12** | **0.61/0.63** | **0.26/0.33** |
| 1k P_decoy | 2.45 [1.44–3.33] | 2.13 [1.21–3.03] | -0.19 | -0.13 | 0.32/0.47 | 0.23/0.21 |
| 5k P_decoy | 2.85 [1.64–3.90] | 2.47 [1.37–3.51] | -0.10 | -0.17 | 0.32/0.38 | 0.21/0.19 |
| 10k P_decoy | 2.85 [1.51–3.88] | 2.48 [1.26–3.50] | -0.04 | -0.17 | 0.32/0.38 | 0.21/0.20 |
| 50k P_decoy | 2.37 [0.35–3.88] | 2.06 [0.30–3.46] | 0.02 | -0.18 | 0.30/0.40 | 0.21/0.20 |
| **PUM1-RNA (PDB: IM8W)** | | | | | | |
| **Protein Decoys** | **RMSE [CI]** | **MAE [CI]** | **Pearson $r$** | **Spearman $\rho$** | **ROC-AUC (S/W)** | **PR-AUC (S/W)** |
| 0 P_decoy | 20.55 [3.60–38.12] | 16.82 [2.96–32.12] | 0.34 | 0.39 | 0.65/0.68 | 0.41/0.34 |
| 1k P_decoy | 0.39 [0.03–1.65] | 0.32 [0.02–1.40] | 0.33 | 0.20 | 0.71/0.53 | 0.47/0.31 |
| **5k P_decoy** | **0.47 [0.03–1.92]** | **0.38 [0.03–1.59]** | **0.42** | **0.29** | **0.77/0.57** | **0.50/0.34** |
| 10k P_decoy | 0.33 [0.03–1.81] | 0.27 [0.02–1.50] | 0.44 | 0.29 | 0.77/0.57 | 0.50/0.34 |
| 50k P_decoy | 0.85 [0.06–3.42] | 0.69 [0.05–2.81] | 0.44 | 0.35 | 0.77/0.62 | 0.43/0.35 |
| **SRP-RNA (PDB: 1HQ1)** | | | | | | |
| **Protein Decoys** | **RMSE [CI]** | **MAE [CI]** | **Pearson $r$** | **Spearman $\rho$** | **ROC-AUC (S/W)** | **PR-AUC (S/W)** |
| 0 P_decoy | 0.81 [0.06–2.31] | 0.58 [0.04–1.86] | 0.33 | 0.11 | 0.50/0.50 | 0.11/0.26 |
| 1k P_decoy | 0.08 [0.01–1.64] | 0.06 [0.01–1.22] | 0.28 | 0.07 | 0.70/0.50 | 0.16/0.26 |
| **5k P_decoy** | **0.44 [0.05–1.29]** | **0.32 [0.03–1.12]** | **0.39** | **0.32** | **0.80/0.50** | **0.21/0.25** |
| 10k P_decoy | 0.43 [0.04–1.29] | 0.31 [0.03–1.09] | 0.39 | 0.32 | 0.80/0.50 | 0.21/0.25 |
| 50k P_decoy | 0.02 [0.02–1.63] | 0.01 [0.01–1.30] | 0.36 | 0.39 | 0.80/0.60 | 0.21/0.29 |
| **U1A-RNA (PDB: 1URN)** | | | | | | |
| **Protein Decoys** | **RMSE [CI]** | **MAE [CI]** | **Pearson $r$** | **Spearman $\rho$** | **ROC-AUC (S/W)** | **PR-AUC (S/W)** |
| 0 P_decoy | 2.45 [0.10–7.47] | 1.82 [0.08–6.14] | -0.05 | -0.01 | 0.47/0.53 | 0.44/0.28 |
| 1k P_decoy | 0.33 [0.02–2.35] | 0.24 [0.02–1.89] | 0.05 | 0.11 | 0.57/0.58 | 0.47/0.29 |
| **5k P_decoy** | **0.22 [0.03–1.36]** | **0.16 [0.02–1.10]** | **0.04** | **0.16** | **0.61/0.58** | **0.57/0.29** |
| 10k P_decoy | 0.31 [0.03–1.31] | 0.23 [0.02–1.02] | -0.02 | 0.08 | 0.57/0.55 | 0.53/0.28 |
| 50k P_decoy | 0.15 [0.03–2.09] | 0.11 [0.02–1.73] | -0.08 | 0.05 | 0.55/0.53 | 0.52/0.26 |

Table S6: **Prediction error, correlation, and classification performance on Fox-1 under varying interface distance cutoffs (0.3–0.5 nm).** Metrics were computed directly between predicted and experimental $\Delta\Delta G$ values. Mean absolute error (MAE) and root-mean-square error (RMSE) were calculated relative to the best-fit line through the origin, with 95% confidence intervals obtained via bootstrap resampling. Pearson ($r$) and Spearman ($\rho$) correlations were computed directly between predicted and experimental values. Strong (S) and weak (W) binders were defined as the lowest and highest experimental $\Delta\Delta G$ quartiles, respectively, and ROC and precision–recall (PR)-AUC values were calculated using quartile-based binarization. The top-performing results for each metric are highlighted in **bold**. IRIS results were obtained using 0 protein decoys and 10,000 RNA decoys, corresponding to the best-performing configuration highlighted in Table S5.

| **Fox-1 (mutation distance cutoff = 0.3 nm)** | | | | | | |
|---|---|---|---|---|---|---|
| **Method** | **RMSE [CI]** | **MAE [CI]** | **Pearson $r$** | **Spearman $\rho$** | **ROC-AUC (S/W)** | **PR-AUC (S/W)** |
| IRIS | **0.46 [0.02–1.78]** | **0.37 [0.02–1.55]** | 0.53 | 0.11 | 0.80/0.25 | 0.23/0.18 |
| DeepBind | 2.61 [0.93–3.54] | 2.09 [0.93–3.28] | -0.49 | -0.32 | 0.60/0.30 | 0.25/0.16 |
| Rosetta-Vienna | 4.50 [0.54–21.19] | 3.61 [0.54–17.76] | -0.37 | 0.29 | 0.60/0.60 | 0.21/0.25 |
| Rosetta-Vienna (aug.) | 1.25 [0.16–2.17] | 1.00 [0.11–1.92] | 0.25 | 0.29 | 0.50/0.65 | **0.20/0.31** |
| DeePNAP | 2.73 [0.92–3.77] | 2.19 [0.92–3.44] | -0.21 | -0.07 | 0.35/0.45 | 0.21/0.27 |
| FoldX | 8.08 [6.48–9.51] | 6.47 [4.26–8.74] | **0.96** | **0.85** | **0.80/0.90** | 0.00/0.32 |
| **Fox-1 (mutation distance cutoff = 0.4 nm)** | | | | | | |
| IRIS | **0.28 [0.02–1.14]** | **0.24 [0.02–0.94]** | 0.38 | 0.33 | 0.72/0.72 | 0.29/0.29 |
| DeepBind | 3.25 [2.53–3.93] | 2.82 [2.02–3.68] | -0.46 | -0.38 | 0.28/0.27 | 0.16/0.15 |
| Rosetta-Vienna | 1.10 [0.04–5.41] | 0.95 [0.03–4.45] | -0.34 | 0.44 | 0.70/0.75 | 0.31/0.31 |
| Rosetta-Vienna (aug.) | 1.36 [0.79–1.90] | 1.18 [0.64–1.72] | **0.55** | **0.59** | 0.78/0.75 | **0.53/0.35** |
| DeePNAP | 3.25 [1.94–3.92] | 2.82 [1.94–3.65] | 0.07 | 0.12 | 0.70/0.56 | 0.29/0.29 |
| FoldX | 9.62 [8.34–10.84] | 8.36 [6.53–10.16] | 0.59 | 0.53 | **0.84/0.76** | 0.27/0.29 |
| **Fox-1 (mutation distance cutoff = 0.5 nm)** | | | | | | |
| IRIS | 0.50 [0.02–1.48] | 0.43 [0.02–1.31] | 0.13 | 0.12 | 0.61/0.63 | 0.27/0.29 |
| DeepBind | 3.62 [2.71–4.34] | 3.15 [2.22–4.02] | -0.45 | -0.38 | 0.20/0.30 | 0.16/0.16 |
| Rosetta-Vienna | **0.21 [0.04–3.31]** | **0.19 [0.04–2.79]** | -0.35 | 0.44 | 0.54/0.77 | 0.30/0.37 |
| Rosetta-Vienna (aug.) | 1.79 [1.05–2.51] | 1.55 [0.88–2.25] | 0.49 | **0.55** | 0.59/0.77 | **0.44/0.44** |
| DeePNAP | 3.65 [2.84–4.32] | 3.18 [2.28–3.99] | -0.01 | 0.06 | 0.52/0.58 | 0.31/0.31 |
| FoldX | 10.09 [8.83–11.27] | 8.77 [7.08–10.50] | **0.47** | 0.43 | **0.92/0.73** | 0.33/0.33 |

Table S7: **Prediction error, correlation, and classification performance on PUM1 under varying interface distance cutoffs (0.3–0.4 nm).** Metrics were computed directly between predicted and experimental $\Delta\Delta G$ values. Mean absolute error (MAE) and root-mean-square error (RMSE) were calculated relative to the best-fit line through the origin, with 95% confidence intervals obtained via bootstrap resampling. Pearson ($r$) and Spearman ($\rho$) correlations were computed directly between predicted and experimental values. Strong (S) and weak (W) binders were defined as the lowest and highest experimental $\Delta\Delta G$ quartiles, respectively, and ROC and precision–recall (PR)-AUC values were calculated using quartile-based binarization. The top-performing results for each metric are highlighted in **bold**. IRIS results were obtained using 5,000 protein decoys and 10,000 RNA decoys, corresponding to the best-performing configuration highlighted in Table S5.

| **PUM1 (mutation distance cutoff = 0.3 nm)** | | | | | | |
|---|---|---|---|---|---|---|
| **Method** | **RMSE [CI]** | **MAE [CI]** | **Pearson $r$** | **Spearman $\rho$** | **ROC-AUC (S/W)** | **PR-AUC (S/W)** |
| IRIS | **0.11 [0.01–1.41]** | **0.09 [0.01–1.24]** | 0.81 | 0.67 | **0.64/0.92** | 0.27/0.39 |
| DeepBind | 2.48 [1.57–3.38] | 2.03 [1.17–3.03] | 0.44 | 0.48 | 0.56/0.78 | 0.26/0.40 |
| Rosetta-Vienna | 14.67 [6.69–20.79] | 11.99 [5.14–18.38] | 0.05 | 0.23 | 0.69/0.64 | 0.37/0.32 |
| Rosetta-Vienna (aug) | 0.80 [0.35–1.13] | 0.65 [0.30–0.99] | **0.82** | **0.70** | 0.69/0.83 | **0.37/0.43** |
| DeePNAP | 2.31 [1.39–3.12] | 1.89 [1.01–2.78] | 0.70 | 0.67 | 0.72/0.83 | 0.30/0.43 |
| FoldX | 5.51 [4.20–6.91] | 4.51 [3.04–6.43] | 0.05 | 0.04 | 0.42/0.42 | 0.28/0.26 |
| **PUM1 (mutation distance cutoff = 0.4 nm)** | | | | | | |
| **Method** | **RMSE [CI]** | **MAE [CI]** | **Pearson $r$** | **Spearman $\rho$** | **ROC-AUC (S/W)** | **PR-AUC (S/W)** |
| IRIS | 0.47 [0.03–1.86] | 0.38 [0.02–1.56] | 0.42 | 0.29 | 0.78/0.57 | 0.49/0.33 |
| DeepBind | 2.53 [1.78–3.19] | 2.07 [1.41–2.80] | -0.08 | 0.12 | 0.54/0.51 | 0.21/0.27 |
| Rosetta-Vienna | 8.39 [2.79–13.54] | 6.87 [2.20–11.57] | -0.04 | 0.11 | 0.60/0.53 | 0.33/0.23 |
| Rosetta-Vienna (aug) | **0.40 [0.03–0.85]** | **0.32 [0.03–0.71]** | **0.76** | **0.75** | **0.92/0.81** | **0.70/0.44** |
| DeePNAP | 2.27 [1.25–2.95] | 1.86 [1.25–2.55] | 0.29 | 0.35 | 0.70/0.63 | 0.45/0.29 |
| FoldX | 5.36 [3.03–5.68] | 4.39 [3.03–5.68] | -0.12 | -0.17 | 0.34/0.36 | 0.26/0.21 |

Table S8: **Prediction error, correlation, and classification performance of different methods on SRP under varying interface distance cutoffs (0.3–0.95 nm).** Metrics include Pearson ($r$) and Spearman ($\rho$) correlations between predicted and experimental $\Delta\Delta G$ values. Mean absolute error (MAE) and root-mean-square error (RMSE) were calculated relative to the best-fit line through the origin, with 95% bootstrap confidence intervals. Strong (S) and weak (W) binders were defined as the lowest and highest experimental $\Delta\Delta G$ quartiles, and ROC and precision–recall (PR)-AUC values were computed using quartile-based binarization. The top-performing results for each metric are highlighted in **bold**. IRIS predictions were generated using 5,000 protein decoys and 10,000 RNA decoys, corresponding to the best-performing configuration highlighted in Table S5.

| Method | RMSE [CI] | MAE [CI] | Pearson $r$ | Spearman $\rho$ | ROC-AUC (S/W) | PR-AUC (S/W) |
|---|---|---|---|---|---|---|
| **SRP (mutation distance cutoff = 0.3 nm)** | | | | | | |
| IRIS | 0.64 [0.06–1.62] | 0.49 [0.05–1.43] | 0.38 | 0.10 | 0.50/0.33 | 0.29/0.16 |
| DeepBind | 2.98 [1.37–4.23] | 2.30 [0.77–3.82] | 0.10 | 0.45 | 0.50/0.25 | **0.83/0.38** |
| Rosetta-Vienna | 2.76 [1.47–5.01] | 2.13 [0.97–4.48] | 0.46 | **0.80** | **0.83/1.00** | 0.29/0.50 |
| Rosetta-Vienna (aug) | **0.20 [0.03–2.15]** | **0.16 [0.02–1.85]** | 0.45 | **0.80** | **0.83/1.00** | 0.29/0.50 |
| DeePNAP | 2.65 [1.05–3.88] | 2.05 [0.55–3.51] | -0.61 | -0.10 | 0.67/0.50 | 0.42/0.16 |
| FoldX | 8.65 [5.97–10.75] | 6.68 [3.25–10.27] | **0.90** | 0.70 | **0.83/1.00** | 0.29/0.50 |
| **SRP (mutation distance cutoff = 0.4 nm)** | | | | | | |
| IRIS | 0.52 [0.06–1.62] | 0.39 [0.05–1.43] | 0.37 | 0.31 | 0.88/0.38 | 0.29/0.25 |
| DeepBind | 2.74 [1.30–3.96] | 2.02 [0.81–3.58] | 0.04 | 0.34 | 0.50/0.20 | **0.81/0.35** |
| Rosetta-Vienna | 2.65 [1.41–4.77] | 1.96 [0.81–4.09] | 0.54 | 0.83 | **1.00/1.00** | 0.50/0.50 |
| Rosetta-Vienna (aug) | **0.19 [0.03–1.57]** | **0.14 [0.03–1.57]** | 0.53 | **0.89** | **1.00/1.00** | 0.50/0.50 |
| DeePNAP | 2.43 [1.01–3.66] | 1.79 [0.64–3.29] | -0.37 | -0.03 | 0.63/0.50 | 0.33/0.13 |
| FoldX | 8.20 [nan–nan] | 6.05 [nan–nan] | **0.87** | 0.54 | 0.50/1.00 | 0.29/0.50 |
| **SRP (mutation distance cutoff = 0.5 nm)** | | | | | | |
| IRIS | 0.44 [0.04–1.34] | 0.32 [0.03–1.15] | 0.39 | 0.32 | 0.80/0.50 | 0.25/0.25 |
| DeepBind | 2.56 [1.21–3.73] | 1.83 [0.82–3.19] | -0.05 | 0.20 | 0.60/0.33 | 0.20/0.33 |
| Rosetta-Vienna | 2.65 [1.55–4.64] | 1.90 [0.90–3.82] | 0.50 | **0.79** | **1.00/1.00** | **0.50/0.50** |
| Rosetta-Vienna (aug) | **0.10 [0.03–1.52]** | **0.07 [0.02–1.52]** | 0.48 | 0.75 | 1.00/0.90 | 0.50/0.29 |
| DeePNAP | 2.26 [0.91–3.30] | 1.62 [0.54–2.77] | -0.43 | 0.04 | 0.70/0.40 | 0.33/0.24 |
| FoldX | 7.92 [6.10–10.06] | 5.67 [3.48–9.06] | **0.88** | 0.43 | 0.40/1.00 | 0.23/0.50 |
| **SRP (mutation distance cutoff = 0.7 nm)** | | | | | | |
| IRIS | 1.18 [0.06–2.04] | 0.88 [0.05–1.82] | -0.12 | -0.01 | 0.41/0.38 | 0.18/0.23 |
| DeepBind | 2.79 [1.57–3.80] | 2.08 [1.11–3.25] | -0.33 | -0.26 | 0.36/0.50 | 0.19/0.31 |
| Rosetta-Vienna | 3.29 [2.08–4.68] | 2.45 [1.35–3.95] | **0.68** | **0.73** | 0.86/0.95 | 0.48/0.57 |
| Rosetta-Vienna (aug) | **0.13 [0.02–1.39]** | **0.10 [0.02–1.13]** | 0.61 | 0.61 | **0.91/0.91** | **0.52/0.52** |
| DeePNAP | 2.45 [1.13–3.40] | 1.82 [0.82–2.91] | -0.26 | -0.12 | 0.57/0.43 | 0.29/0.26 |
| FoldX | 8.82 [7.07–10.70] | 6.56 [4.51–9.18] | 0.63 | 0.48 | 0.62/0.95 | 0.22/0.57 |
| **SRP (mutation distance cutoff = 0.95–0.96 nm)** | | | | | | |
| IRIS | 1.14 [0.05–2.04] | 0.82 [0.05–1.73] | 0.03 | 0.12 | 0.59/0.48 | 0.11/0.22 |
| DeepBind | 2.61 [1.50–3.48] | 1.89 [1.07–2.89] | 0.22 | -0.02 | 0.59/0.56 | 0.17/0.29 |
| Rosetta-Vienna | 3.23 [2.17–4.39] | 2.34 [1.41–3.64] | **0.71** | **0.76** | **1.00/0.96** | **0.67/0.57** |
| Rosetta-Vienna (aug) | **0.20 [0.02–1.33]** | **0.15 [0.02–1.05]** | 0.66 | 0.70 | 1.00/0.89 | 0.67/0.48 |
| DeePNAP | 2.27 [1.17–3.21] | 1.65 [0.81–2.62] | -0.18 | 0.01 | 0.69/0.43 | 0.28/0.20 |
| FoldX | 8.53 [6.81–10.27] | 6.18 [4.21–8.64] | 0.64 | 0.55 | 0.52/0.96 | 0.22/0.50 |

Table S9: **Prediction error, correlation, and classification performance on U1A under varying interface distance cutoffs (0.3–0.96 nm).** Metrics were computed directly between predicted and experimental $\Delta\Delta G$ values. Mean absolute error (MAE) and root-mean-square error (RMSE) were calculated relative to the best-fit line through the origin, with 95% confidence intervals obtained via bootstrap resampling. Pearson ($r$) and Spearman ($\rho$) correlations were computed directly between predicted and experimental values. Strong (S) and weak (W) binders were defined as the lowest and highest experimental $\Delta\Delta G$ quartiles, respectively, and ROC and precision–recall (PR)-AUC values were calculated using quartile-based binarization. The top-performing results for each metric are highlighted in **bold** IRIS results were obtained using 5,000 protein decoys and 10,000 RNA decoys, corresponding to the best-performing configuration highlighted in Table S5.

| **U1A (mutation distance cutoff = 0.3 nm)** | | | | | | |
|---|---|---|---|---|---|---|
| **Method** | **RMSE [CI]** | **MAE [CI]** | **Pearson $r$** | **Spearman $\rho$** | **ROC-AUC (S/W)** | **PR-AUC (S/W)** |
| IRIS | **0.25 [nan–nan]** | **0.19 [nan–nan]** | **0.95** | **0.80** | 0.50/1.00 | 0.17/0.00 |
| Rosetta-Vienna | 0.79 [nan–nan] | 0.59 [nan–nan] | 0.30 | 0.40 | 1.00/0.50 | 0.00/0.17 |
| Rosetta-Vienna (aug) | 0.60 [nan–nan] | 0.45 [nan–nan] | **0.95** | **0.80** | 1.00/0.00 | 1.00/0.00 |
| DeePNAP | 0.90 [nan–nan] | 0.67 [nan–nan] | 0.55 | 0.40 | 0.50/1.00 | 0.17/0.00 |
| FoldX | 8.88 [nan–nan] | 6.65 [nan–nan] | 0.05 | 0.20 | 0.50/1.00 | 0.17/0.00 |
| **U1A (mutation distance cutoff = 0.4 nm)** | | | | | | |
| IRIS | **0.22 [0.03–1.39]** | **0.17 [0.02–1.12]** | 0.04 | 0.16 | 0.62/0.59 | 0.58/0.27 |
| Rosetta-Vienna | 0.72 [0.07–1.36] | 0.53 [0.05–1.01] | 0.24 | 0.07 | 0.46/0.48 | 0.47/0.24 |
| Rosetta-Vienna (aug) | 0.41 [0.03–0.81] | 0.30 [0.03–0.66] | 0.48 | **0.41** | **0.70/0.60** | **0.66/0.36** |
| DeePNAP | 1.25 [0.72–1.79] | 0.93 [0.54–1.42] | -0.05 | 0.07 | 0.52/0.68 | 0.42/0.37 |
| FoldX | 3.34 [0.15–9.27] | 2.48 [0.11–7.94] | **0.70** | 0.14 | 0.61/0.49 | 0.53/0.22 |
| **U1A (mutation distance cutoff = 0.7 nm)** | | | | | | |
| IRIS | **0.25 [0.02–1.28]** | **0.19 [0.01–1.00]** | 0.04 | 0.16 | 0.59/0.63 | 0.53/0.28 |
| Rosetta-Vienna | 0.68 [0.05–1.27] | 0.51 [0.04–1.00] | 0.23 | 0.05 | 0.45/0.51 | 0.43/0.25 |
| Rosetta-Vienna (aug) | 0.43 [0.03–0.81] | 0.32 [0.02–0.65] | 0.46 | **0.35** | **0.65/0.52** | **0.69/0.33** |
| DeePNAP | 1.23 [0.71–1.74] | 0.93 [0.54–1.39] | -0.05 | 0.07 | 0.48/0.68 | 0.39/0.35 |
| FoldX | 3.68 [0.14–9.44] | 2.77 [0.10–8.15] | **0.70** | 0.14 | 0.59/0.46 | 0.55/0.21 |
| **U1A (mutation distance cutoff = 0.95 nm)** | | | | | | |
| IRIS | **0.29 [0.02–1.07]** | **0.22 [0.01–0.82]** | 0.09 | 0.26 | **0.59/0.80** | 0.54/0.42 |
| Rosetta-Vienna | 0.60 [0.05–1.14] | 0.45 [0.05–0.88] | 0.27 | 0.12 | 0.47/0.62 | 0.42/0.31 |
| Rosetta-Vienna (aug) | 0.45 [0.04–0.77] | 0.33 [0.03–0.59] | 0.48 | **0.30** | 0.59/0.42 | **0.76/0.41** |
| DeePNAP | 1.17 [0.71–1.63] | 0.87 [0.54–1.24] | -0.03 | 0.11 | 0.49/0.71 | 0.36/0.39 |
| FoldX | 4.13 [0.24–9.25] | 3.08 [0.16–7.91] | **0.69** | 0.12 | 0.54/0.41 | 0.69/0.36 |

Table S10: **Ablation analysis of IRIS on the MS2 dataset.** This analysis compares sequence-only, structure-exclusive, and fully integrated sequence-structure predictions (the augmented IRIS version). Sequence-only predictions were generated using sequence similarity metrics relative to the native sequence: Hamming distance (Seq-Ham) and 3-mer Jaccard distance (computed as 1− Jaccard to maintain consistent ordering as Hamming distance). For all IRIS and sequence-based predictions, error metrics (MAE, RMSE) were computed after fitting predicted vs. experimental values with a best-fit line constrained to pass through the origin. The structure-exclusive mode reports the mean ± standard deviation across 100 independent runs with randomized sequence inputs. Strong and weak binders are defined as the lowest and highest quartiles of experimental $\Delta\Delta G$ within each MS2–RNA group. Results are shown separately for MS2 RNA variants with 0–3 substitutions and those with 4 substitutions. Seq-Ham is not applicable to the 4-substitution group, as all variants share the same hamming distance.

| **MS2 (0–3 substitutions)** | | | | | | |
|---|---|---|---|---|---|---|
| **Method** | **RMSE** | **MAE** | **Pearson $r$** | **Spearman $\rho$** | **ROC-AUC (S/W)** | **PR-AUC (S/W)** |
| Seq-Ham | 0.54 [0.38–0.71] | 0.51 [0.36–0.67] | 0.21 | 0.13 | 0.65 / 0.32 | 0.25 / 0.12 |
| Seq-Jacc | 0.45 [0.31–0.60] | 0.43 [0.29–0.57] | 0.41 | 0.38 | 0.81 / 0.59 | 0.31 / 0.27 |
| Structure | 5.20 ± 3.85 | 4.95 ± 3.67 | 0.01 ± 0.17 | 0.01 ± 0.17 | 0.50 ± 0.09 / 0.51 ± 0.10 | 0.26 ± 0.05 / 0.31 ± 0.09 |
| IRIS (aug.) | 0.22 [0.05–0.37] | 0.21 [0.05–0.36] | 0.72 | 0.74 | 0.91 / 0.80 | 0.72 / 0.55 |
| **MS2 (4 substitutions)** | | | | | | |
| **Method** | **RMSE** | **MAE** | **Pearson $r$** | **Spearman $\rho$** | **ROC-AUC (S/W)** | **PR-AUC (S/W)** |
| Seq-Ham | 0.71 [0.48–0.91] | 0.67 [0.44–0.86] | NA | NA | NA | NA |
| Seq-Jacc | 1.08 [0.81–1.34] | 1.02 [0.77–1.27] | -0.08 | 0.16 | 0.51 / 0.64 | 0.26 / 0.28 |
| Structure | 5.35 ± 3.17 | 5.04 ± 2.98 | -0.01 ± 0.18 | -0.0057 ± 0.19 | 0.49 ± 0.11 / 0.50 ± 0.10 | 0.30 ± 0.08 / 0.32 ± 0.08 |
| IRIS (aug.) | 2.07 [1.49–2.71] | 1.94 [1.39–2.56] | 0.32 | 0.43 | 0.71 / 0.74 | 0.49 / 0.41 |

(S1) Alipanahi, B.; Delong, A.; Weirauch, M. T.; Frey, B. J. Predicting the sequence specificities of DNA- and RNA-binding proteins by deep learning. *Nat Biotechnol* **2015**, *33*, 831–838.

(S2) Kappel, K.; Jarmoskaite, I.; Vaidyanathan, P. P.; Greenleaf, W. J.; Herschlag, D.; Das, R. Blind tests of RNA-protein binding affinity prediction. *Proc Natl Acad Sci U S A* **2019**, *116*, 8336–8341.

(S3) Dias, R.; Kolazckowski, B. Different combinations of atomic interactions predict protein-small molecule and protein-DNA/RNA affinities with similar accuracy. *Proteins* **2015**, *83*, 2100–2114.

(S4) Leaver-Fay, A.; Tyka, M.; Lewis, S. M.; Lange, O. F.; Thompson, J.; Jacak, R.; Kaufman, K.; Renfrew, P. D.; Smith, C. A.; Sheffler, W.; Davis, I. W.; Cooper, S.; Treuille, A.; Mandell, D. J.; Richter, F.; Ban, Y.-E. A.; Fleishman, S. J.; Corn, J. E.; Kim, D. E.; Lyskov, S.; Berrondo, M.; Mentzer, S.; Popovi, Z.; Havranek, J. J.; Karanicolas, J.; Das, R.; Meiler, J.; Kortemme, T.; Gray, J. J.; Kuhlman, B.; Baker, D.; Bradley, P. ROSETTA3: an object-oriented software suite for the simulation and design of macromolecules. *Methods Enzymol* **2011**, *487*, 545–574.

(S5) Lorenz, R.; Bernhart, S. H.; Hner Zu Siederdissen, C.; Tafer, H.; Flamm, C.; Stadler, P. F.; Hofacker, I. L. ViennaRNA Package 2.0. *Algorithms Mol Biol* **2011**, *6*, 26.

(S6) Lyskov, S.; Chou, F.-C.; Conchir, S. .; Der, B. S.; Drew, K.; Kuroda, D.; Xu, J.; Weitzner, B. D.; Renfrew, P. D.; Sripakdeevong, P.; Borgo, B.; Havranek, J. J.; Kuhlman, B.; Kortemme, T.; Bonneau, R.; Gray, J. J.; Das, R. Serverification of Molecular Modeling Applications: The Rosetta Online Server That Includes Everyone (ROSIE). *PLoS ONE* **2013**, *8*, e63906.

(S7) Shen, X.; Hou, Y.; Wang, X.; Zhang, C.; Liu, J.; Shen, H.; Wang, W.; Yang, Y.;

Yang, M.; Li, Y.; Zhang, J.; Sun, Y.; Chen, K.; Shi, L.; Li, X. A deep learning model for characterizing protein-RNA interactions from sequences at single-base resolution. *Patterns* **2025**, *6*, 101150.

(S8) Pandey, U.; Behara, S. M.; Sharma, S.; Patil, R. S.; Nambiar, S.; Koner, D.; Bhukya, H. DeePNAP: A Deep Learning Method to Predict Protein-Nucleic Acid Binding Affinity from Their Sequences. *J Chem Inf Model* **2024**, *64*, 1806–1815.

(S9) Schymkowitz, J.; Borg, J.; Stricher, F.; Nys, R.; Rousseau, F.; Serrano, L. The FoldX web server: an online force field. *Nucleic Acids Res* **2005**, *33*, W382–388.

(S10) Delgado, J.; Radusky, L. G.; Cianferoni, D.; Serrano, L. FoldX 5.0: working with RNA, small molecules and a new graphical interface. *Bioinformatics* **2019**, *35*, 4168–4169.

(S11) Deng, L.; Yang, W.; Liu, H. PredPRBA: Prediction of Protein-RNA Binding Affinity Using Gradient Boosted Regression Trees. *Front Genet* **2019**, *10*, 637.

(S12) Lambert, N.; Robertson, A.; Jangi, M.; McGeary, S.; Sharp, P. A.; Burge, C. B. RNA Bind-n-Seq: quantitative assessment of the sequence and structural binding specificity of RNA binding proteins. *Mol Cell* **2014**, *54*, 887–900.

(S13) Dominguez, D.; Freese, P.; Alexis, M. S.; Su, A.; Hochman, M.; Palden, T.; Bazile, C.; Lambert, N. J.; Van Nostrand, E. L.; Pratt, G. A.; Yeo, G. W.; Graveley, B. R.; Burge, C. B. Sequence, Structure, and Context Preferences of Human RNA Binding Proteins. *Mol Cell* **2018**, *70*, 854–867.e9.

(S14) Wayment-Steele, H. K.; Kladwang, W.; Strom, A. I.; Lee, J.; Treuille, A.; Becka, A.; Eterna Participants; Das, R. RNA secondary structure packages evaluated and improved by high-throughput experiments. *Nat Methods* **2022**, *19*, 1234–1242.

(S15) Zhu, H.; Tang, F.; Quan, Q.; Chen, K.; Xiong, P.; Zhou, S. K. Deep generalizable prediction of RNA secondary structure via base pair motif energy. *Nat Commun* **2025**, *16*, 5856.

(S16) Pak, M. A.; Markhieva, K. A.; Novikova, M. S.; Petrov, D. S.; Vorobyev, I. S.; Maksimova, E. S.; Kondrashov, F. A.; Ivankov, D. N. Using AlphaFold to predict the impact of single mutations on protein stability and function. *PLoS One* **2023**, *18*, e0282689.
